# Systematic exploration of *Escherichia coli* phage-host interactions with the BASEL phage collection

**DOI:** 10.1101/2021.03.08.434280

**Authors:** Enea Maffei, Aisylu Shaidullina, Marco Burkolter, Valentin Druelle, Luc Willi, Fabienne Estermann, Sarah Michaelis, Hubert Hilbi, David S. Thaler, Alexander Harms

**Author notes:** These authors contributed equally (Enea Maffei, Aisylu Shaidullina). Correspondence should be addressed to Dr. Alexander Harms.

## Abstract

Bacteriophages, the viruses infecting bacteria, hold great potential for the treatment of multidrug-resistant bacterial infections and other applications due to their unparalleled diversity and recent breakthroughs in their genetic engineering. However, fundamental knowledge of molecular mechanisms underlying phage-host interactions is mostly confined to a few traditional model systems and did not keep pace with the recent massive expansion of the field. The true potential of molecular biology encoded by these viruses has therefore remained largely untapped, and phages for therapy or other applications are often still selected empirically. We therefore sought to promote a systematic exploration of phage-host interactions by composing a well-assorted library of 66 newly isolated phages infecting the model organism *Escherichia coli* that we share with the community as the BASEL collection (BActeriophage SElection for your Laboratory). This collection is largely representative of natural *E. coli* phage diversity and was intensively characterized phenotypically and genomically alongside ten well-studied traditional model phages. We experimentally determined essential host receptors of all phages, quantified their sensitivity to eleven defense systems across different layers of bacterial immunity, and matched these results to the phages’ host range across a panel of pathogenic enterobacterial strains. Our results reveal clear patterns in the distribution of phage phenotypes and genomic features that highlight systematic differences in the potency of different immunity systems and point towards the molecular basis of receptor specificity in several phage groups. Strong trade-offs were detected between fitness traits like broad host recognition and resistance to bacterial immunity that might drive the divergent adaptation of different phage groups to specific niches. We envision that the BASEL collection will inspire future work exploring the biology of bacteriophages and their hosts by facilitating the discovery of underlying molecular mechanisms as the basis for an effective translation into biotechnology or therapeutic applications.

## Introduction

Bacteriophages, the viruses infecting bacteria, are the most abundant biological entities on earth with key positions in all ecosystems and carry large part of our planet’s genetic diversity in their genomes [1–3]. Out of this diversity, a few phages infecting *Escherichia coli* became classical models of molecular biology with roles in many fundamental discoveries and are still major workhorses of research today [4]. The most prominent of these are the seven “T phages” T1 – T7 [5, reviewed in reference 6] and bacteriophage lambda [7]. Like the majority of known phages, these classical models are tailed phages or *Caudovirales* that use characteristic tail structures to bind host surface receptors and to inject their genomes from the virion head into the host cell. Three major virion morphotypes of *Caudovirales* are known, myoviruses with a contractile tail, siphoviruses with a long and flexible tail, and podoviruses with a very short, stubby tail [2] (Fig 1A). While the T phages are all so-called lytic phages and kill their host to replicate at each infection event, lambda is a temperate phage and can either kill the host to directly replicate or decide to integrate into the host’s genome as a prophage for transient passive replication by vertical transmission in the so-called lysogen [2, 8]. These two alternative lifestyles as lytic or temperate phages have major implications for viral ecology and evolution: While lytic phages have primarily been selected to overcome host defenses and maximize virus replication, temperate phages characteristically encode genes that increase the lysogens’ fitness, e.g., by providing additional bacterial immunity systems to fight other phages [8, 9].

**Fig 1.**
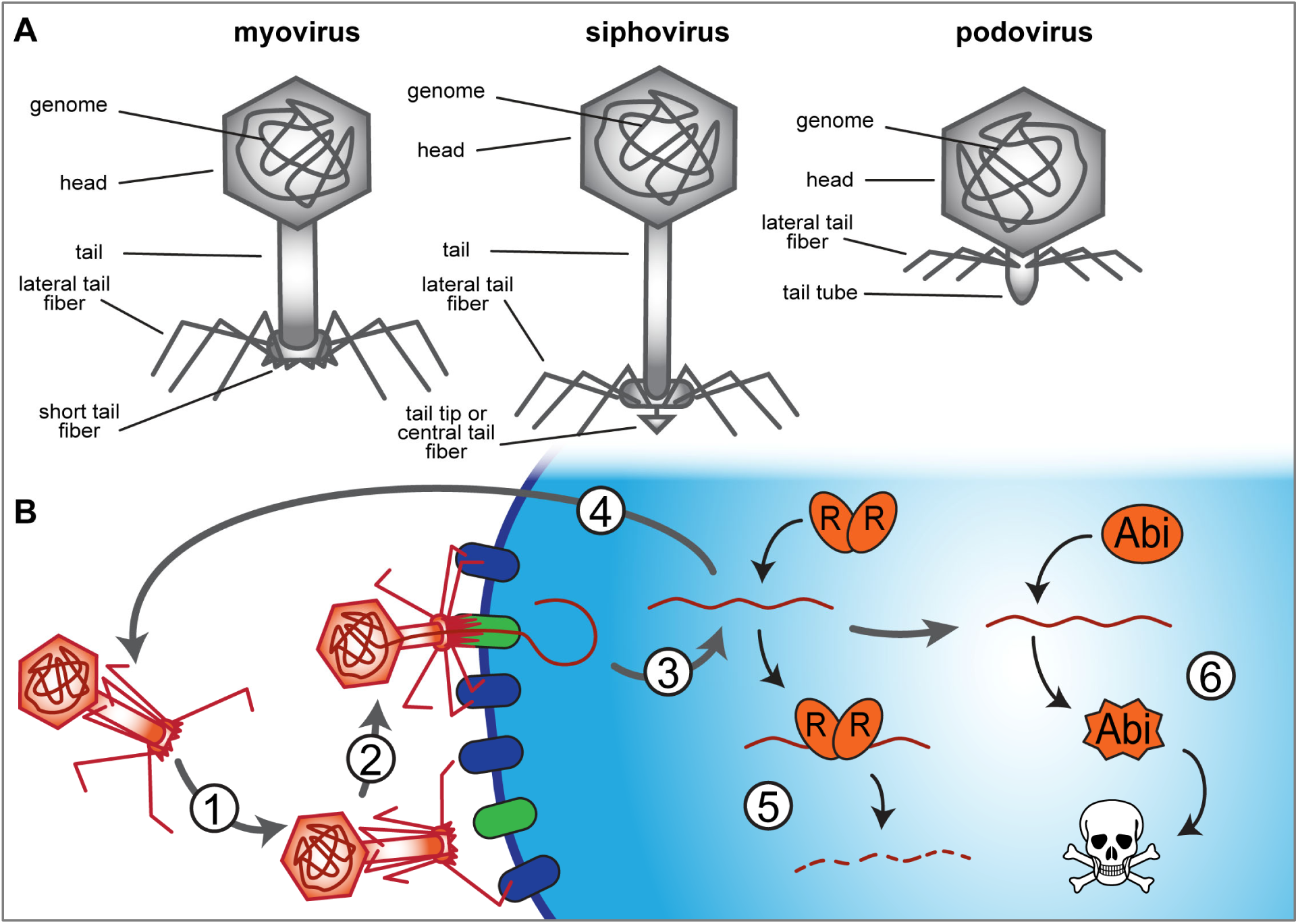
The three morphotypes of *Caudovirales* and two lines of defense in bacterial immunity. **(A)** The virions of tailed phages or *Caudovirales* can be assigned to three general morphotypes including myoviruses (contractile tail), siphoviruses (long and flexible, non-contractile tail), and podoviruses (short and stubby tail). **(B)** The life cycle of a typical lytic phage begins with reversible attachment to a so-called primary receptor on the bacterial cell surface (1), usually via lateral tail fibers at the virion. Subsequently, irreversible attachment to secondary or terminal receptors usually depends on structures at the end of the tail, e.g., short tail fibers for many myoviruses and central tail fibers or tail tip proteins for siphoviruses (2; see also (A)). After genome injection (3), the phage takes over the host cell, replicates, and releases the offspring by host cell lysis (4). Inside the host cell, the bacteriophage faces two lines of host defenses, first bacterial immunity systems that try to clear the infection by directly targeting the phage genome (5) and then abortive infection systems that kill the infected cell when a viral infection is sensed (6).

The ubiquity of phage predation has driven the evolution of a vast arsenal of bacterial immunity systems targeting any step of phage infection [9, 10]. Inside host cells, phages encounter two lines of defense of which the first primarily comprises restriction-modification (RM) systems or CRISPR-Cas, the bacterial adaptive immunity, that directly attack viral genomes [11] (Fig 1B). A second line of defense is formed by diverse abortive infection (Abi) systems that protect the host population by triggering an altruistic suicide of infected cells when sensing viral infections [9–12] (Fig 1B). While RM systems and CRISPR-Cas are highly abundant and have been successfully adapted for biotechnology (honored with Nobel Prizes in 1978 and 2020, respectively), the molecular mechanisms underlying the function of collectively abundant, but each individually rare, Abi systems have remained elusive with few exceptions [9–12].

Research on phages has expanded at breathless pace over the last decade with a focus on biotechnology and on clinical applications against bacterial infections (“phage therapy”) [13, 14]. Besides or instead of the few traditional model phages, many researchers now employ comparably poorly described, newly isolated phages that are often only used for a few studies and available only in their laboratory. The consequence of this development is a rapidly growing amount of very patchy data such as, e.g., the currently more than 14’000 available unique phage genomes [15] for which largely no linked phenotypic data are available. Despite the value of proof-of-principle studies and a rich genome database, this lack of systematic, interlinked data in combination with the diversity of bacteriophages makes it very difficult to gain a mechanistic understanding of phage biology or to uncover patterns in the data that would support the discovery of broad biological principles beyond individual models.

As an example, phage isolates for treating a specific case of bacterial infection are necessarily chosen largely empirically due to the lack of systematic data about relevant phage properties. Currently, the selection of native phages and their engineering for therapeutic applications primarily focus on a lytic lifestyle, a broad host range, and very occasionally on biofilm- or cell wall-degrading enzymes that are comparably well understood genetically and mechanistically [13, 14, 16, 17]. However, the molecular mechanisms and genetic basis underlying other desired features such as resistance to different bacterial immunity systems and, in general, the distribution of all these features across different groups of phages have remained understudied. Given the notable incidence of treatment failure in phage therapy [18–20], a better understanding of the links between phage taxonomy, genome sequence, and phenotypic properties seems timely to select more effective native phages for therapeutic applications and to expand the potential of phage engineering.

In this work we therefore present the BASEL (<u>Ba</u>cteriophage <u>Se</u>lection for your <u>L</u>aboratory) collection as a reference set of 66 newly isolated lytic bacteriophages that infect the laboratory strain *E. coli* K-12 and make it accessible to the scientific community. We provide a systematic phenotypic and genomic characterization of these phages alongside ten classical model phages regarding host receptors, sensitivity and resistance to bacterial immunity, and host range across diverse enterobacteria. Our results highlight clear phenotypic patterns between and within taxonomic groups of phages that reveal strong trade-offs between important bacteriophage traits. These findings greatly expand our understanding of bacteriophage ecology, evolution, and their interplay with bacterial immunity systems. We therefore anticipate that our work will not only establish the BASEL collection as a reference point for future studies exploring fundamental bacteriophage biology but also promote a rational application of phage therapy based on an improved selection and engineering of bacteriophages.

## Results

### Composition of the BASEL collection

The first aim of our study was to generate a collection of new phage isolates infecting the ubiquitously used *E. coli* K-12 laboratory strain that would provide representative insight into the diversity of tailed, lytic phages by covering all major groups and containing a suitable selection of minor ones. Similar collections exist, e.g., for *E. coli* in form of the ECOR collection of 72 *E. coli* strains that are commonly used to study how this model organism varies in certain traits or can deal with / evolves under certain conditions [21, 22]. Though nothing truly comparable is available for phages, it is notable that the seven T phages had originally been chosen in the early 1940s with the explicit aim of providing a reference point that would enable comparative, systematic research on bacteriophages [reviewed in reference 6]. However, while these and the other classical model phages such as lambda have been invaluable to uncover many fundamental principles of molecular biology, their number and taxonomic range are too limited to serve as a representative reference even for bacteriophages infecting *E. coli*.

As a first step, we generated a derivative of *E. coli* K-12 without any of the native barriers that might limit or bias phage isolation unfavorably. This strain, *E. coli* K-12 MG1655 ΔRM, therefore lacks the O-antigen glycan barrier (see below), all restriction systems, as well as the RexAB and PifA Abi systems (see *Materials and Methods* as well as S1 Text). *E. coli* K-12 ΔRM was subsequently used to isolate hundreds of phages from environmental samples such as river water or compost, but mostly from the inflow of different sewage treatment facilities (Fig 2A and *Materials and Methods*).

**Fig 2.**
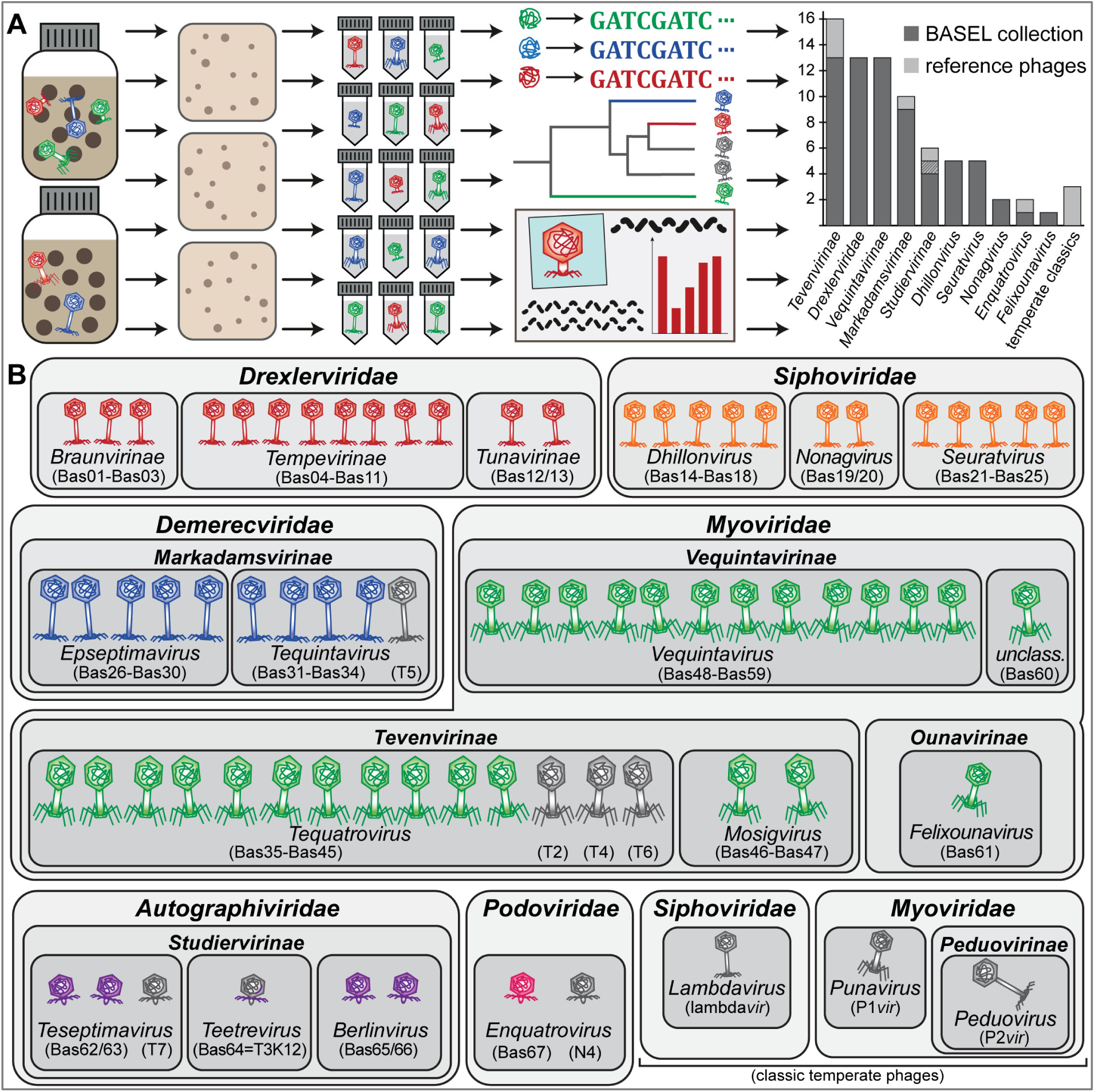
Overview of the BASEL collection. **(A)** Illustration of the workflow of bacteriophage isolation, characterization, and selection that resulted in the BASEL collection (details in *Materials and Methods*). **(B)** Taxonomic overview of the bacteriophages included in the BASEL collection and their unique Bas## identifiers. Newly isolated phages are colored according to their family while well-studied reference phages are shown in grey.

Previous bacteriophage isolation studies had already provided deep insight into the diversity of tailed, lytic *E. coli* phages in samples ranging from sewage over diverse natural environments to infant guts or blood and urine of patients in tertiary care [23–29]. Despite the wide diversity of known *E. coli* phages [30], nearly all phage isolates reported in these studies belonged to five major groups and were either myoviruses of 1) *Tevenvirinae* or 2) *Vequintavirinae* subfamilies and close relatives, 3) large siphoviruses of the *Markadamsvirinae* subfamily within *Demerecviridae*, or small siphoviruses of 4) diverse *Drexlerviridae* subfamilies or 5) the genera *Dhillonvirus*, *Nonagvirus*, and *Seuratvirus* of the *Siphoviridae* family. Podoviruses of any kind were rarely reported, and if, then were mostly *Autographiviridae* isolated using enrichment cultures that are known to greatly favor such fast-growing phages [24, 31]. This pattern does not seem to be strongly biased by any given strain of *E.coli* as isolation host because a large, very thorough study using diverse *E. coli* strains reported essentially the same composition of taxonomic groups [23].

Whole-genome sequencing of around 120 different isolates from our isolation experiments largely reproduced this pattern, which suggests that several intrinsic limitations of our approach did not strongly affect the spectrum of phages that we sampled (S3 Text). After eliminating closely related isolates, the BASEL collection was formed as a set of 66 new phage isolates (Fig 2; see also *Materials and Methods* and S5 Table). We deliberately did not give full proportional weight to highly abundant groups like *Tevenvirinae* so that the BASEL collection is not truly representative in a narrow quantitative sense. Instead, we included as many representatives of rarely isolated groups (e.g., podoviruses) as possible to increase the biological diversity of phages in our collection that could be an important asset for studying the genetics of phage / host interactions or for unraveling genotype / phenotype relationships. Besides these 66 new isolates (serially numbered as Bas01, Bas02, etc.; Fig 2 and S5 Table), we included a panel of ten classical model phages in our genomic and phenotypic characterization and view them as an accessory part of the BASEL collection. Beyond the T phages (without T1 that is a notorious laboratory contaminant [32]), we included well-studied podovirus N4 and obligately lytic mutants of the three most commonly studied temperate phages lambda, P1, and P2 [5, 7, 33, 34] (Fig 2; see also S5 Table).

### Overview: Identification of phage surface receptors

The infection cycle of most tailed phages begins with host recognition by reversible adsorption to a first “primary” receptor on host cells (often a sugar motif on surface glycans) followed by irreversible binding to a terminal or “secondary” receptor directly on the cell surface which results in DNA injection (Fig 1B) [35]. Importantly, the adsorption to many potential hosts is blocked by long O-antigen chains on the LPS and other exopolysaccharides that effectively shield the cell surface unless they can be degraded or specifically serve as the phage’s primary receptor [35–38]. Tailed phages bind to primary and secondary receptors using separate structures at the phage tail which display the dedicated receptor-binding proteins (RBPs) in the form of tail fibers, tail spikes (smaller than fibers, often with enzymatic domains), or central tail tips [comprehensively reviewed in reference 35]. Most commonly, surface glycans such as the highly variable O-antigen chains or the enterobacterial common antigen (ECA) are bound as primary receptors. Secondary receptors on Gram-negative hosts are near-exclusively porin-family outer membrane proteins for siphoviruses and core LPS sugar structures for podoviruses, while myoviruses were found to use either one of the two depending on phage subfamily or genus [35, 39, 40]. Notably, no host receptor is known for the vast majority of phages that have been studied, but understanding the genetic basis and molecular mechanisms underlying host recognition as the major determinant of phage host range is a crucial prerequisite for host range engineering or a rational application of phage therapy [41]. Since tail fibers and other host recognition modules are easily identified in phage genomes, we used phage receptor specificity as a model to demonstrate the usefulness of systematic phenotypic data with the BASEL collection as a key to unlock information hidden in the genome databases.

We therefore first experimentally determined the essential host receptor(s) of all phages in the BASEL collection before analyzing the phages’ genomes for the mechanisms underlying receptor specificity. Briefly, the dependence on surface proteins was assessed by plating each phage on a set of more than fifty single-gene mutants (S1 Table) or whole-genome sequencing of spontaneously resistant bacterial clones (see *Materials and Methods*). The role of host surface glycans was quantified with *waaG* and *waaC* mutants that display different truncations of the LPS core, a *wbbL(+)* strain with restored O16-type O-antigen expression, and a *wecB* mutant that is specifically deficient in production of the ECA [42–44] (Fig 3A, S1 Table, and *Materials and Methods*).

**Fig 3.**
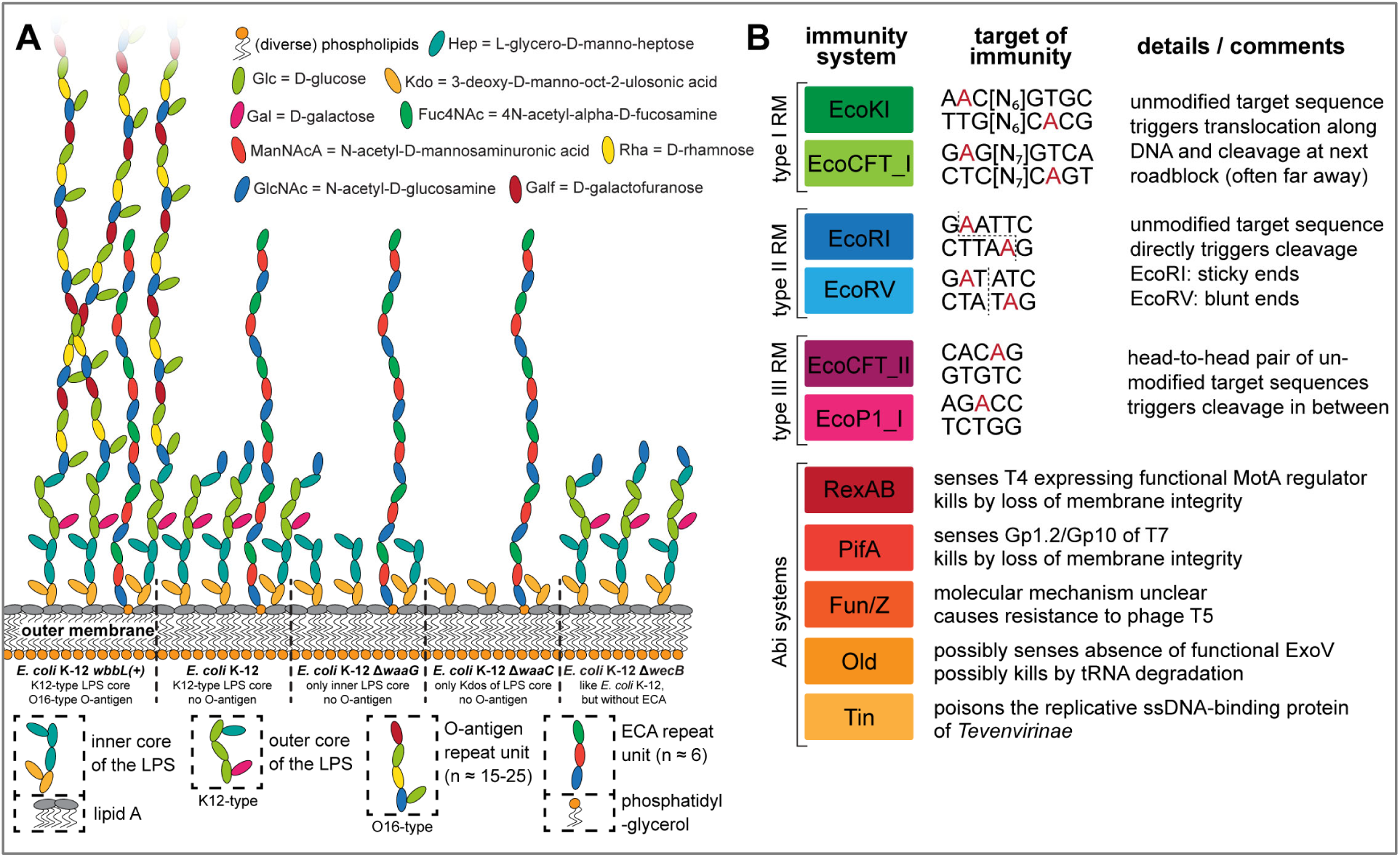
Overview of *E. coli* surface glycan variants and the immunity systems used in this study. **(A)** The surface glycans of different *E. coli* K-12 MG1655 variants are shown schematically (details in running text and *Materials and Methods*). Note that the *E. coli* K-12 MG1655 laboratory wildtype does not merely display the K12-type core LPS (classical rough LPS phenotype) but also the most proximal glucose of the O16-type O-antigen. **(B)** Key features of the six RM systems (each two of type I, type II, and type III) and the five Abi systems used for the phenotyping of this study are summarized schematically. Recognition sites of RM systems have either been determined experimentally or were predicted in REBASE (red nucleotides: methylation sites; dotted lines: cleavage sites) [47–49, 128]. The Abi systems have been characterized to very different extent but constitute the most well-understood representatives of these immunity systems of *E. coli* [10, 50].

### Overview: Phenotyping of sensitivity / resistance to bacterial immunity systems

Some bacterial immunity systems are very common among different strains of a species (like certain types of RM systems for *E. coli*), while others – especially the various Abi systems – have each a very patchy distribution but are no less abundant if viewed together [11]. Each bacterial strain therefore encodes a unique repertoire of a few very common and a larger number of rarer, strain-specific immunity systems, but it is unknown how far these systems impact the isolation of phages or their efficacy in therapeutic applications. A systematic view of the potency and target range of different immunity systems and how the diverse groups of phages differ in sensitivity / resistance to these systems might enable us to select or engineer phages with a higher and more reliable potency for phage therapy or biotechnology. As an example, previous work showed that *Tevenvirinae* or *Seuratvirus* and *Nonagvirus* phages exhibit broad resistance to RM systems due to the hypermodification of cytosines or guanosines in their genomes, respectively, which could be an interesting target for phage engineering [45, 46]. However, it is unknown whether this mechanism of RM resistance (or any other viral anti-immunity function) actually results in a measurably broader phage host range.

We therefore systematically quantified the sensitivity of all phages of the BASEL collection against a panel of eleven immunity systems and scored their infectivity on a range of pathogenic enterobacteria that are commonly used as model systems (see *Materials and Methods* and Fig 3B). Shortly, we tested six RM systems by including each two type I, type II, and type III systems that differ in the molecular mechanisms of DNA modification and cleavage [10, 47–49] (Fig 3B). Besides these, we included the most well-studied Abi systems of *E. coli*, RexAB of the lambda prophage, PifA of the F-plasmid, as well as the Old, Tin, and Fun/Z systems of the P2 prophage (Fig 3B). Previous work suggested that RexAB and PifA sense certain proteins of phages T4 and T7, respectively, to trigger host cell death by membrane depolarization [10]. Conversely, Old possibly senses the inhibition of RecBCD / ExoV during lambda infections and might kill by tRNA degradation, Fun/Z abolishes infections by phage T5 via an unknown mechanism, and Tin poisons the replicative ssDNA binding protein of *Tevenvirinae* [50]. Beyond *E. coli* K-12, we quantified the infectivity of the BASEL collection on uropathogenic *E. coli* (UPEC) strains UTI89 and CFT073, enteroaggregative *E. coli* (EAEC) strain 55989, alternative laboratory strain *E. coli* B REL606, and *Salmonella enterica* subsp. *enterica* serovar Typhimurium strains 12023s and SL1344 (see *Materials and Methods* and S1 Table).

### Properties of the *Drexlerviridae* family

Phages of the *Drexlerviridae* family (previously also known as T1 superfamily [30]) are small siphoviruses with genome sizes of ca. 43-52 kb (Figs 4A-D). The BASEL collection contains thirteen new *Drexlerviridae* phages that are broadly spread out across the various subfamilies and genera of this family (Fig 4B). Though it has apparently never been directly demonstrated, it seems likely that the *Drexlerviridae* use their lateral tail fibers in the same way as their larger cousins of the *Markadamsvirinae* (see below) to contact specific O-antigen glycans as primary receptors without depending on this interaction for host recognition [51] (Fig 4A). Consistently, the locus encoding the lateral tail fibers is very diverse among *Drexlerviridae* and the proteins forming these tail fibers are only highly related at the far N-terminus of the distal subunits where these are likely attached to the tail (S1B Fig). Among the *Drexlerviridae* that we isolated, only JakobBernoulli (Bas07) shows robust plaque formation on *E. coli* K-12 MG1655 with restored O16-type O-antigen expression, suggesting that its lateral tail fibers can use this O-antigen as primary receptor (Figs 4A and 4D).

**Fig 4.**
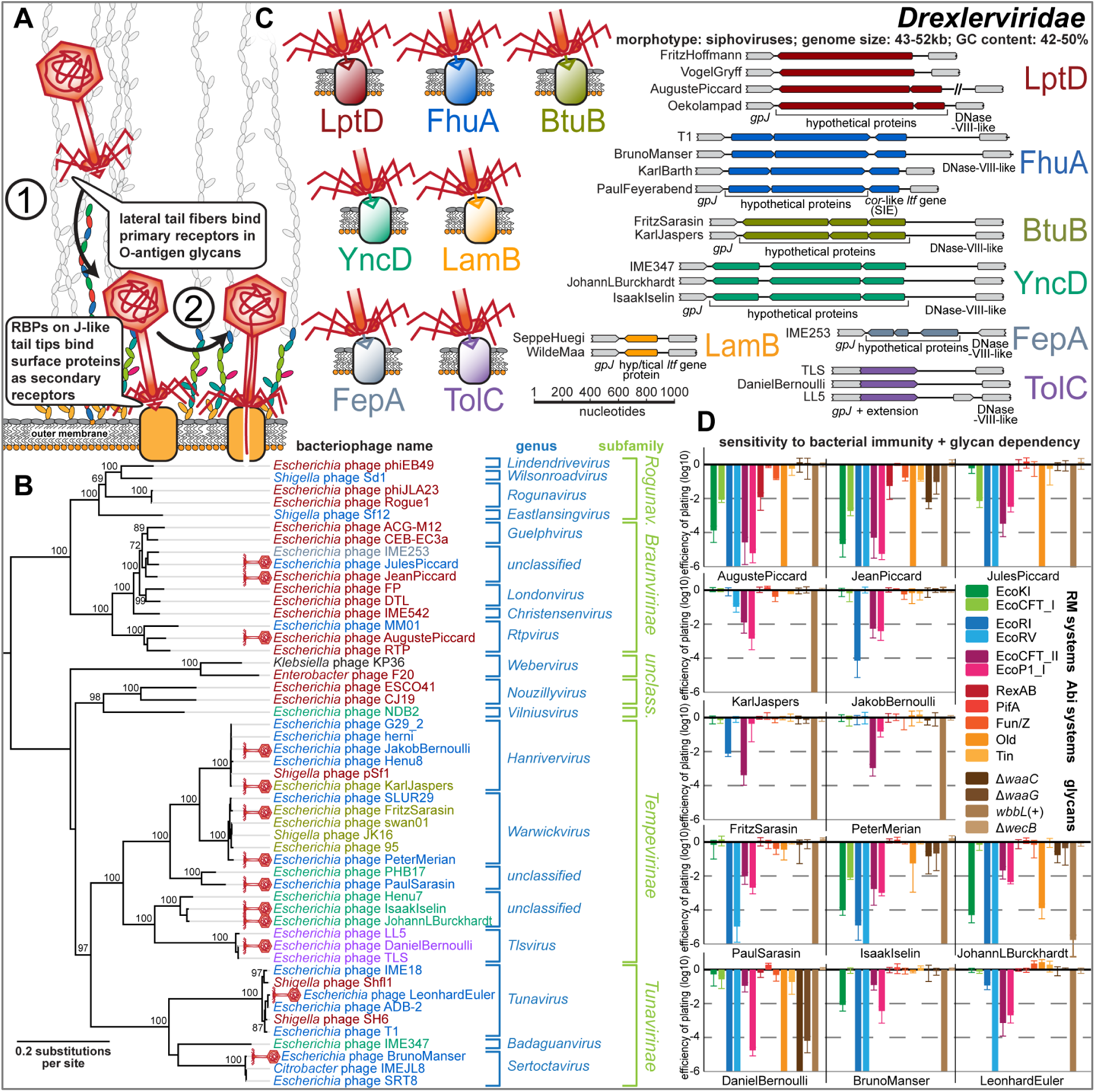
Overview of *Drexlerviridae* phages. **(A)** Schematic illustration of host recognition by *Drexlerviridae*. **(B)** Maximum-Likelihood phylogeny of *Drexlerviridae* based on several core genes with bootstrap support of branches shown if > 70/100. Newly isolated phages of the BASEL collection are highlighted by red phage icons and the determined or proposed terminal receptor specificity is highlighted at the phage names using the color code highlighted in (C). The phylogeny was rooted based on a representative phylogeny including *Dhillonvirus* sequences as outgroup (S1A Fig). **(C)** On the left, the seven identified receptors of small siphoviruses are shown with a color code that is also used to annotate demonstrated or predicted receptor specificity in the phylogenies of Fig 4B and Figs 6A + 6C). On the right, we show representative *bona fide* RBP loci that seem to encode the receptor specificity of these small siphoviruses (with the same color code). Note that the loci linked to each receptor are very similar while the genetic arrangement differs considerably between loci linked to different terminal host receptors (see also S1C Fig). **(D)** The results of quantitative phenotyping experiments with *Drexlerviridae* phages regarding sensitivity to altered surface glycans and bacterial immunity systems are presented as efficiency of plating (EOP). Data points and error bars represent average and standard deviation of at least three independent experiments.

Just like the much larger *Markadamsvirinae* siphoviruses (see below), *Drexlerviridae* phages use a small set of outer membrane porins as their secondary / final receptors for irreversible adsorption and DNA injection. To the best of our knowledge, previous work had only identified the receptors of T1 and IME18 (FhuA), IME347 (YncD), IME253 (FepA), and of LL5 as well as TLS (TolC) while for an additional phage, RTP, no protein receptor could be identified [40, 52–54]. Using available single-gene mutants, we readily determined the terminal receptor of eleven out of our thirteen *Drexlerviridae* phages as FhuA, BtuB, YncD, and TolC (Figs 4B and 4C) but failed for two others, AugustePiccard (Bas01) and JeanPiccard (Bas02). However, whole-genome sequencing of spontaneously resistant *E. coli* mutants showed that resistance was linked to mutations in the gene coding for LptD, the LPS export channel [36], strongly suggesting that this protein was the terminal receptor of these phages (Fig 5 and *Materials and Methods*).

**Fig 5.**
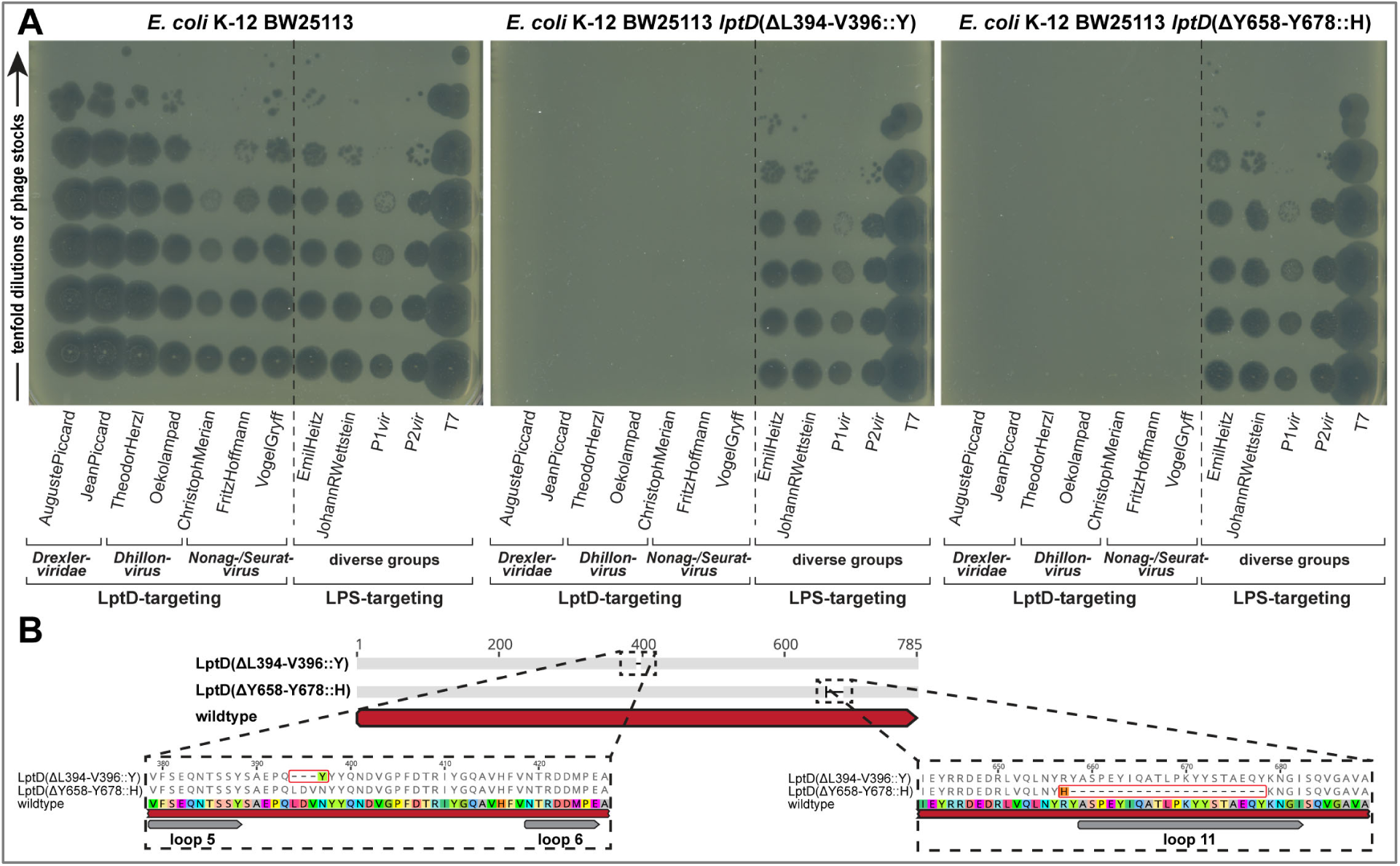
LptD is a commonly targeted terminal receptor of small siphoviruses. **(A)** Whole-genome sequencing of bacterial mutants exhibiting spontaneous resistance to seven small siphoviruses with no previously known receptor revealed different mutations or small deletions in the essential gene *lptD* that encodes the LptD LPS export channel. Top agar assays with two representative mutants in comparison to the ancestral *E. coli* K-12 BW25113 strain were performed with serial tenfold dilutions of twelve different phages (undiluted high-titer stocks at the bottom and increasingly diluted samples towards the top). Both mutants display complete resistance to the seven small siphoviruses of diverse genera within *Drexlerviridae* and *Siphoviridae* families that share the same *bona fide* RBP modules (S1C Fig) while no other phage of the BASEL collection was affected. In particular, we excluded indirect effects, e.g., via changes in the LPS composition in the *lptD* mutants, by confirming that five LPS-targeting phages of diverse families (see below) showed full infectivity on all strains. **(B)** The amino acid sequence alignment of wildtype LptD with the two mutants highlighted in (A) shows that resistance to LptD-targeting phages is linked to small deletions in or adjacent to regions encoding extracellular loops as defined in previous work [147], suggesting that they abolish the RBP-receptor interaction.

Similar to the central tail fibers of *Markadamsvirinae*, the RBPs of *Drexlerviridae* are thought to be displayed at the distal end of a tail tip protein related to well-studied GpJ of bacteriophage lambda [51, 54, 55]. The details of this host recognition module had remained elusive, though it was suggested that (like for T5 and unlike for lambda) dedicated RBPs are non-covalently attached to the J-like protein and might be encoded directly downstream of the *gpJ* homologs together with cognate superinfection exclusion proteins [54, 55]. By comparing the genomes of all *Drexlerviridae* phages with experimentally determined surface protein receptors, we were able to match specific allelic variants of this *bona fide* RBP loci to each known surface receptor of this phage family (Fig 4C). Notably, for the three known phages targeting TolC including DanielBernoulli (Bas08), the RBP locus is absent and apparently functionally replaced by a C-terminal extension of the GpJ-like tail tip protein that probably directly mediates receptor specificity like GpJ of bacteriophage lambda (Fig 4C) [56, 57]. Interestingly, similar RBP loci with homologous alleles are also found at the same genomic locus in small *Siphoviridae* of *Dhillonvirus*, *Nonagvirus*, and *Seuratvirus* genera (see below) where they also match known receptor specificity without exception (Figs 4-6 and S1C). Pending experimental validation, we therefore conclude that these distantly related groups of small siphoviruses share a common, limited repertoire of RBPs that enables receptor specificity of most representatives to be predicted *in silico*. As an example, it seems clear that phage RTP targets LptD as terminal receptor (Figs 4B and S1D). The poor correlation of predicted receptor specificity with the phylogenetic relationships of small siphoviruses (Figs 4B, 6B, and 6D) is indicative of frequent horizontal transfer of these RBP loci. This highlights the modular nature of this host recognition system which might enable the targeted engineering of receptor specificity to generate “designer phages” for different applications as previously shown for siphoviruses infecting *Listeria* [14, 58]. LptD would be a particularly attractive target for such engineering because it is strictly essential under all conditions, highly conserved, and heavily constrained due to multiple interactions that are critical for its functionality [36].

**Fig 6.**
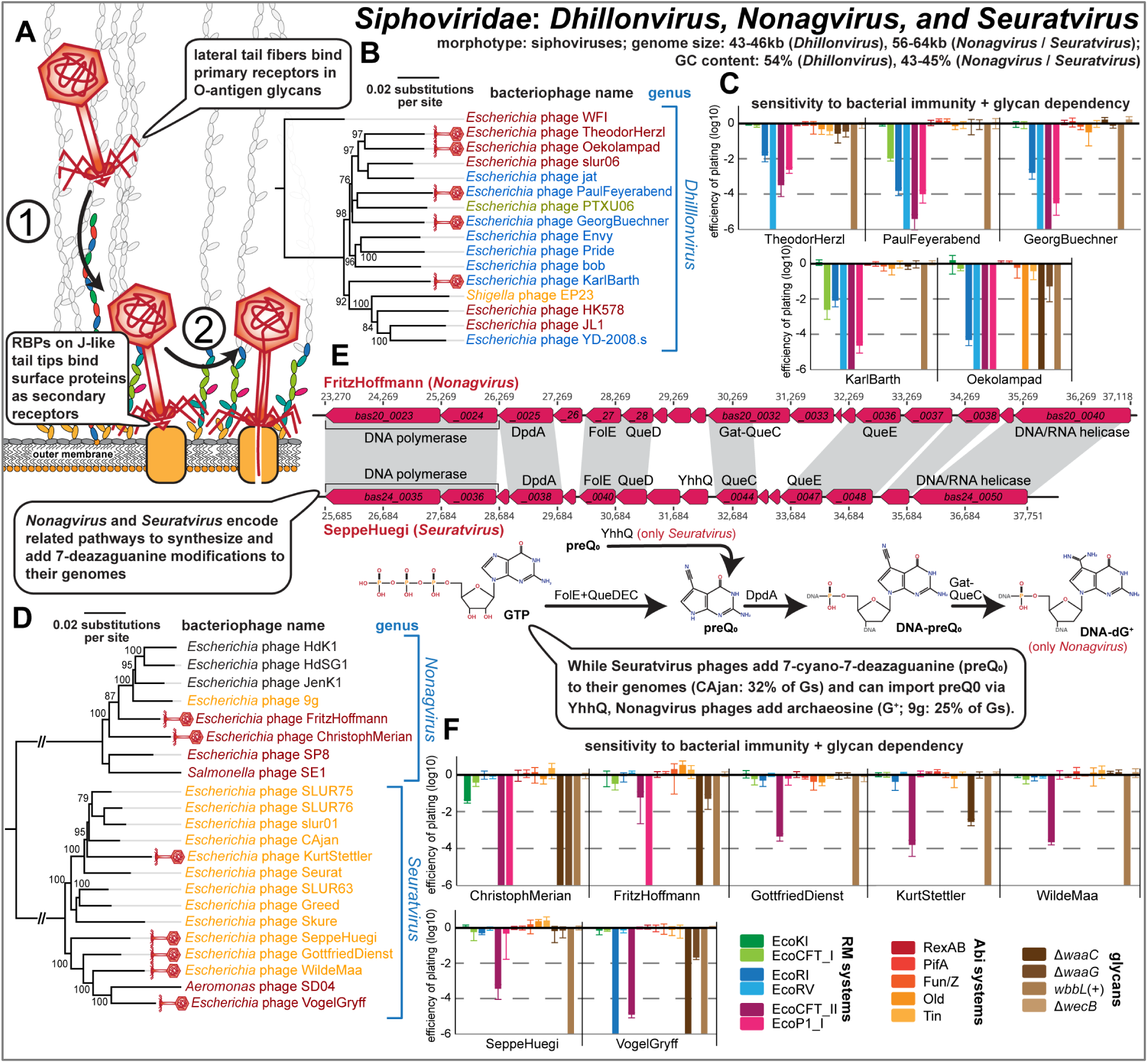
Overview of Siphoviridae genera Dhillonvirus, Nonagvirus, and Seuratvirus. **(A)** Schematic illustration of host recognition by small siphoviruses. **(B)** Maximum-Likelihood phylogeny of the *Dhillonvirus* genus based on a whole-genome alignment with bootstrap support of branches shown if > 70/100. Newly isolated phages of the BASEL collection are highlighted by red phage icons and the determined or proposed terminal receptor specificity is highlighted at the phage names using the color code highlighted in Fig 4C. The phylogeny was rooted between phage WFI and all others based on a representative phylogeny including *Drexlerviridae* sequences as outgroup (S1A Fig). **(C)** The results of quantitative phenotyping experiments with *Dhillonvirus* phages regarding sensitivity to altered surface glycans and bacterial immunity systems are presented as efficiency of plating (EOP). **(D)** Maximum-Likelihood phylogeny of the *Nonagvirus* and *Seuratvirus* genera based on a whole-genome alignment with bootstrap support of branches shown if > 70/100. Newly isolated phages of the BASEL collection are highlighted by red phage icons and the determined or proposed terminal receptor specificity is highlighted at the phage names using the color code highlighted in Fig 4C. The phylogeny was rooted between the two genera. **(E)** *Nonagvirus* and *Seuratvirus* phages share a core 7-deazaguanosine biosynthesis pathway involving FolE, QueD, QueE, and QueC which synthesizes dPreQ_0_ that is inserted into their genomes by DpdA. In *Nonagvirus* phages, the fusion of QueC with a glutamate amidotransferase (Gat) domain to Gat-QueC results in the modification with dG^+^ instead of dPreQ_0_ [45]. **(F)** The results of quantitative phenotyping experiments with *Nonagvirus* and *Seuratvirus* phages regarding sensitivity to altered surface glycans and bacterial immunity systems are presented as efficiency of plating (EOP). In (C) and (F), data points and error bars represent average and standard deviation of at least three independent experiments.

Once inside the host cell, our results show that *Drexlerviridae* phages are highly diverse in their sensitivity or resistance to diverse bacterial immunity systems (Fig 4D). While few representatives like FritzSarasin (Bas04) or PeterMerian (Bas05) are highly resistant, most *Drexlerviridae* are very sensitive to any kind of RM systems sometimes including the type I machineries that are largely unable to target any other phage that we tested (Fig 4D). Previous work suggested that *Drexlerviridae* might employ DNA methyltransferases as a defense strategy against host restriction [24], and these phages indeed encode variable sets of N6-adenine and C5-cytosine methyltransferases. However, we found no obvious pattern that would link DNA methyltransferases or any other specific genomic features to restriction resistance / sensitivity. Instead, sensitivity and resistance to bacterial immunity strongly correlate with *Drexlerviridae* phylogeny: While *Hanrivervirus* and *Warwickvirus* genera of *Tempevirinae* are highly resistant and *Tunavirinae* show comparably intermediate sensitivity, the other phages (in particular *Braunvirinae*, but also *Tlsvirus* phage DanielBernoulli (Bas08)) are highly sensitive (Figs 4B and 4D).

### Properties of Siphoviridae genera Dhillonvirus, Nonagvirus, and Seuratvirus

Phages of the *Dhillonvirus*, *Seuratvirus*, and *Nonagvirus* genera within the *Siphoviridae* family are small siphoviruses that are superficially similar to *Drexlerviridae* and have genomes with characteristic size ranges of 43-46 kb (*Dhillonvirus*), 56-61 kb (*Seuratvirus*), and 56-64 kb (*Nonagvirus*; see Fig 6A-E). Our twelve isolates included in the BASEL collection are spread out broadly across the phylogenetic ranges of these genera (Figs 6B and 6D). Similar to *Drexlerviridae* (and *Markadamsvirinae*, see below), we suggest that they also recognize glycan motifs at the O-antigen as their primary receptor on different host strains (Fig 6A). This notion is strongly supported by the remarkable variation exhibited by the different genomes at the lateral tail fiber locus (S2A Fig), as observed previously [23, 24]. None of these phages can infect *E. coli* K-12 MG1655 with restored O16-type O-antigen expression, but (like for *Drexlerviridae*) some require an intact LPS core for infectivity (Figs 6C and 6F). Experimental identification of the terminal receptor of all small *Siphoviridae* confirmed that receptor specificity seems to be encoded by the same system of *bona fide* RBP loci downstream of *gpJ* as for the *Drexlerviridae* (Figs 4B, 6B, and 6D). Many of these phages target LptD or FhuA as terminal receptors while others, unlike any *Drexlerviridae*, bind to LamB (Figs 6C and 6F). Notably, three related *Nonagvirus* phages encode a distinct *bona fide* RBP module that could not be matched to any known terminal receptor (S2B Fig), suggesting that these phages might target a different protein.

In difference to *Dhillonvirus* phages that are relatives of *Drexlerviridae* [23] (see also S1A Fig), *Nonagvirus* and *Seuratvirus* phages are closely related genera with genomes that are 10-15 kb larger than those of the other small siphoviruses in the BASEL collection (S5 table) [59]. This difference in genome size is largely due to genes encoding their signature feature, the 7-deazaguanine modification of 2′-deoxyguanosine (dG) in their genomes into 2′-deoxy-7-cyano-7-deazaguanosine (dpreQ_0_) for *Seuratvirus* phages and into 2′-deoxyarchaosine (dG^+^) for *Nonagvirus* phages [45] (Fig 6E). These modifications were shown to provide considerable (dG^+^) or at least moderate (dpreQ_0_) protection against restriction of genomic DNA of *Seuratvirus* phage CAjan and *Nonagvirus* phage 9g *in vitro*, although chemical analyses showed that only around one third (CAjan) or one fourth (9g) of the genomic dG content is modified [45] (Fig 6E). Consistently, we found that both *Nonagvirus* and *Seuratvirus* phages are remarkably resistant to type I and type II RM systems in our infection experiments, particularly if compared to other small siphoviruses of *Dhillonvirus* or *Drexlerviridae* groups (Figs 4D, 6C, and 6F). Among *Nonagvirus* and *Seuratvirus* phages, only ChristophMerian (Bas19, EcoKI) and VogelGryff (Bas25, EcoRI) show some sensitivity to these RM systems. However, all phages of these genera are highly sensitive to type III RM systems (Fig 6F). This observation does not necessarily indicate a lower sensitivity of type III RM systems to guanosine modifications *per se* but is likely simply a consequence of the much larger number of type III RM recognition sites in their genomes (S5 Table, see also Fig 3B).

### Properties of *Demerecviridae: Markadamsvirinae*

Phages of the *Markadamsvirinae* subfamily of the *Demerecviridae* family (previously known as T5-like phages [30]) are large siphoviruses with genomes of 101-116 kb length (Fig 7A-C). Our nine new isolates in the BASEL collection (plus well-studied phage T5) are well representative of the two major genera *Eseptimavirus* and *Tequintavirus* and their subclades (Fig 7B). These phages characteristically use their lateral tail fibers to bind each one or a few types of O-antigen very specifically as their primary host receptor, but this interaction is not essential for hosts like *E. coli* K-12 laboratory strains that don’t express an O-antigen barrier [51] (Fig 7A). As expected and observed previously [51, 60], the diversity of O-antigen glycans is reflected in a high genetic diversity at the lateral tail fiber locus of the *Markadamsvirinae* of the BASEL collection (S3A Fig). Only one of these phages, IrisVonRoten (Bas32), can infect *E. coli* K-12 with restored expression of O16-type O-antigen, suggesting that it can recognize this glycan as its primary receptor, but several others can infect different *E. coli* strains with smooth LPS or even *Salmonella* (see below in Fig 12). Similar to what was proposed for the different small siphoviruses described above, the terminal receptor specificity of T5 and relatives is determined by dedicated RBPs attached non-covalently to the tip of a straight central tail fiber [51]. Compared to their smaller relatives, the repertoire of terminal receptors among *Markadamsvirinae* seems limited – the vast majority target BtuB (shown for, e.g., EPS7 [61] and S132 [62]), and each a few others bind FhuA (T5 itself [51] and probably S131 [62]) or FepA (H8 and probably S124 [62]). Sequence analyses of the RBP locus allowed us to predict the terminal receptor of all remaining *Markadamsvirinae*, confirming that nearly all target BtuB (Fig 7B and S3B Fig).

**Fig 7.**
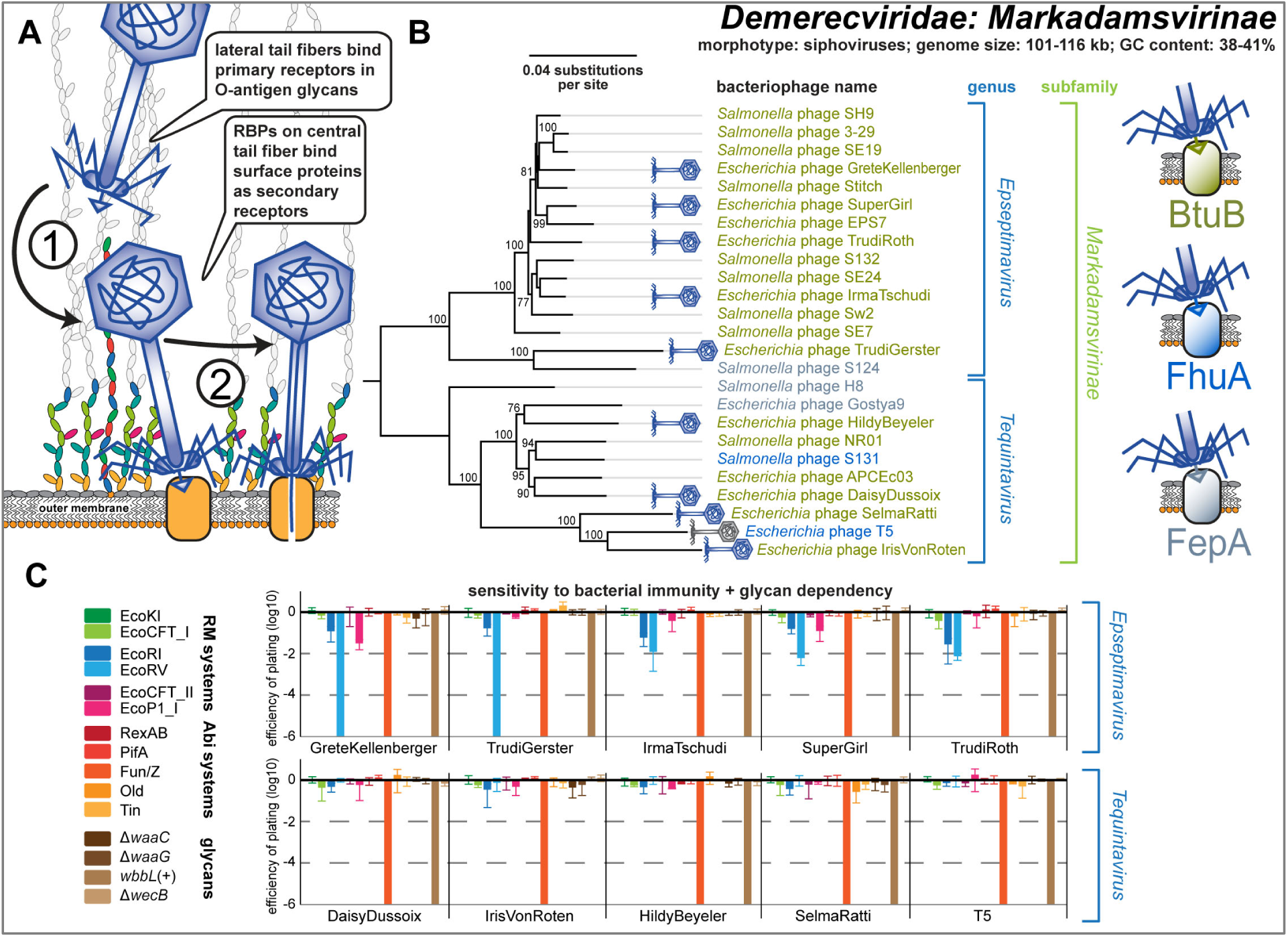
Overview of Demerecviridae subfamily Markadamsvirinae. **(A)** Schematic illustration of host recognition by T5-like siphoviruses. **(B)** Maximum-Likelihood phylogeny of the *Markadamsvirinae* subfamily of *Demerecviridae* based on several core genes with bootstrap support of branches shown if > 70/100. Phages of the BASEL collection are highlighted by little phage icons and the determined or proposed terminal receptor specificity is highlighted at the phage names using the color code highlighted at the right side (same as for the small siphoviruses). The phylogeny was rooted between the *Epseptimavirus* and *Tequintavirus* genera. **(C)** The results of quantitative phenotyping experiments with *Markadamsvirinae* phages regarding sensitivity to altered surface glycans and bacterial immunity systems are presented as efficiency of plating (EOP). Data points and error bars represent average and standard deviation of at least three independent experiments.

It has long been known that phage T5 is largely resistant to RM systems, and this was thought to be due to elusive DNA protection functions encoded in the early-injected genome region largely shared by T5 and other *Markadamsvirinae* [64]. Surprisingly, we find that this resistance to restriction is a shared feature only of the *Tequintavirus* genus, while their sister genus *Epseptimavirus* shows detectable yet variable sensitivity in particular to the type II RM system EcoRV (Fig 7B). It seems likely that this difference between the genera is due to the largely different number of EcoRV recognition sites in *Epseptimavirus* genomes (70-90 sites, S5 Table) and *Tequintavirus* genomes (4-18 sites, S5 Table). This observation suggests that efficient DNA ligation and RM site avoidance [65] play important roles in the restriction resistance of T5 and relatives besides their putative DNA protection system. Remarkably, all *Markadamsvirinae* are invariably sensitive to the Fun/Z immunity system of phage P2 which had already been shown for T5 previously [50].

### Properties of *Myoviridae: Tevenvirinae*

Phages of the *Tevenvirinae* subfamily within the *Myoviridae* family are large myoviruses with characteristically prolate capsids and genomes of 160-172 kb size that infect a wide variety of Gram-negative hosts [30] (Figs 8A-E). The kinked lateral tail fibers of these phages contact primary receptors on the bacterial cell surface that are usually surface proteins like OmpC, Tsx, and FadL for prototypic *Tevenvirinae* phages T4, T6, and T2, respectively, but can also be sugar motifs in the LPS like in case of T4 when OmpC is not available [66] (Fig 8A). Robust interaction with the primary receptor unpins the short tail fibers from the myovirus baseplate which enables their irreversible adsorption to terminal receptors in the LPS core followed by contraction of the tail sheath. Consequently, the phage tail penetrates the cell envelope and the viral genome is injected via a syringe-like mechanism [35] (Fig 8A).

**Fig 8.**
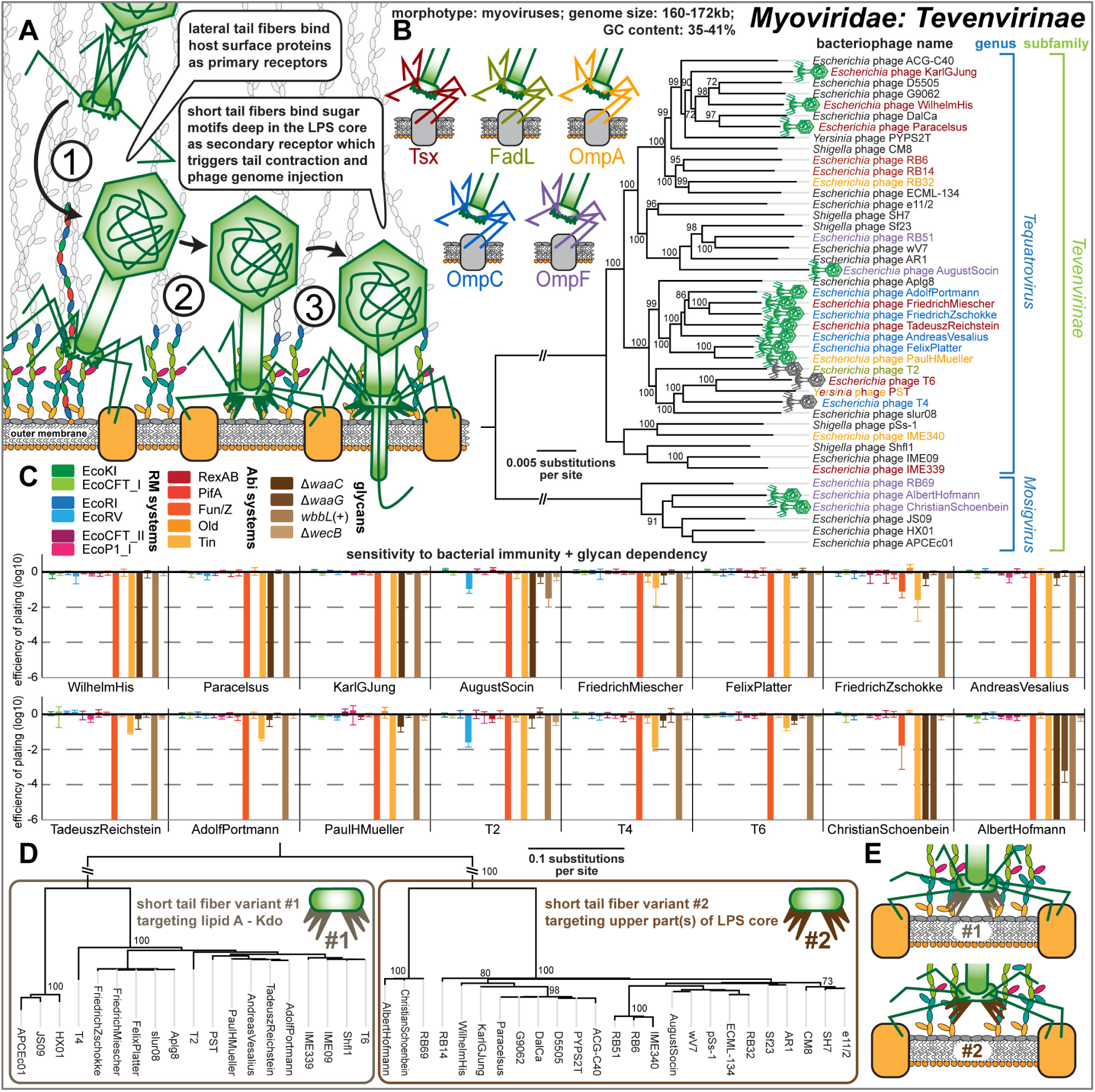
Overview of the *Myoviridae* subfamily *Tevenvirinae*. **(A)** Schematic illustration of host recognition by T4-like myoviruses. **(B)** Maximum-Likelihood phylogeny of the *Tevenvirinae* subfamily of *Myoviridae* based on a curated whole-genome alignment with bootstrap support of branches shown if > 70/100. The phylogeny was rooted between the *Tequatrovirus* and *Mosigvirus* genera. Phages of the BASEL collection are highlighted by little phage icons and experimentally determined primary receptor specificity is highlighted at the phage names using the color code highlighted at the top left. Primary receptor specificity of *Tevenvirinae* depends on RBPs expressed either as a C-terminal extension of the distal half fiber (T4 and other OmpC-targeting phages) or as separate small fiber tip adhesins [66], but sequence analyses of the latter remained ambiguous. We therefore only annotated experimentally determined primary receptors (see also S4A Fig) [53, 66]. **(C)** The results of quantitative phenotyping experiments with *Tevenvirinae* phages regarding sensitivity to altered surface glycans and bacterial immunity systems are presented as efficiency of plating (EOP). Data points and error bars represent average and standard deviation of at least three independent experiments. **(D)** The Maximum-Likelihood phylogeny *Tevenvirinae* short tail fiber proteins reveals two homologous, yet clearly distinct, clusters that correlate with the absence (variant #1, like T4) or presence (variant #2) of detectable LPS core dependence as shown in (C). **(E)** The results of (D) indicate that variant #1, as shown for T4, binds the deep lipid A – Kdo region of the enterobacterial LPS core, while variant #2 binds a more distal part of the (probably inner) core.

Our thirteen newly isolated *Tevenvirinae* phages in the BASEL collection mostly belong to different groups of the *Tequatrovirus* genus that also contains well-studied reference phages T2, T4, and T6, but two isolates were assigned to the distantly related *Mosigvirus* genus (Fig 8B). As expected, the infectivity of our thirteen isolates and of T2, T4, and T6 reference phages shows strong dependence on each one of a small set of *E. coli* surface proteins that have previously been described as *Tevenvirinae* primary receptors, i.e., Tsx, OmpC, OmpF, OmpA, and FadL [66] (Fig 8B). Interestingly, while for some of these phages the absence of the primary receptor totally abolished infectivity, others still showed detectable yet greatly reduced plaque formation (S4A Fig). It seems likely that these differences are caused by the ability of some *Tevenvirinae* lateral tail fibers to contact several primary receptors such as, e.g., OmpC and the truncated *E. coli* B LPS core in case of T4 [66, 67].

Specificity for the secondary receptor depends on the short tail fibers that, in case of T4, target the lipid A – Kdo region deep in the enterobacterial LPS core [68]. Given that this region is still present in the *waaC* mutant, the most deep-rough mutant of *E. coli* K-12 that is viable (Fig 3A), it is unsurprising that T4 and some other *Tevenvirinae* did not seem to show a dependence on the LPS core in our experiments (Fig 8C). However, some *Tequatrovirus* and all tested *Mosigvirus* isolates required an intact inner core of the host LPS for infectivity (Fig 8C). This phenotype is correlated with an alternative allele of the short tail fiber gene that varies between *Tevenvirinae* phages irrespective of their phylogenetic position (Fig 8D). We therefore suggest that those phages encoding the alternative allele express a short tail fiber that targets parts of the LPS core above the lipid A – Kdo region (Figs 8D and 8E; see also Fig 3A).

Besides receptor specificity, the phenotypes of our diverse *Tevenvirinae* phages were highly homogeneous. The hallmark of this *Myoviridae* subfamily is a high level of resistance to DNA-targeting immunity like RM systems because of cytosine hypermodifications (hydroxymethyl-glucosylated for *Tequatrovirus* and hydroxymethyl-arabinosylated for *Mosigvirus*) [46, 69]. Consistently, no or only very weak sensitivity to any of the six RM systems was detected for any tested *Tevenvirinae* phage (Fig 8B). The weak sensitivity to EcoRV observed for AugustSocin (Bas38) and phage T2 does, unlike for *Markadamsvirinae* (see above), not correlate with a higher number of recognition sites and therefore possibly depends on another feature of these phages such as, e.g., differences in DNA ligase activity (S5 Table). Conversely, all tested *Tevenvirinae* were sensitive to the Fun/Z and Tin Abi systems (Fig 8B). Tin was previously shown to specifically target the DNA replication of T-even phages [50], and we found that indeed all *Tevenvirinae* but no other tested phage are sensitive to this Abi system (Fig 8B). No sensitivity was observed for RexAB, the iconic Abi system targeting *rIIA/B* mutants of phage T4 [10] (S4B), suggesting that this Abi system is generally unable to affect wildtype *Tevenvirinae* phages (Fig 8C).

### Properties of *Myoviridae: Vequintavirinae* and relatives

Phages of the *Vequintavirinae* subfamily within the *Myoviridae* family are myoviruses with genomes of 131-140 kb size that characteristically encode three different sets of lateral tail fibers [23, 70] (Figs 9A-E). Besides *Vequintavirinae* of the *Vequintavirus* genus, this feature is shared by two groups of phages that are closely related to these *Vequintavirinae sensu stricto*: One group forms a cluster around phage phAPEC8 (recently proposed as a new genus *Phapecoctavirus* within *Myoviridae* [23]), the other group is their sister clade including phage phi92 [71] that is unclassified (Fig 9B). Despite not being *Vequintavirinae* by current taxonomic classification, we are covering them together due to their considerable similarities and propose to classify the phi92-like phages (including PaulScherrer (Bas60)) as *Nonagintaduovirus* genus within the *Vequintavirinae*.

**Fig 9.**
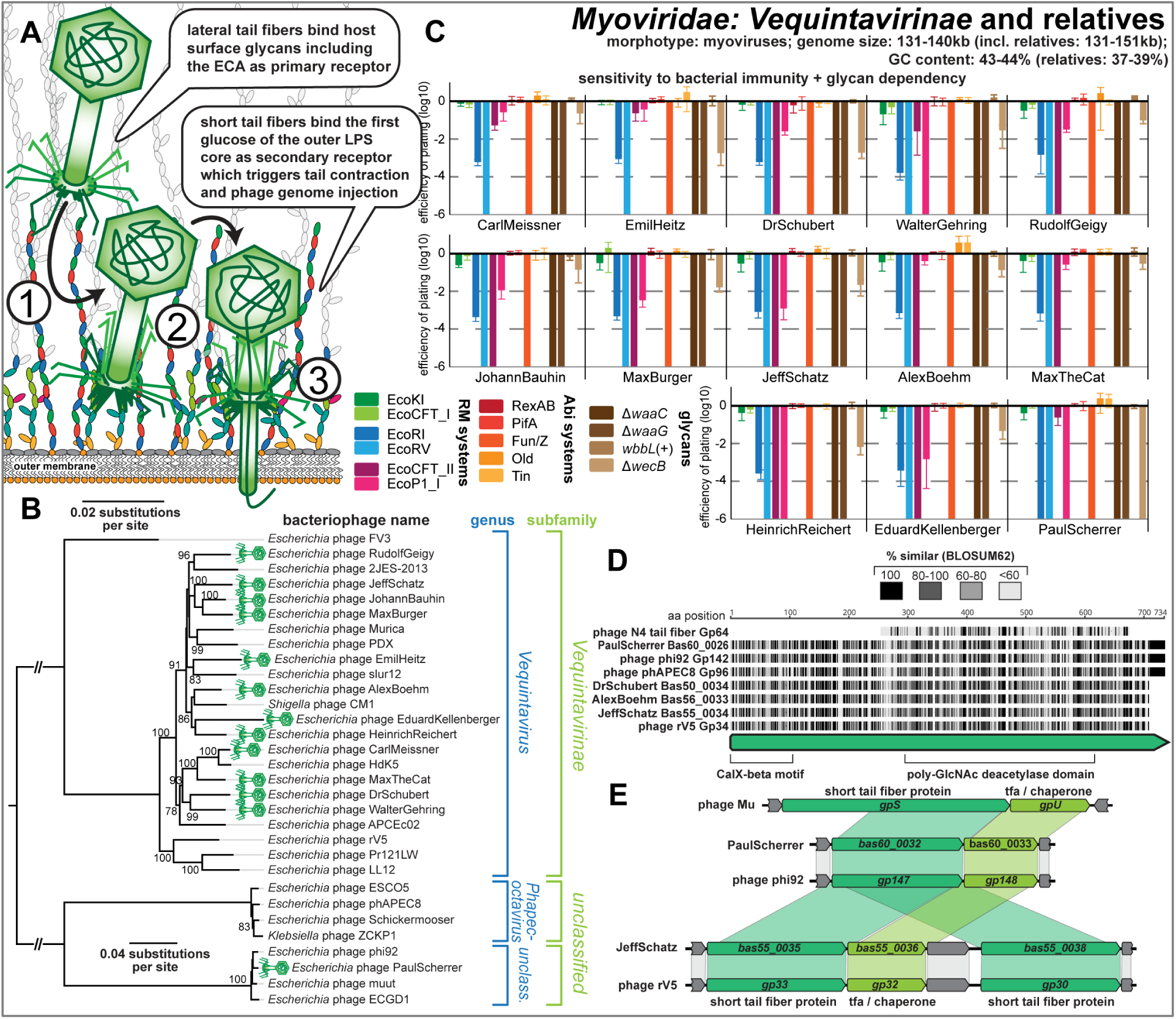
Overview of the *Myoviridae* subfamily *Vequintavirinae* and relatives. (**A**) Schematic illustration of host recognition by *Vequintavirinae* and related myoviruses. **(B)** Maximum-Likelihood phylogeny of the *Vequintavirinae* subfamily of *Myoviridae* and relatives based on a curated whole-genome alignment with bootstrap support of branches shown if > 70/100. The phylogeny was rooted between the *Vequintavirus* genus and the two closely related, unclassified groups at the bottom. Newly isolated phages of the BASEL collection are highlighted by green phage icons. **(C)** The results of quantitative phenotyping experiments with *Vequintavirinae* and phage PaulScherrer regarding sensitivity to altered surface glycans and bacterial immunity systems are presented as efficiency of plating (EOP). Data points and error bars represent average and standard deviation of at least three independent experiments. **(D)** Amino acid sequence alignment of the lateral tail fiber Gp64 of phage N4 (*Enquatrovirus*, see below) and a lateral tail fiber conserved among *Vequintavirinae* and relatives (representatives shown). The proteins share a predicted poly-GlcNAc deacetylase domain as identified by Phyre2 [136]. **(E)** *Vequintavirinae sensu stricto* (represented by rV5 and Jeff Schatz) encode two paralogous short tail fiber proteins and a tail fiber chaperone that are homologous to the corresponding locus in phi92-like phages incl. PaulScherrer and, ultimately, to short tail fiber GpS and chaperone GpU of Mu(+) (which targets a different glucose in the K12-type LPS core GpS [95, 96]).

Three different, co-expressed lateral tail fibers have been directly visualized in the cryogenic electron microscopy structure of phi92, and large orthologous loci are found in all *Vequintavirinae* and relatives [23, 70, 71]. The considerable repertoire of glycan-hydrolyzing protein domains in these tail fiber proteins has been compared to a “nanosized Swiss army knife” and is probably responsible for the exceptionally broad host range of these phages and their ability to infect even diverse capsulated strains of enterobacteria [23, 52, 70, 71]. However, it is far from clear which genes code for which components of the different lateral tail fibers. While three lateral tail fiber genes exhibit considerable allelic variation between phage genomes and might therefore encode the distal parts of the tail fibers with the receptor-binding domains, the biggest part of the lateral tail fiber locus is highly conserved including diverse proteins with sugar-binding or glycan-hydrolyzing domains (S5 Fig). This observation fits well with our finding that the lysis host range of all *Vequintavirinae* available to us is remarkably homogeneous and suggests that, in the absence of capsules and other recognized exopolysaccharides, these phages do not vary widely in their host recognition (see below in *Host range across pathogenic enterobacteria and* E. coli *B*). Interestingly, all *Vequintavirinae* in the BASEL collection are detectably inhibited when infecting the *wecB* knockout (albeit to variable degree), strongly suggesting that the ECA is a shared primary receptor (Figs 9A and 9B). This phenotype and the specific lysis host range of *Vequintavirinae* are shared with podoviruses of the *Enquatrovirus* family (see below). Notably, one of the conserved lateral tail fiber proteins of the *Vequintavirinae* is homologous to the lateral tail fiber protein of *Enquatrovirus* phages (Fig 9D), suggesting that these two groups of phages might target the ECA in a similar way.

The terminal receptor of *Vequintavirinae* has been unraveled genetically for phage LL12, a close relative of rV5 (Fig 9B), and seems to be at the first, heptose-linked glucose of the LPS outer core which is shared by all *E. coli* core LPS types [52, 72] (Fig 9A). Consistently, we found that all *Vequintavirinae* in the BASEL collection are completely unable to infect the *waaC* and *waaG* mutants with core LPS defects that result in the absence of this sugar (Fig 9C, see also Fig 3A). This observation would be compatible with the idea that all (tested) *Vequintavirinae* and also PaulScherrer of the phi92-like phages use the same secondary receptor. Indeed, the two very similar short tail fiber paralogs of *Vequintavirinae sensu stricto* do not show any considerable allelic variation between genomes (S5A Fig; unlike, e.g., the *Tevenvirinae* short tail fibers presented in Figs 8D and 8E) and are also closely related to the single short tail fiber protein of phi92-like phages or *Phapecoctavirus* (Fig 9E).

Unlike large myoviruses of the *Tevenvirinae* subfamily, the *Vequintavirinae* and relatives are exceptionally susceptible to type II and type III RM systems, albeit with (minor) differences in the pattern of sensitivity and resistance from phage to phage (Fig 9C). We therefore see no evidence for effective mechanisms of these phages to overcome RM systems like, as proposed previously, covalent DNA modifications which could be introduced by the notable repertoire of sugar-related enzymes found in their genomes [24]. Similarly, the *Vequintavirinae* are invariably sensitive to the Fun/Z Abi system.

### Properties of the *Autographiviridae* family and *Podoviridae: Enquatrovirus*

Phages of the *Autographiviridae* family and the genus *Enquatrovirus* in the *Podoviridae* family are podoviruses that have been well studied in form of their representatives T3 and T7 (*Autographiviridae* subfamily *Studiervirinae*) and N4 (*Enquatrovirus*). While they differ in their genome size (ca. 37-41 kb for *Studiervirinae*, ca. 68-74 kb for *Enquatrovirus*) and each exhibit characteristic unique features, the overall mode of infection of these podoviruses is the same: Lateral tail fibers of the virion contact bacterial surface glycans at the cell surface for host recognition after which direct contact of the stubby tail with a terminal receptor on the host triggers irreversible adsorption and DNA injection (Fig 10A).

**Fig 10.**
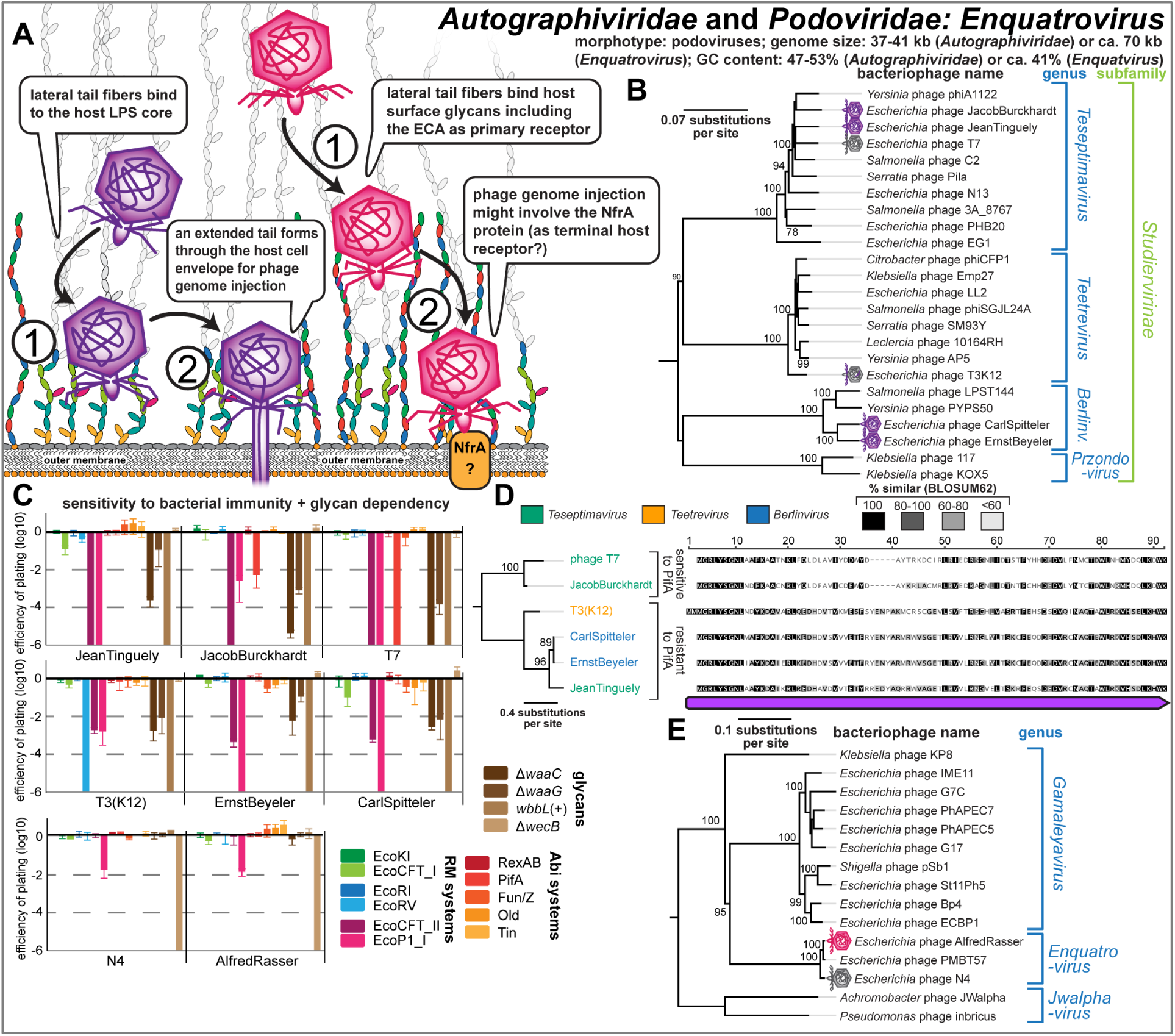
Overview of Autographiviridae phages and Podoviridae genus Enquatrovirus. **(A)** Schematic illustration of host recognition by *Autographiviridae* and *Enquatrovirus* phages. **(B)** Maximum-Likelihood phylogeny of the *Studiervirinae* subfamily of *Autographiviridae* based on several core genes with bootstrap support of branches shown if > 70/100. The phylogeny was midpoint-rooted between the clade formed by *Teseptimavirus* and *Teetrevirus* and the other genera. Phages of the BASEL collection are highlighted by little phage icons. **(C)** The results of quantitative phenotyping experiments with *Autographiviridae* regarding sensitivity to altered surface glycans and bacterial immunity systems are presented as efficiency of plating (EOP). Data points and error bars represent average and standard deviation of at least three independent experiments. **(D)** Amino acid sequence alignment and Maximum-Likelihood phylogeny of Gp1.2 orthologs in all tested *Autographiviridae* phages. Phage JeanTinguely belongs to the *Teseptimavirus* genus but encodes an allele of *gp1.2* that is closely related to those of the *Berlinvirus* genus, possibly explaining its resistance to PifA (see (C)). **(E)** Maximum-Likelihood phylogeny of the *Enquatrovirus* genus and related groups of *Podoviridae* based on several core genes with bootstrap support of branches shown if > 70/100. The phylogeny was midpoint-rooted between the distantly related *Jwalphavirus* genus and the others. Phages of the BASEL collection are highlighted by little phage icons.

This process has been studied in detail for the archetype of all *Autographiviridae*, phage T7, though several open questions remain. The lateral tail fibers of this phage contact a receptor in rough LPS of *E. coli* K-12 that is not fully understood, possibly because several alternative and overlapping sugar motifs in the K-12 LPS core can be targeted [73–75] (Fig 10A). Conformational changes in the stubby tail tube triggered by this receptor interaction then initiate the injection of the phage genome from the virion [75, 76]. Remarkably, at first several internal virion proteins are ejected and then fold into an extended tail that spans the full bacterial cell envelope which, in a second step, enables DNA injection into the host cytosol [77] (Fig 10A). Notably, our knowledge of this process does not allow the distinction between a “primary” receptor for host recognition and a “secondary” terminal receptor for irreversible adsorption and DNA injection. However, other *Autographiviridae* follow this classical scheme more closely and feature enzymatic domains at their tail fibers or tail spikes that likely mediate attachment to surface glycans as primary receptors followed by enzyme-guided movement towards the cell surface [38, 78].

The *Autographiviridae* are a large family of podoviruses hallmarked (and named) with reference to their single-subunit T3/T7-type RNA polymerase that plays several key roles for the phage infection but has also become a ubiquitous tool in biotechnology [79]. All *Autographiviridae* isolates of the BASEL collection belong to several different genera within the very broad *Studiervirinae* subfamily that also contains the classical T7 and T3 phages which we included as references (Fig 10B). Because phage T3 recognizes the peculiarly truncated R1-type LPS core of *E. coli* B (see also below) and cannot infect K-12 strains, we generated a T3(K12) chimera that encodes the lateral tail fiber gene of T7 similar as was reported previously by others (see *Materials and Methods*) [80]. As expected, all tested *Autographiviridae* use core LPS structures as host receptor and show impaired plaque formation on *waaC* and *waaG* mutants, but in almost all cases some infectivity is retained even on the *waaC* mutant (Fig 10C). This suggests that, as postulated for T7 [73–75], these phages are not strictly dependent on a single glycan motif but might recognize a broader range target structures at the LPS core.

The overall pattern of restriction sensitivity and resistance is similar for all tested *Autographiviridae*. While type I and type II RM systems are largely ineffective, type III RM systems show remarkable potency across all phage isolates. Good part of the resistance to type II RM systems is likely due to the near-complete absence of EcoRI and EcoRV recognition sites in the genomes of these phages as described previously [65] (S5 Table) with the exception of ten EcoRV sites in T3(K12) which, consequently, cause massive restriction (Fig 10C). The relatively few recognition sites for type I RM systems do not result in considerable sensitivity for any phage, because at the *gp0.3* locus all of them either encode an Ocr-type DNA mimic type I restriction inhibitor like T7 (*Teseptimavirus*) or an adenosylmethionine (SAM) hydrolase that deprives these RM systems of their substrate like T3 (all other genera) [9, 81, 82]. Notably, phages JacobBurckhardt (Bas63) and its close relative T7 are the only phages tested in this work that show any sensitivity to the PifA Abi system encoded on the *E. coli* K-12 F-plasmid (Fig 10C). Previous work showed that sensitivity of T7 to PifA immunity – as opposed to T3 which is resistant – depends on the dGTPase Gp1.2 of the phage, though the major capsid protein Gp10 also seems to play some role in sensitivity [83, 84]. Consequently, we find that phages T7 and JacobBurckhardt, but not closely related *Teseptimavirus* JeanTingely (Bas62), encode a distinct variant of dGTPase Gp1.2 that likely causes their sensitivity to PifA (Fig 10D).

Bacteriophage N4 is the archetype of *Enquatrovirus* phages that are hallmarked by using a large, virion-encapsidated RNA polymerase for the transcription of their early genes [33], and our new *Enquatrovirus* isolate AlfredRasser (Bas67) is a very close relative of N4 (Fig 10E). Based primarily on genetic evidence, phage N4 is thought to initiate infections by contacting the host’s ECA with its lateral tail fibers [73, 85, 86] (Fig 10A). We indeed confirmed a remarkable dependence of N4 and AlfredRasser on *wecB* (Fig 10C) and detected homology between the glycan deacetylase domain of the N4 lateral tail fiber and a lateral tail fiber protein of *Vequintavirinae* which also seem to use the ECA as their primary receptor (see above, Figs 9 and 10C). Similar to the O-antigen deacetylase tail fiber of its relative G7C (Fig 10E), the enzymatic activity of the N4 tail fiber might provide a directional movement towards the cell surface that is essential for the next steps of infection [38, 87] (Fig 10A). At the cell surface, the elusive outer membrane porin NfrA was suggested to be the terminal receptor of phage N4 based on the findings that it is required for N4 infection and interacts with the stubby tail of this phage [73, 85, 88]. To the best of our knowledge, this would make phage N4 the first podovirus with an outer membrane protein as terminal receptor [39, 40], though the absence of homologs to the tail extension proteins of *Autographiviridae* indeed suggests a somewhat different mode of DNA injection (Fig 10A).

With regard to bacterial immunity, *Enquatrovirus* phages are highly resistant to any tested antiviral defenses with exception to a slight sensitivity to the EcoP1_I type III RM system (Fig 10C). While their resistance to tested type II RM systems is due to the absence of EcoRI or EcoRV recognition sites (S5 Table), the genetic basis for their only slight sensitivity to type III RM systems is not known. We note that *Enquatrovirus* phages encode an *rIIAB* locus homologous to the long-known yet poorly understood *rIIAB* locus found in many large myoviruses which, in case of phage T4, provides resistance to the RexAB Abi system of phage lambda [89] (S4B and S4C Figs). Both phage N4 and AlfredRasser are indifferent to the presence of (O16-type) O-antigen or the tested truncations of the K-12 LPS core at the host cell surface (Fig 10C), suggesting that LPS structures play no role for their infection process.

### Properties of *Myoviridae*: *Ounavirinae* and classical temperate phages

The *Felixounavirus* genus in the *Ounavirinae* subfamily of *Myoviridae* comprises a group of phages with genomes of 84-91 kb and characteristically straight lateral tail fibers of which *Salmonella* phage Felix O1 has been most well studied [30, 90] (Figs 11A and 11B). Previous work aimed at the isolation of *E. coli* phages differed greatly in the reported abundance of *Felixounavirus* phages, ranging from no detection [23] over a moderate number of isolates in most studies [25, 91] to around one fourth of all [24]. Across our phage sampling experiments we found a single *Felixounavirus* phage, JohannRWettstein (Bas61), which is rather distantly related to Felix O1 within the eponymous genus (Fig 11B). Prototypic phage Felix O1 has been used for decades in *Salmonella* diagnostics because it lyses almost every *Salmonella* strain but only very few other *Enterobacteriaceae* (reviewed in reference [90]). The mechanism of its host recognition and the precise nature of its host receptor(s) have remained elusive, but it is generally known to target bacterial LPS and not any kind of protein receptors [90]. While many *Ounavirinae* seem to bind O-antigen glycans of smooth LPS, the isolation of multiple phages of this subfamily on *E. coli* K-12 with rough LPS and on *E. coli* B with an even further truncated LPS indicate that the functional expression of O-antigen is not generally required for their host recognition [24, 90, 92]. Phage JohannRWettstein is only slightly inhibited on *E. coli* K-12 with restored O16-type O-antigen, suggesting that it can either use or bypass these glycans, and totally depends on an intact LPS core (Fig 11E). In addition, it is remarkably sensitive to several tested RM systems and shares sensitivity to the Fun/Z Abi system with all other tested *Myoviridae* and the *Markadamsvirinae* (Fig 11E).

**Fig 11.**
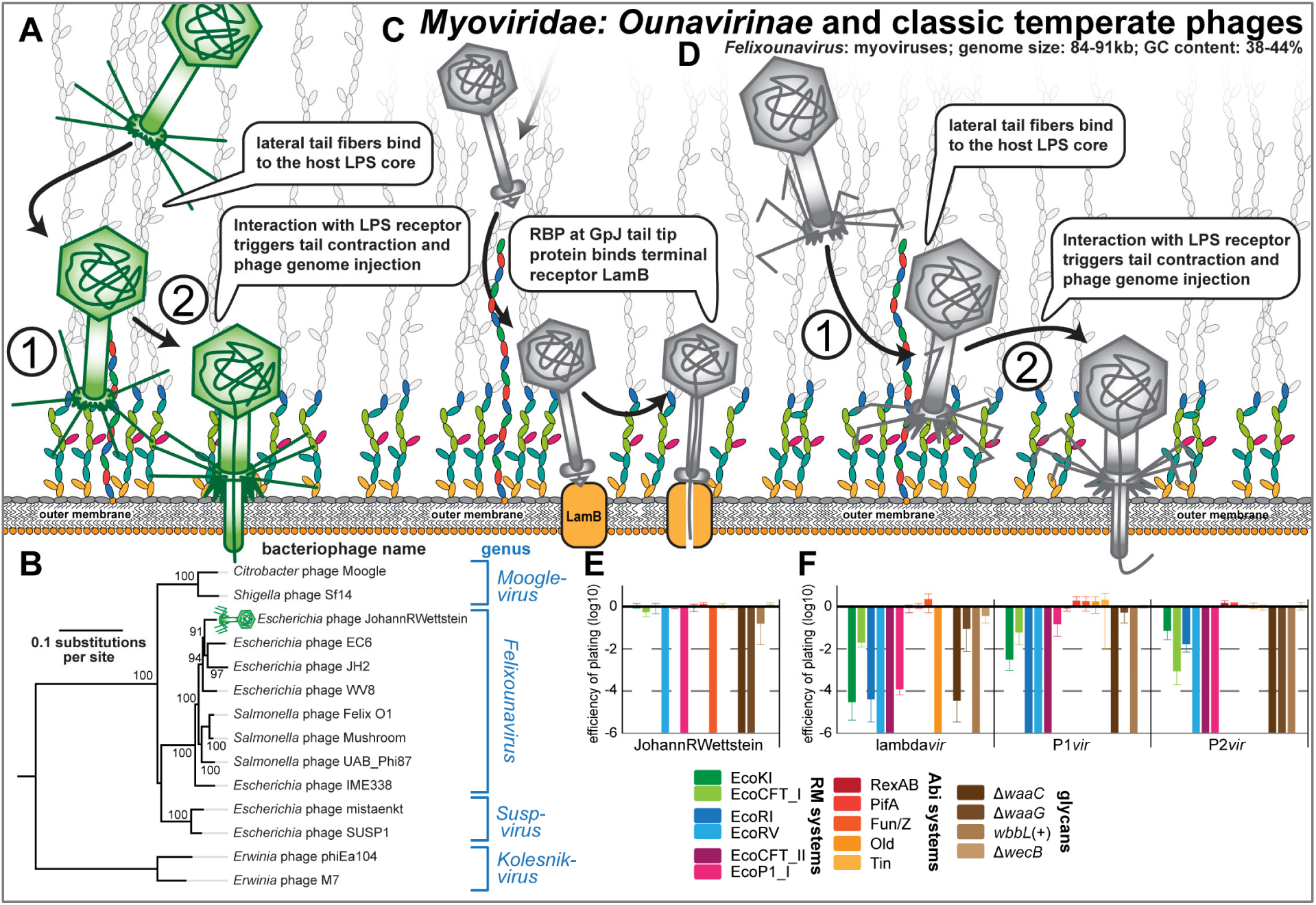
Overview of *Myoviridae: Ounavirinae* and classic temperate phages. **(A)** Schematic illustration of host recognition by *Ounavirinae: Felixounavirus* phages. Note that the illustration shows short tail fibers simply in analogy to *Tevenvirinae* or *Vequintavirinae* (Figs 8A and 9A), but any role for such structures has not been explored for Felix O1 and relatives. **(B)** Maximum-Likelihood phylogeny of the *Ounavirinae* subfamily of *Myoiviridae* based on several core genes with bootstrap support of branches shown if > 70/100. The phylogeny was midpoint-rooted between *Kolesnikvirus* and the other genera. Our new isolate JohannRWettstein is highlighted by a green phage icon. **(C,D)** Schematic illustration of host recognition by classic temperate phages lambda, P1, and P2. Note the absence of lateral tail fibers due to a mutation in lambda PaPa laboratory strains [94]. **(E,F)** The results of quantitative phenotyping experiments with JohannRWettstein and classic temperate phages regarding sensitivity to altered surface glycans and bacterial immunity systems are presented as efficiency of plating (EOP). Data points and error bars represent average and standard deviation of at least three independent experiments.

Besides the lytic T phages, temperate phages lambda, P1, and P2 have been extensively studied both as model systems for fundamental biology questions as well as regarding the intricacies of their infection cycle [7, 34, 50]. Phage lambda was a prophage encoded in the original *E. coli* K-12 isolate and forms siphovirus particles that display the GpJ tail tip to contact the LamB porin as the terminal receptor for DNA injection [7, 93], while the lateral tail fibers (thought to contact OmpA as primary receptor) are missing in most laboratory strains of this phage due to a mutation [7, 94] (Fig 11C). We reproduced the dependence of our lambda*vir* variant on LamB and, as reported previously, found that an intact inner core of the LPS was required for infectivity [73 and literature cited therein] (Fig 11F and S5 Table). Myoviruses P1 and P2 were both shown to require a rough LPS phenotype to contact their receptors in the LPS core, but the exact identity of these receptors has not been unraveled [40, 50, 73, 95]. While the molecular details of the infection process after adsorption have not been well studied for P1, previous work showed that an interaction of the P2 lateral tail fibers with the LPS core of K-12 strains triggers penetration of the outer membrane by the tail tip [40, 50, 96] (Fig 11D). Consistently, our results confirm that mutations compromising the integrity of the K-12 core LPS abolished bacterial sensitivity to P1*vir* and P2*vir* [50, 73, 95] (Fig 11F). The quantification of these phages’ sensitivity to different immunity systems revealed that they are remarkably sensitive to all tested RM systems (Fig 11F), a property only shared with a few *Drexlerviridae* (Fig 4D). As expected from the considerably lower number of restriction sites, the sensitivity was less pronounced for type I RM systems (Fig 11F and S5 Table). Phage lambda was additionally specifically sensitive to the Old Abi system of the P2 prophage (Fig 11F), as shown previously [50].

### Host range across pathogenic enterobacteria and laboratory wildtype *E. coli* B

The host range of bacteriophages has repeatedly been highlighted as a critical feature for phage therapy because broad infectivity can enable phages to be used against different strains of the same pathogen without repeated sensitivity testing “analogous to the use of broad-spectrum antibiotics” [16]. Intuitively, across strains of the same host species the infectivity of phages depends on their ability to successfully bind the variable surface structures of host cells and to overpower or evade strain-specific bacterial immunity [16, 97]. While simple qualitative tests assessing the lysis host range primarily inform about host recognition alone, the more laborious detection of robust plaque formation as a sign of full infectivity is the gold standard of host range determination [16, 97]. We therefore challenged a panel of commonly used pathogenic enterobacterial strains with the BASEL collection and recorded the phages’ lysis host range as well as plaque formation separately to gain insight into their host recognition and the ability to overcome immunity barriers inside host cells (Fig 12; see *Materials and Methods*).

**Fig 12.**
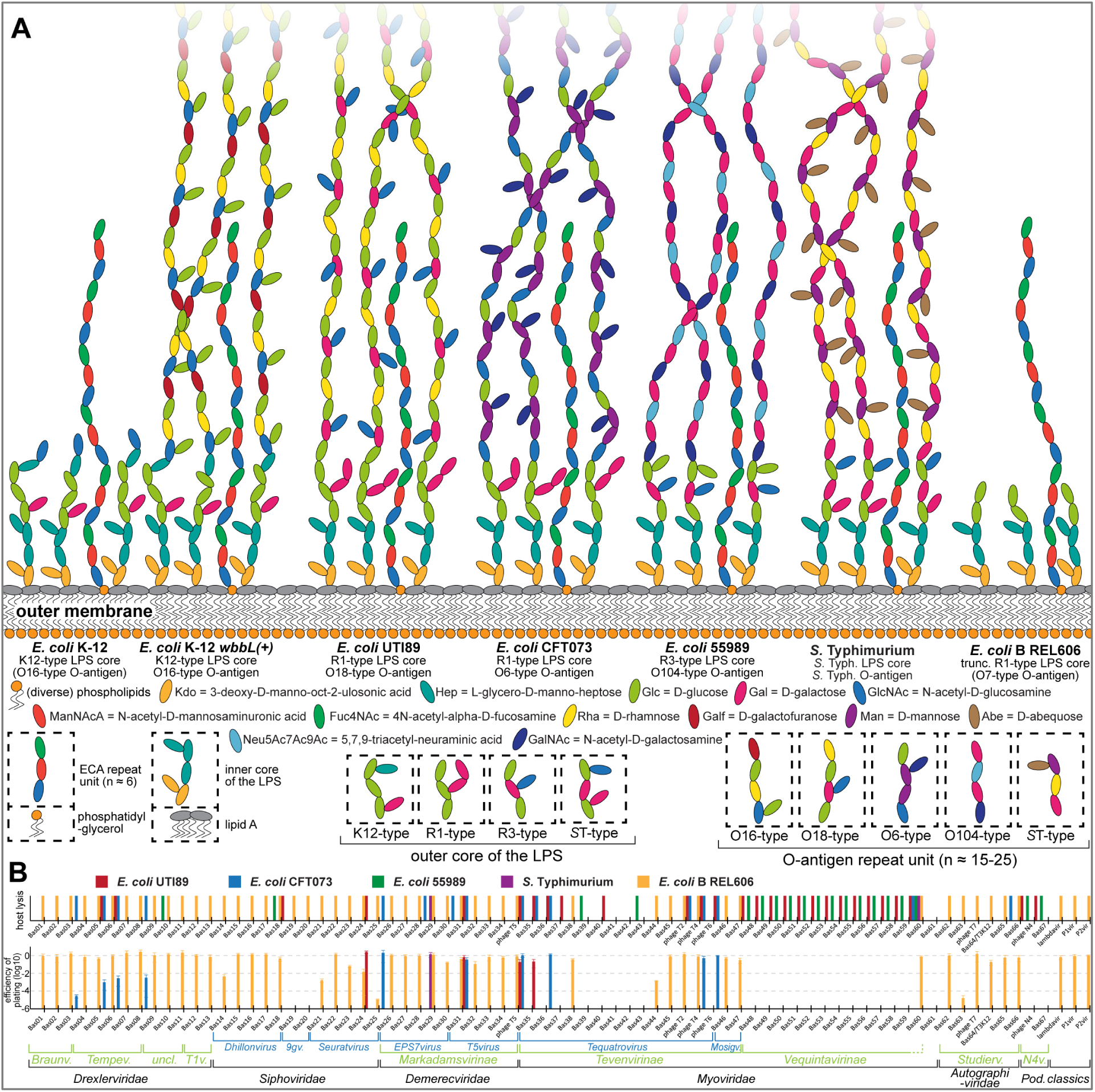
Host range of phages in the BASEL collection. **(A)** Surface glycans of the enterobacterial strains used in this work (see *Materials and Methods* for details on how the illustration of glycan chains was composed). **(B)** The ability of all phages in the BASEL collection to infect different enterobacteria was studied qualitatively (lysis host range; top) and, more stringently, based on the ability to form plaques (bottom). Top: The observartion of lysis zones with high-titer lysate (>10^9^ pfu/ml) in at least three independent experiments is indicated by colored bars. Bottom: The infectivity of BASEL collection phages on diverse enterobacterial hosts was quantified as efficiency of plating (EOP). Data points and error bars represent average and standard deviation of at least three independent experiments. Since the data obtained with *Salmonella* Typhimurium 12023s and SL1344 were indistinguishable, we only show the results of one representative strain (*S.* Typhimurium 12023s).

As expected from previous work, *Vequintavirinae* phages showed an outstanding lysis host range [23, 25, 70, 71] and invariably infected *E. coli* UTI89, *E. coli* 55989, and *E. coli* K-12 with restored O16-type O-antigen (Figs 9C and 12B). The diversity of infected hosts suggests that *Vequintavirinae* can readily bypass the O-antigen barrier, but the homogeneity of their lysis host ranges is clearly at odds with the polyvalent nature of their virions that display several different lateral tail fibers and variable RBPs [70, 71] (S5 Fig). Notably, we showed that *Enquatrovirus* phages share the exact same host range and an insensitivity to the O-antigen barrier with *Vequintavirinae* (Figs 10C and 12B), possibly because these phages target the highly conserved ECA as primary receptor using a homologous tail fiber (Fig 9D). Recognition of the ECA molecules among smooth LPS (Fig 12A) might enable *Vequintavirinae* and *Enquatrovirus* to bypass the O-antigen barrier and move to the cell surface along this glycan chain, possibly using the deacetylase domain of their shared tail fiber as previously described for *Salmonella* phage P22 and the N4 relative G7C [38, 87] (Fig 10E). If true, the *Vequintavirinae* would be effectively monovalent on the tested hosts and might primarily use their “nanosized Swiss army knife” of tail fibers to bind and overcome capsules or other more specialized exopolysaccharides [70, 71]. Given the remarkable lysis host range of *Vequintavirinae* and *Enquatrovirus*, it would be valuable to explore the molecular basis of their host recognition in future studies in order to use this knowledge for phage host range engineering [14].

A decently broad lysis host range was also observed for *Tevenvirinae* phages (Fig 12B), in line with previous work [23], though more scattered and far less remarkable than for the *Vequintavirinae*. For both groups of phages it is astonishing how poorly their broad host recognition is reflected by actual plaque formation (Fig 12B). Despite robust “lysis-from-without” in some cases [98], no reliable plaque formation on any other host than *E. coli* K-12 was observed for *Vequintavirinae*, and the range of plaque formation of *Tevenvirinae* was also severely contracted relative to the lysis host range (Fig 12B).

Besides *Vequintavirinae* and *Enquatrovirus* that can apparently bypass the O-antigen barrier, we find that the lysis host range of around a third of the tested phages (24 / 61) included at least one host strain with smooth LPS, while the O16-type O-antigen of *E. coli* K-12 blocked infections by all but five of these phages scattered across the different taxonomic groups (5 / 61; Fig 12; see also Figs 4 and 6-11). These results confirm an important role of the O-antigen as a formidable barrier to bacteriophage infection that can only be overcome by specific recognition with tail fibers to breach it, e.g., via enzymatic activities [35]. Notably, less than half of the phages that could lyse any strain besides the *E. coli* K-12 ΔRM isolation host showed robust plaque formation on that strain (Fig 12B), probably due to different layers of bacterial immunity. In comparison to *Vequintavirinae* and *Tevenvirinae*, it is surprising how the lysis host range of *Markadamsvirinae* is almost exactly reflected in their range of plaque formation (Fig 12B). One of these phages, SuperGirl (Bas29) is the only phage in the BASEL collection that shows robust plaque formation on *Salmonella* Typhimurium (Fig 12B).

The *E. coli* B lineage comprises a number of laboratory strains including REL606 that have, like *E. coli* K-12, lost O-antigen expression and even part of its R1-type LPS core during domestication but are significant as the original hosts of the T phages [5, 99]. In line with the barrier function of the O-antigen (see above), the vast majority of phages in the BASEL collection can at least lyse this strain with the notable exception of *Vequintavirinae* and *Enquatrovirus* (Fig 12B). We can only speculate about the molecular basis of this observation, but maybe the *Vequintavirinae* are unable to use the truncated core LPS of this strain for their infections (Figs 12A and 12B; compare Fig 9C). If inherited from earlier *E. coli* B strains, it seems likely that this total resistance to *Vequintavirinae* is the reason why the T phages contain no representative of this group despite their abundance. Similarly, the observation that phage T5 with its FhuA receptor (Fig 7B) is a rather unusual member of the *Markadamsvirinae* can be explained by the loss of *btuB* in older variants of *E. coli* B, but this mutation reverted in an ancestor of *E. coli* B REL606 [100]. Consequently, this modern *E. coli* B strain is sensitive to the various BtuB-targeting *Markadamsvirinae* (Fig 12B). Similarly, *E. coli* B strains lack expression of *ompC* and the REL606 strain additionally lacks functional *tsx* that both encode common primary receptors of the *Tevenvirinae* (Fig 8B) [66, 100]. Though phage T4 itself can use the *E. coli* B LPS core as a primary receptor when OmpC is absent [67], the inability of several other *Tevenvirinae* to infect *E. coli* B REL606 corresponds well to the dependency on OmpC and Tsx as primary receptors (Figs 8B, 12B, and S4A). A number of additional phages of, e.g., the *Nonagvirus* genus can lyse but not form plaques on *E. coli* B REL606 (Fig 12B). Based on previous work, we suggest that this phenotype is caused by immunity systems encoded in the specific repertoire of cryptic prophages of *E. coli* B strains that differ considerably between *E. coli* B and K-12 strains and are known to harbor active immunity systems [100].

## Discussion

### Remarkable patterns of bacteriophage receptor specificity

Our results regarding the terminal receptor specificity of different groups of siphoviruses inherently present the question why all these phages target less than ten of the more than 150 outer membrane proteins of *E. coli* K-12 of which around two dozen are porins [39, 40, 101] (Figs 4C and 7B). Similarly, *Tevenvirinae* use only five different outer membrane proteins as primary receptors (Fig 8B) [39, 40]. Previous work suggested that this bias might be linked to the abundance of the targeted proteins, both via a preference for particularly numerous proteins (favoring phage adsorption) or very scarce ones (avoid competition among phages) [102]. However, we feel that this stark bias in phage preference might be largely driven by functional constraints. Notably, the large siphoviruses of *Markadamsvirinae* bind exactly a subset of those terminal receptors targeted by the small siphoviruses despite using non-homologous RBPs, while there is no overlap between these proteins and the primary receptors of the *Tevenvirinae* [39, 40] (Figs 4-8). It seems therefore likely that the receptors targeted by siphoviruses need to have certain properties, e.g., favoring DNA injection, that greatly limit the repertoire of suitable candidates. We envision that future studies might unravel the molecular mechanisms of how siphoviruses recognize and use outer membrane proteins as terminal receptors on Gram-negative hosts similar to, e.g., how it has been studied for myoviruses and podoviruses that directly puncture the outer membrane [68, 77].

Many open questions remain regarding the host recognition and receptor specificity of *E. coli* phages that could be tackled with a combination of systematic phenotypic and genomic analyses. Why do some small siphoviruses in *Drexlerviridae* and *Siphoviridae* but none of the larger ones of *Demerecviridae: Markadamsvirinae* strongly depend on an intact LPS core, seemingly independent of primary receptor usage or terminal receptor specificity (Figs 4D, 6C, 6F, and 7B)? Is it possible to perform analyses like we did it for the RBPs of siphoviruses or *Tevenvirinae* for the much more variable lateral tail fibers (S1B, S2A, and S3A Figs) that probably bind to the around 200 O-antigen types of *E. coli* [103]? And is there a genetically encoded specificity for the inner membrane channels used by siphoviruses (like ManYZ by phage lambda or PtsG for HK97 [7]) similar to the mix-and-match of RBPs and outer membrane proteins? Addressing these questions in future studies would not only greatly expand our knowledge regarding the molecular basis of bacteriophage ecology and evolution, but would also help making use of this knowledge to optimize the application of phages in biotechnology and for therapeutic applications, e.g., for “phage steering” [14, 104].

### Bacteriophage sensitivity and resistance to host immunity

Our systematic analysis of the sensitivity and resistance profiles across the different taxonomic of phages of the BASEL collection revealed several strong patterns that are informative about underlying molecular mechanisms. While some groups of phages like *Tevenvirinae* or *Nonagvirus* and *Seuratvirus* genera are highly homogeneous in their profiles of sensitivity or resistance (Figs 6F and 8C) – probably largely driven by conserved DNA modifications – others are more heterogeneous but never showed merely random differences. For example, the two genera of *Markadamsvirinae* differ systematically in their resistance to EcoRV (Fig 7C), likely due to vastly different numbers of recognition sites (S5 Table). Specific deviations of single phages from these patterns are sometimes readily explained like, e.g., the PifA sensitivity of T7 and JacobBurckhardt (due to their *gp1.2* allele, Figs 10C and 10D) or the EcoRV sensitivity of T3 (as the only podovirus failing in RM site avoidance regarding EcoRV; S5 Table and Fig 10C). Notably, podoviruses generally show a remarkable lack of recognition sites of type I and type II RM systems (S5 Table) which makes them phenotypically resistant [65], but this evolutionary strategy does not seem to be effective for type III RM systems (Fig 11C), possibly due to the abundance of their (shorter) recognition sequences (Fig 3B and S5 Table). Overall, we observed that phages with bigger genomes such as *Tevenvirinae*, *Vequintavirinae*, and *Markadamsvirinae* are broadly targeted by Abi systems, while smaller siphoviruses or podoviruses are either not or only sparingly targeted (Figs 4 and 6-11, see also S3 Text). Temperate phages in general seem to be highly sensitive to any kind of host immunity (Fig 11F), possibly because their evolution is less driven by selection to overcome host defenses but rather by optimizing the lysogens’ fitness, e.g., by providing additional bacterial immunity systems [8, 9].

Regarding the immunity systems themselves, we confirmed the intuitive expectation that the potency of RM systems is linked to the number of recognition sites in each phage genome unless they are masked by DNA modifications. Consequently, large phages with many recognition sites (like *Vequintavirinae*) are exceptionally sensitive to RM systems (Fig 8C), while type I RM systems perform very poorly against most phages because their long recognition sites are rare (Fig 3B and S5 Table; see also Figs 4 and 6-11). For several groups of phages like *Markadamsvirinae* and *Tevenvirinae*, the EcoRV system is the only RM system with significant impact (Figs 7C and 8C). We suggest that this is due to the blunt-cutting activity of EcoRV that is much more difficult to seal for DNA repair than the sticky ends introduced by the other tested RM systems (Fig 3B), though phage DNA ligases are known to differ in their efficiency on different kinds of DNA breaks [105].

One surprising result of our phenotyping was the remarkably poor target range of the most well-studied Abi systems of *E. coli* K-12, RexAB and PifA, that each affected only two or three phages (Figs 4 and 6-11). We do not think that this finding is an artifact of, e.g., the genetic constructs that we used, because both systems protected its host very well against the phages that they were known to target [12]: An *rIIAB* mutant of T4 was highly sensitive to RexAB (S4B Fig), and phage T7 was unable to infect a host with PifA (Fig 10C). In stark difference to RexAB and PifA, the three Abi systems of phage P2 protected each against a considerable number of phages, especially Fun/Z that inhibited all *Tevenvirinae*, *Vequintavirinae*, and *Markadamsvirinae* plus the *Felixounavirus* JohannRWettstein (Figs 4 and 6-11). Despite this remarkable potency, the molecular mechanism of Fun/Z immunity has remained unknown [50]. A common first step to unraveling the activities of an Abi system is the analysis of insensitive phage escape mutants which can be particularly insightful if they were isolated from several very different phages [106]. It seems clear that the BASEL collection as a well-assorted set of very diverse phages could be an effective tool for this purpose and also in general for a first phenotypic profiling of novel immunity systems that are commonly tested against much smaller, less well-defined sets of phages [106, 107].

Though the technicalities of our immunity phenotyping are subject to a few minor caveats (see *Materials and Methods* and S3 Text), our results demonstrate how the BASEL collection can be used to explore the biology of bacterial immunity systems and the underlying molecular mechanisms. Several important open questions remain: What is the evolutionarily or mechanistic reason for the stark differences in Abi system target range? To which extent do the different layers of bacterial immunity limit bacteriophage host range? Why are some groups of phages like the *Tevenvirinae* so strongly targeted by seemingly unrelated Abi systems and other groups of phages apparently not at all? And how well would it be possible, based on systematic phenotypic data of the BASEL collection or extensions thereof, to predict the sensitivity of newly isolated phages to diverse immunity systems just based on their genome sequences?

### Trade-offs between phage traits limit their effective host range

Our results show very clearly that seemingly advantageous phage traits such as the broad host recognition of *Vequintavirinae* or the remarkable resistance of *Tevenvirinae* to RM systems do not confer these phages a particularly large effective host range (Figs 8C and 12B). This observation can be explained by strong trade-offs between different phage fitness traits as proposed already in earlier work [108]. Unlike the hypothetical “Darwinian Demon” that would maximize all fitness traits simultaneously and dominate its ecosystem alone [109], the limits imposed by these trade-offs instead drive the adaptation of different phage groups towards specific niches that enable their long-term coexistence [108]. In marine environments, the coexistence of a few specialized, highly successful phage groups was proposed to be stable in space and time because extinction of individual phages would usually result in their replacement by relatives from the same, highly adapted group [110]. This “royal family model” could also explain why the same groups of *E. coli* phages sampled in our work have already been found again and again in previous studies that sampled diverse other environments [23–29] (Fig 2).

With our data we can at least shed light on the molecular basis of the proximal causes of the trade-offs between phage fitness traits that limit very broad effective host ranges in the selected example: *Vequintavirinae* phages are highly sensitive to various RM systems (Fig 9C), while the *Tevenvirinae* appear to be a major target of different Abi systems and have to evade a variety of those already only to infect regular *E. coli* K-12 (Figs 8C, S4B, and S4C; see also S3 Text) [9, 10, 12]. The deeper evolutionary or mechanistic basis of our observations is not clear – as an example, phages of *Nonagvirus* and *Seuratvirus* genera with genomes that are far less than half the size of *Vequintavirinae* afford full pathways for guanosine modifications that protect them reasonably well from RM systems (Figs 6E and 6F). It is not intuitive why natural selection has not resulted in similarly effective mechanisms for the *Vequintavirinae*, particularly because their promiscuity in host adsorption should bring them into very frequent contact with the diverse RM systems that form a major pillar of *E. coli* immunity [11, 49]. A good illustration for this paradox is phage PaulScherrer (Bas60), a phi92-like relative of the *Vequintavirinae* (Fig 9B), that is the only phage in the BASEL collection lysing all tested host strains but without robust plaque formation on any strain besides *E. coli* K-12 (Fig 12B).

The observed trade-offs between desired phage traits suggest that newly isolated phages might greatly profit from improvements by experimental evolution or genetic engineering to help them escape these constraints at least long enough for applications like phage therapy [14]. Such improvements seem to be principally feasible without immediate loss of fitness because, e.g., phage FriedrichZschokke (Bas41) of the *Tevenvirinae* displays only weak sensitivity to the Tin and Fun/Z Abi systems that are highly potent against its relatives (Fig 8C). However, targeted improvements of phage properties directly depend on fundamental knowledge of the molecular mechanisms underlying receptor specificity, the phages’ anti-immunity systems, and their host counterparts that are frequently unavailable [14, 41]. Future studies targeting the fundamental biology of phage-host interactions could therefore be directly helpful for such applications, and for *E. coli* phages the BASEL collection might be a suitable tool for such research.

A different way to rationalize the paradox highlighted above is that the observed trade-offs might be less pronounced in natural environments, e.g., when bacteria are starved, stressed, in biofilms, or in some other physiological state not well reflected by our laboratory experiments. Consistently, the insufficiency of regular phage phenotyping using standard laboratory growth conditions has already been recognized previously, but only limited data comparing, e.g., different host physiologies are available [16, 111]. Further studies exploring how phage infectivity, host range, and sensitivity to immunity systems change under different growth conditions would therefore be important to better understand the challenges for successful phage therapy and, more generally, the ecology and evolution of phages in natural environments.

### Concluding remarks

A recent landmark study systematically explored the genetic profile of host requirements for several *E. coli* phages and uncovered multiple unexpected and exciting twists of phage biology such as the dependence of phage N4 on high levels of the second messenger cyclic di-GMP [73]. Our current study based on the BASEL collection represents a complementary approach to explore the biology of *E. coli* phages not with elaborate genome-wide screens on the host side but rather by combining phenotypic and genomic analyses of a well-assorted set of bacteriophages. With this strategy we achieved significant advances in bacteriophage biology regarding the recognition of host receptors, the sensitivity or resistance of phages to host immunity, and how these factors come together to determine bacteriophage host range. Our work therefore establishes the BASEL collection as a powerful tool to explore new aspects of bacteriophage biology by unraveling links between phage phenotypes and their genome sequences. Furthermore, our extensive characterization of the most abundant lineages of *E. coli* phages also provides a useful field guide for teaching and outreach activities analogous to the successful SEA-PHAGES initiative [112] and the diverse student and high school projects that have enabled our study (see *Acknowledgments* and S5 Table).

## Materials and Methods

### Preparation of culture media and solutions

Lysogeny Broth (LB) was prepared by dissolving 10 g/l tryptone, 5 g/l yeast extract, and 10 g/l sodium chloride in Milli-Q H_2_O and sterilized by autoclaving. LB agar plates were prepared by supplementing LB medium with agar at 1.5% w/v before autoclaving. The M9RICH culture medium was conceived as a variant of regular M9 medium [113] supplemented with trace elements and 10% v/v LB medium prepared without NaCl to promote the growth of diverse enterobacterial strains. It was prepared from sterilized components by mixing (for 50 ml) 33.75 ml Milli-Q H_2_O, 10 ml 5x M9 salts solution, 5 ml LB medium without NaCl, 500 µl 40% w/v D-glucose solution, 100 µl 1 M MgSO_4_, and 5 µl 1 M CaCl_2_ using sterile technique. Unless indicated otherwise, all components were sterilized by filtration (0.22 µm).

Phosphate-buffered saline (PBS) was prepared as a solution containing 8 g/l NaCl, 0.2 g/l KCl, 1.44 g/l NA_2_HPO_4_ꞏ2H_2_O, and 0.24 g/l KH_2_PO_4_ with the pH adjusted to 7.4 using 10 M NaOH and sterilized by autoclaving. SM buffer was prepared as 0.1 M NaCl, 10 mM MgSO_4_, and 0.05 M Tris (pH 7.5) using sterile technique.

### Bacterial handling and culturing

*Escherichia coli* and *Salmonella* Typhimurium strains were routinely cultured in LB medium at 37°C in glass culture tubes or Erlenmeyer flasks with agitation at 170rpm. For all phenotyping assays, the bacteria were instead grown in M9RICH which supports robust growth of all strains to high cell densities. LB agar plates were routinely used as solid medium. Selection for genetic modifications or plasmid maintenance was performed with ampicillin at 50 µg/ml, kanamycin at 25 µg/ml, and zeocin at 50 µg/ml.

### Bacteriophage handling and culturing

Bacteriophages were generally cultured using the double-agar overlay method [114] with a top agar prepared as LB agar with only 0.5% w/v agar supplemented with 20 mM MgSO_4_ and 5 mM CaCl_2_. Top agar plates were incubated at 37°C and plaques were counted as soon as they were identified by visual inspection. However, the plates were always incubated for at least 24 hours to record also slow-growing plaques. We routinely used the improvements of classical phage techniques published by Kauffman and Polz [115].

High-titer stocks of bacteriophages were generated using the plate overlay method. Briefly, top agar plates were set up to grow almost confluent plaques of a given phage and then covered with 12 ml of SM buffer. After careful agitation for 24 hours at 4°C, the suspension on each plate was pipetted off and centrifuged at 8’000g for 10 minutes. Supernatants were sterilized with few drops of chloroform and stored in the dark at 4°C. For archiving, bacteriophages were stored as virocells at −80°C [116]

### Bacterial strains and strain construction

All bacterial strains used in this work are listed in S1 Table, all oligonucleotide primers in S2 Table, and all plasmids in S3 Table.

### *Escherichia coli* K-12 MG1655 ΔRM

The standard laboratory strain *E. coli* K-12 MG1655 (CGSC #6300) was engineered into a more permissive host for bacteriophage isolation by knocking out the EcoKI type I restriction-modification system (encoded by *hsdRMS*) and the McrA, Mrr, and McrBC type IV restriction systems using lambda red recombineering [117] (see Text S1 and our previous work for more technical details [118]). The resulting strain lacks all known *E. coli* K-12 restriction systems and was therefore called *E. coli* K-12 ΔRM. To enable isolation of bacteriophages with tropism for the sex pilus of the F-plasmid conjugation system, we supplied *E. coli* K-12 MG1655 ΔRM with a variant of the F-plasmid in which the *pifA* immunity gene (which might otherwise have interfered with bacteriophage isolation) had been replaced with a zeocin resistance cassette by one-step recombineering (see S1 Text for details). The strain was additionally transformed with plasmid pBR322_ΔP*tet* of reference [119] as an empty vector control for experiments with plasmid-encoded immunity systems (see below).

### *Escherichia coli* K-12 MG1655 ΔRM mutants with altered surface glycans

To test the effect of altered surface glycans of bacteriophage infection, we generated derivatives of *E. coli* K-12 MG1655 ΔRM in which genes linked to specific glycans were knocked out. The *waaC* and *waaG* genes of the LPS core biosynthesis pathway were knocked out to generate mutants displaying a deep-rough or extremely deep-rough phenotype (Fig 3A). Expression of the O16-type O-antigen of *E. coli* K-12 was restored by precisely removing the IS5 element disrupting *wbbL* [44] (Fig 3A). These mutants were generated by two-step recombineering (see details in S1 Text and a list of all strains in S1 Table). A strain specifically lacking the enterobacterial common antigen (ECA) was obtained by flipping out the kanamycin resistance cassette of the *wecB* mutant of the KEIO collection using plasmid pCP20 [117, 120]. This strain is deficient in WecB, the enzyme synthesizing UDP-ManNAcA (UDP-N-acetyl-D-mannosaminuronic acid), which is specifically required for the synthesis of the ECA but no other known glycan of *E. coli* K-12 [43].

### *Escherichia coli* K-12 BW25113 *btuB* and *tolC* knockout mutants

*btuB* and *tolC* knockout mutants isogenic with the KEIO collection were generated to use these strains lacking known bacteriophage receptors for qualitative top agar assays (see below). Details of the strain construction are provided in S1 Text.

### Other enterobacterial strains used for bacteriophage phenotyping

*E. coli* B REL606 is a commonly used laboratory strain with a parallel history of domestication to *E. coli* K-12 MG1655 [99]. *E. coli* UTI89 (ST95, O18:K1:H7) and *E. coli* CFT073 (ST73, O6:K2:H1; we used the variant with restored *rpoS* described previously [121]) are commonly used as model strains for uropathogenic *E. coli* (UPEC) and belong to phylogroup B2 [122]. *E. coli* 55989 (phylogroup B1, ST678, O104:H4) is commonly used as a model strain for enteroaggregative *E. coli* (EAEC) and closely related to the Shiga toxin-producing *E. coli* which caused the 2011 outbreak in Germany [123, 124]. *Salmonella enterica* subsp. *enterica* serovar Typhimurium strains 12023s (also known as ATCC 14028) and SL1344 are both commonly used in laboratory experiments but exhibit phylogenetic and biological differences [125].

### Plasmid construction

Plasmid vectors were generally cloned following the method of Gibson et al. (“Gibson Assembly”) [126] in which two or more linear fragments (usually PCR products) are ligated directionally guided by short 25 bp overlaps. Initially, some plasmids were also constructed using classical restriction-based molecular cloning. Briefly, a PCR-amplified insert and the vector backbone were each cut with appropriate restriction enzymes (New England Biolabs). After dephosphorylation of the backbone (using FastAP dephosphorylase; Thermo Scientific), insert and backbone were ligated using T4 DNA ligase (Thermo Scientific). Local editing of plasmid sequences was performed by PCR with partially overlapping primers as described by Liu and Naismith (127). *E. coli* strain EC100 *pir(+)* was used as host for all clonings. The successful construction of every plasmid was confirmed by Sanger Sequencing. A list of all plasmids used in this study is found in S3 Table and their construction is summarized in S4 Table. The sequences of all oligonucleotide primers used in this study are listed in S2 Table.

For the series of plasmids encoding the eleven different bacterial immunity systems studied in this work (see Fig 3B), we used the EcoRI and EcoRV constructs of Pleška, Qian et al. [119] as templates and, consequently, the corresponding empty vector pBR322_ΔP*tet* as general cloning backbone and experimental control. The different immunity systems were generally cloned together with their own transcriptional promoter region. However, the *rexAB* genes are transcribed together with the cI repressor of the lambda prophage [7]. We therefore cloned them directly downstream of the P*tet* promoter of the pBR322 backbone and obtained a functional construct (validated in S4B Fig). The EcoKI, EcoRI, EcoRV, and EcoP1_I RM systems have been studied intensively in previous work [47–49]. Besides the type IA RM system EcoKI we also cloned EcoCFT_I that is nearly identical to the well-characterized type IB RM system EcoAI but encoded in the genome of *E. coli* CFT073 that was available to us [47, 128]. Similarly, the EcoCFT_II system was identified as a type III RM system in the *E. coli* CFT073 genome using REBASE [49, 128]. Due to problems with toxicity, some immunity systems were cloned not into pBR322_ΔP*tet* but rather into a similar plasmid carrying a low-copy SC101 origin of replication (pAH186SC101e [121], see S3 and S4 Tables as well as the considerations in S3 Text). Since we failed to obtain any functional construct for the ectopic expression of *pifA* (as evidenced by lack of immunity against T7 infection), we instead replaced the F(*pifA::zeoR*) plasmid in *E. coli* K-12 MG1655 ΔRM with a wildtype F plasmid that had merely been tagged with kanamycin resistance and encodes a functional *pifA* (pAH200e).

### Bacteriophage isolation

#### Basic procedure

Bacteriophages were isolated from various different samples between July 2019 and November 2020 using *E. coli* K-12 MG1655 ΔRM as the host (see S5 Table for details) using a protocol similar to common procedures in the field [16]. Phage isolation was generally performed without an enrichment step to avoid biasing the isolation towards fast-growing phages (but see below).

For aqueous samples we directly used 50 ml, while samples with major solid components (like soil or compost) were agitated overnight at 4°C in 50 ml of PBS to release viral particles. Subsequently, all samples were centrifuged at 8’000 g for 15 minutes to spin down particles larger than viruses. The supernatants were sterilized treated with 5% v/v chloroform which safely inactivates any bacteria as well as enveloped viruses but will generally leave most *Caudovirales* intact [16]. Subsequently, viral particles were precipitated by adding 1 ml of a 2 M ZnCl_2_ solution per 50 ml of sample, mixing shortly by inversion, and incubating the suspension at 37°C without agitation for 15 minutes [129]. After precipitation, the samples were centrifuged again at 8’000 g for 15 minutes and the supernatant was discarded. The pellets were carefully resuspended in each 500 µl of SM buffer by agitation at 4°C for 15 minutes. Subsequently, the suspensions were cleared quickly using a tabletop spinner and mixed with 500 µl of bacterial overnight culture (resuspended in fresh LB medium to induce resuscitation). After incubation at room temperature for 15 minutes to promote phage adsorption, each mixture was added to 9 ml of pre-warmed top agar and poured onto a pre-warmed square LB agar plate (ca. 12 cm x 12 cm). After solidification, the plates were incubated at 37°C for up to 24 hours.

#### Isolation of bacteriophage clones

Bacteriophages were visible as plaques forming in the dense bacterial growth of the top agar. For isolation of bacteriophage clones, they were picked from clearly separated plaques of diverse morphologies with sterile toothpicks and propagated at least three times via single plaques on top agars of the isolation host strain *E. coli* K-12 MG1655 ΔRM. To avoid isolating temperate phages or phages that are poorly adapted to *E. coli* hosts, we only picked clear plaques (indicative of lytic phages) and discarded isolates that showed poor plaque formation [16].

#### Isolation of *Autographiviridae* using enrichment cultures

The direct plating procedure outlined above never resulted in the isolation of phages belonging to the *Autographiviridae* like iconic T phages T3 and T7. Given that these phages are known for fast replication and high burst sizes [130], we therefore performed a series of enrichment culture isolation experiments to obtain phages forming the characteristically large, fast-growing plaques of *Autographiviridae*. For this purpose, we prepared M9 medium using a 5x M9 salts solution and chloroform-sterilized sewage plant inflow instead of water (i.e., containing ca. 40 ml of sewage plant inflow per 50 ml of medium) and supplemented it with 0.4% w/v D-glucose as carbon source. 50 ml cultures were set up by inoculating these media with each 1 ml of an *E. coli* K-12 MG1655 ΔRM overnight culture and agitated the cultures at 37°C for 24 hours. Subsequently, the cultures were centrifuged at 8’000 g for 15 minutes and each 50 µl of supernatant was plated with the *E. coli* K-12 MG1655 ΔRM isolation strain in a top agar on one square LB agar plate. After incubation at 37°C for three or four hours, the first *Autographiviridae* plaques characteristically appeared (before most other plaques) and were picked and propagated as described above. Using this procedure, we isolated four different new *Autographiviridae* isolates (see S5 Table and Fig 10B).

#### Composition of the BASEL collection

Bacteriophages were mostly isolated and characterized from randomly picked plaques in direct selection experiments, but we later adjusted the procedure to specifically isolate *Autographiviridae* which were the only major group of phages previously shown to infect *E. coli* K-12 missing from our collection (see above). After every set of 10-20 phages that had been isolated, we performed whole-genome sequencing and preliminary phylogenetic analyses to keep an overview of the growing collection (see below). In total, more than 120 different bacteriophages were sequenced and analyzed of which we selected 66 tailed, lytic phage isolates to compose the BASEL collection (see Fig 2 and S5 Table for details). Phages closely related to other isolates were deliberately excluded unless they displayed obvious phenotypic differences such as, e.g., a different host receptor. In addition to the 66 newly isolated bacteriophages, ten classical model phages were included for genomic and phenotypic characterization, and we view these phages as an accessory part of the BASEL collection. These ten phages were six of the seven T phages (excluding T1 because it is a notorious laboratory contaminant [32]), phage N4, and obligately lytic mutants of the three well-studied temperate phages lambda, P1, and P2 [5, 7, 33, 34] (Fig 2; see also S5 Table). To generate the T3(K12) chimera, the 3’ end of *gp17* lateral tail fiber gene of phage T7 was cloned into low-copy plasmid pUA139 with flanking regions exhibiting high sequence similarity to the phage T3 genome (generating pUA139_T7(*gp17*), see S4 Table). Phage T3 was grown on *E. coli* B REL606 transformed with this plasmid and then plated on *E. coli* K-12 MG1655 ΔRM to isolate recombinant clones. Successful exchange of the parental T3 *gp17* allele with the variant of phage T7 was confirmed by Sanger Sequencing.

#### Qualitative top agar assays

The lysis host range of isolated bacteriophages on different enterobacterial hosts and their ability to infect strains of a set of KEIO collection mutants lacking each one surface protein (or isogenic mutants that were generated in this study, see S1 Table and S1 Text) were tested by qualitative top agar assays. For this purpose, top agars were prepared for each bacterial strain on LB agar plates. For by overlaying them with top agar supplemented with a suitable bacterial inoculum. For regular round Petri dishes (ca. 9.4 cm diameter) we used 3 ml of top agar supplemented with 100 µl of bacterial overnight culture, while for larger square Petri dishes (ca. 12 cm x 12 cm) we used 9 ml of top agar supplemented with 200 µl of overnight culture. After solidification, each 2.5 µl of undiluted high-titer stocks of all tested bacteriophages (>10^9^ pfu/ml) were spotted onto the top agar plates and dried into the top agar before incubation at 37°C for at least 24 hours. If lysis zones on any enterobacterial host besides our *E. coli* K-12 ΔRM reference strain were observed, we quantified phage infectivity in efficiency of plating assays (see below). Whenever a phage failed to show lysis on a mutant strain lacking a well-known phage receptor, we interpreted this result as indicating that the phage infecting depends on this factor as host receptor.

#### Efficiency of Plating assays

The infectivity of a given bacteriophage on a given host was quantified by determining the efficiency of plating (EOP), i.e., by quantitatively comparing its plaque formation on a certain experimental host to plaque formation on reference strain *E. coli* K-12 MG1655 ΔRM carrying F(*pifA::zeoR*) and pBR322_ΔP*tet* [131]. Experimental host strains we identical to the reference strain with the difference that they either carried plasmids encoding a certain bacterial immunity system (S3 Table) or had a chromosomal modification changing surface glycan expression (Fig 3A and S1 Table).

For quantitative phenotyping, top agars were prepared for each bacterial strain on LB agar plates by overlaying them with top agar (LB agar containing only 0.5% agar and additionally 20 mM MgSO_4_ as well as 5 mM CaCl_2_; stored at 60°C) supplemented with a suitable bacterial inoculum. For regular round Petri dishes (ca. 9.4 cm diameter) we used 3 ml of top agar supplemented with 100 µl of bacterial overnight culture, while for larger square Petri dishes (ca. 12 cm x 12 cm) we used 9 ml of top agar supplemented with 200 µl of overnight culture. While the top agars were solidifying, serial dilutions of bacteriophage stocks (previously grown on *E. coli* K-12 MG1655 ΔRM to erase any EcoKI methylation) were prepared in sterile phosphate-buffered saline (PBS). Subsequently, each 2.5 µl of all serial dilutions were spotted on all top agar plates and dried into the top agar before incubation at 37°C for at least 24 hours. Plaque formation was recorded repeatedly throughout this time (starting after 3 hours of incubation for fast-growing phages). The EOP of a given phage on a certain host was determined by calculating the ratio of plaques obtained on this host over the number of plaques obtained on the reference strain *E. coli* K-12 MG1655 ΔRM carrying F(*pifA::zeoR*) and pBR322_ΔP*tet* [131].

When no plaque formation could be unambiguously recorded by visual inspection, the EOP was determined to be below detection limit even if the top agar showed lysis from without (i.e., lysis zones caused by bacterial cell death without phage infection at an efficiency high enough to form plaques [98]). However, for all non-K12 strains of *E. coli* as well as *Salmonella* Typhimurium we determined the lysis host range (i.e., the range of hosts on which lysis zones were observed) besides the numerical determination of EOP (see above). Occasionally, we found that certain phage / host pairs were on the edge between merely strong lysis from without and very poor plaque formation. Whenever in doubt, we recorded the result conservatively as an EOP below detection limit (e.g., for phage FriedrichMiescher on *E. coli* 55989, *Enquatrovirus* phages and EmilHeitz on *E. coli* UTI89, and all *Vequintavirinae* on the host expressing the Fun/Z Abi system).

#### Bacteriophage genome sequencing and assembly

Genomic DNA of bacteriophages was prepared from high-titer stocks using the Norgen Biotek Phage DNA Isolation Kit according to the manufacturer’s guidelines and sequenced at the Microbial Genome Sequencing Center (MiGS) using the Illumina NextSeq 550 platform. Trimmed sequencing reads were assembled using the Geneious Assembler implemented in Geneious Prime 2021.0.1 with a coverage of typically 50-100x (S5 Table). Usually, circular contigs (indicating a complete assembly due to the fusion of characteristically repeated sequence at the genome ends [132]) were easily obtained using the “Medium Sensitivity / Fast” setting. Consistently incomplete assemblies or local ambiguities were solved by PCR amplification using the high-fidelity polymerase Phusion (NEB) followed by Sanger Sequencing. For annotation and further analyses, sequences were linearized with the 5’ end set either to the first position of the small terminase subunit gene or the first position of the operon containing the small terminase subunit gene.

#### Bacteriophage genome annotation

A preliminary, automated annotation of the genes in all genomes was generated using MultiPhate [133] and then manually refined. For this purpose, whole-genome alignments of all new isolates within a given group of phages and well-studied and / or well-annotated references were generated using progressiveMauve [134] implemented in Geneious 2021.0.1 and used to inform the annotation based on identified orthologs. *Bona fide* protein-coding genes without clear functional annotation were translated and analyzed using the blastp tool on the NCBI server (https://blast.ncbi.nlm.nih.gov/Blast.cgi), the InterPro protein domain signature database [135], as well as the Phyre2 fold recognition server [136; http://www.sbg.bio.ic.ac.uk/~phyre2/html/page.cgi?id=index]. tRNA genes were predicted using tRNA-ScanSE in the [137; http://lowelab.ucsc.edu/tRNAscan-SE/] and spanins were annotated with help of the SpaninDataBase tool [138]. While endolysin genes were easily recognized by homology to lysozyme-like proteins and other peptidoglycan hydrolases, holins were more difficult to identify when no holins were annotated in closely related bacteriophage genomes. In these cases, we analyzed all small proteins (<250 amino acids) for the presence of transmembrane helices – a prerequisite for the functionality of known holins [139] – and studied their possible relationships to previously described holins using blastp and InterPro. In most but not all cases, *bona fide* holins encoded close to endolysin and / or spanin genes could be identified. The annotation of our genomes in a comparative genomics setup made it easily possible to identify the boundaries of introns associated with putative homing endonucleases and to precisely identify inteins (see also S5 Table). The annotated genome sequences of all 66 newly isolated phages as well as T3(K12) have been submitted to the NCBI GenBank database under accession numbers listed in S5 Table.

#### Bacteriophage naming and taxonomy

Newly isolated bacteriophages were named according to rules and conventions in the field and classified in line with the rules of the International Committee on the Taxonomy of Viruses (ICTV) [79, 140] (S5 Table). As a first step of taxonomic classification, phages were roughly sorted by family and genus based on whole-genome blastn searches against the non-redundant nucleotide collection database [https://blast.ncbi.nlm.nih.gov/Blast.cgi; see also reference 141]. For each such broad taxonomic group, we selected a set of reference sequences from NCBI GenBank that would cover the diversity of this group and that always included all members which had already previously been studied intensively. Subsequently, we generated whole-genome alignments of these sets of sequences using progressiveMauve implemented in Geneious Prime 2021.0.1 [134]. These alignments or merely regions encoding highly conserved marker genes such as the terminase subunits, portal protein gene, major capsid protein gene, or other suitable loci were then used to generate Maximum-Likelihood phylogenies to unravel the evolutionary relationships between the different bacteriophage isolates and database references (see S2 Text for details). Care was taken to avoid genome regions that are recognizably infested with homing endonucleases or that showed obvious signs of having recently moved by horizontal gene transfer (e.g., an abrupt shift in the local sequence identity to the different other genomes in the alignment). Clusters of phage isolates observed in these phylogenies generally correlated very well with established taxonomy as inferred from NCBI taxonomy (https://www.ncbi.nlm.nih.gov/taxonomy), the ICTV [https://talk.ictvonline.org/taxonomy/; reference 142], and other reports in the literature [30]. To classify our phage isolates on the species level, we generated pairwise whole-genome alignments using progressiveMauve implemented in Geneious Prime 2021.0.1 [134] with genomes of related phages as identified in our phylogenies. From these alignments, the nucleotide sequence identity was determined as the query coverage multiplied by the identity of aligned segments [24]. When the genomes were largely syntenic and showed >95% nucleotide sequence identity, we classified our isolates as the same species as their close relative [141], at least if this phage has been assigned to a species by the ICTV [142].

Most phages were named in honor of scientists and other historically relevant personalities with links to the city of Basel, Switzerland, where the majority of phages had been isolated. Despite efforts to name many phages after female personalities, a considerable gender imbalance remains. This imbalance is a consequence of the biases in how science and history have been made and recorded before the 20^th^ century.

#### Sequence alignments and phylogenetic analyses

The NCBI GenBank accession numbers of all previously published genomes used in this study are listed in S6 Table. Sequence alignments of different sets of homologous genes were generated using MAFFT v7.450 implemented in Geneious Prime 2021.0.1 [143]. Whenever required, poor or missing annotations in bacteriophage genomes downloaded from NCBI GenBank were supplemented using the ORF finder tool of Geneious Prime 2021.0.1 guided by orthologous sequence parts of related genomes. Alignments were set up using default settings typically with the fast FFT-NS-2 algorithm and 200PAM/k=2 or BLOSUM62 scoring matrices for nucleotide and amino acid sequences, respectively. Subsequently, alignments were curated manually to improve poorly aligned sequence stretches and to mask non-homologous parts.

For phylogenetic analyses, sequence alignments of orthologous stretches from different genomes (genes, proteins, or whole genome) were used to calculate Maximum-Likelihood phylogenies with PhyML 3.3.20180621 implemented in Geneious Prime 2021.0.1 [144]. Phylogenies were calculated with the HYK85 substitution model for nucleotide sequences and with the LG substitution model for amino acid sequences. For the inference of phylogenetic relationships between phage genomes, we sometimes used curated whole-genome alignments, but the infestation with homing endonucleases and the associated gene conversion made this approach impossible for all larger genomes. Instead, we typically used curated sequence alignments of several conserved core genes (on nucleotide or amino acid level, depending on the distance between the genomes) as the basis for Maximum-Likelihood phylogenies. The detailed procedures for each phylogeny shown in this article are described in S2 Text.

#### LPS structures of enterobacterial strains

*Escherichia* K-12 MG1655 (serogroup A) codes for a K12-type LPS core and an O16-type O-antigen, but functional expression of the O-antigen was lost during laboratory adaptation due to inactivation of *wbbL* by an IS5 insertion so that only a single terminal GlcNAc is attached to the LPS core [44, 72]. *E. coli* strains UTI89 (serogroup B2), CFT073 (serogroup B2), and 55989 (serogroup B1) express O-antigens of the O18, O6, and O104 types, respectively [122, 123]. Their core LPS types were determined to be R1 (UTI89 and CFT073) and R3 (55989) by BLAST searches with diagnostic marker genes similar to the PCR-based approached described previously [72]. For the illustration in Fig 12A, the structures of LPS cores and linked O-antigen polysaccharides were drawn as described in the literature with typical O-antigen chain lengths of 15 - 25 repeat units [72, 103, 145]. For the O18-type of O-antigen polysaccharides, different subtypes with slight differences in the repeat unit have been described [103], but to the best of our knowledge it has remained elusive which subtype is expressed by *E. coli* UTI89. In Fig 12A we therefore chose an O18A type as exemplary O-antigen polysaccharide for *E. coli* UTI89. Similar to the laboratory adaptation of *E. coli* K-12 MG1655, strains of the *E. coli* B lineage like *E. coli* B REL606 lack O-antigen expression due to an IS1 insertion in *wbbD* but have an additional truncation in their R1-type LPS core due to a second IS1 insertion in *waaT* that leaves only two glucoses in the outer core (Fig 12A) [67, 99]. *Salmonella enterica* subsp. *enterica* serovar Typhimurium strains 12023s (also known as ATCC 14028) and SL1344 share the common *S.* Typhimurium LPS core and *Salmonella* serogroup B / O4 O-antigen [42, 146] (Fig 12A).

#### Bacterial genome sequencing and analyses to identify host receptors

While top agar assays with *E. coli* mutants carrying defined deletions in genes coding for different surface proteins or genes involved in the LPS core biosynthesis readily identified the host receptor of most bacteriophages (see above), few small siphoviruses of the *Drexlerviridae* family and the *Dhillonvirus*, *Nonagvirus*, and *Seuratvirus* genera of the *Siphoviridae* family could not be assigned to any surface protein as secondary receptor. This was the case for phages AugustePiccard (Bas01) and JeanPiccard (Bas02) of *Drexlerviridae*, TheodorHerzl (Bas14) and Oekolampad (Bas18) of *Dhillonvirus*, ChristophMerian (Bas19) and FritzHoffmann (Bas20) of *Nonagvirus*, and VogelGryff (Bas25) of *Seuratvirus*. However, as siphoviruses infecting Gram-negative bacteria it seemed highly likely that these phages would use an outer membrane porin as their secondary receptor [40]. We therefore isolated resistant mutants of *E. coli* K-12 BW25113 by plating bacteria on LB agar plates which had been densely covered with high-titer lysates of each of these bacteriophages. While verifying that the resistance of isolated bacterial clones was specific to the phage on which they had been isolated, we found that many clones were fully resistant specifically but without exception to all these small siphoviruses with yet no known host receptor, suggesting that they all targeted the same receptor.

For multiple of these phage-resistant clones, genomic DNA was prepared using the GenElute Bacterial Genomic DNA Kit (Sigma-Aldrich) according to the manufacturer’s guidelines and sequenced at the Microbial Genome Sequencing Center (MiGS) using the Illumina NextSeq 550 platform. Sequencing reads were assembled to the *E. coli* K-12 MG1655 reference genome (NCBI GenBank accession U000096.3) using Geneious Prime 2021.0.1 with a coverage of typically 100-200x to identify the mutations underlying their phage resistance. These analyses uncovered a diversity of in-frame deletions in *lptD* for the different mutant clones (Fig 5).

#### Quantification and statistical analysis

Quantitative data sets were analyzed by calculating mean and standard deviation of at least three independent biological replicates for each experiment. Detailed information about replicates and statistical analyses for each experiment is provided in the figure legends.

## Acknowledgments

The authors are grateful to Prof. Urs Jenal, Prof. Marek Basler, Prof. Martin Loessner, and Dr. Julie Sollier for valuable input and critical reading of the manuscript. The authors are deeply indebted to Prof. Urs Jenal and Prof. Christoph Dehio for sharing multiple different *E. coli* strains and plasmids (S1 and S3 Tables). Prof. Calin Guet is acknowledged for sharing plasmids (S3 Table). The authors thank high school students participating in the Biozentrum Summer Science Academy 2019, Julia Harms, and Dr. Thomas Schubert for assistance with the isolation of multiple bacteriophages described in this study (S5 Table). *Salmonella* Typhimurium strains 12023s and SL1344 were obtained from Prof. Dirk Bumann and Prof. Mederic Diard, respectively, while *E. coli* B REL606 was obtained from Dr. Jenna Gallie. Phage P2*vir* was generously shared by Dr. Nicolas Wenner and Prof. Jay Hinton. The Coli Genetic Stock Center (CGSC) and the Deutsche Sammlung von Mikroorganismen und Zellkulturen (DSMZ) are acknowledged for sharing different strains of *E. coli* or different bacteriophages, respectively (S1 and S5 Tables). Dr. Christian Beisel and Dr. Geoffrey Fucile are acknowledged for advice regarding DNA sequencing. The authors are grateful to ARA Basel, ARA Canius (Lenzerheide), and ARA Limeco (Dietikon) for providing samples of sewage plant inflow and to Universitätsspital Basel for hospital sewage.

## Data reporting

All data generated or analyzed during this study are included in this published article.

## Accession numbers

The annotated genome sequences of all 66 newly isolated phages as well as T3(K12) have been submitted to the NCBI GenBank database under accession numbers listed in S5 Table.

## Supporting Information Captions

**S1 Table. List of all bacterial strains used in this study.**

The abbreviations in the selection column indicate the drug and its concentration that were used. Amp = ampicillin, Cam = chloramphenicol, Kan = kanamycin, Zeo = zeocin; 25 / 50 / 100 refer to 25 µg/ml, 50 µg/ml, and 100 = 100 µg/ml, respectively. The following mutants of the KEIO collection were used for qualitative top agar assays but are not included in the strain list because no phage showed any growth phenotype on them: *ompW::kanR*, *phoE::kanR*, *flgG::kanR*, *fepA::kanR*, *hofQ::kanR*, *cirA::kanR*, *fhuE::kanR*, *fiu::kanR*, *ompN::kanR*, *pgaA::kanR*, *chiP::kanR*, *ompL::kanR*, *yddB::kanR*, *fecA::kanR*, *uidC::kanR*, *nanC::kanR*, *yfaZ::kanR*, *bglH::kanR*, *bcsC::kanR*, *cusC::kanR*, *gfcE::kanR*, *mdtP::kanR*, *ompG::kanR*, *ompX::kanR*, *yfeN::kanR*, *csgF::kanR*, *wza::kanR*, *flu::kanR*, *nmpC::kanR*, *eaeH::kanR*, *ydiY::kanR*, *yiaT::kanR*, *yaiO::kanR*, *mdtQ::kanR*, *pgaB::kanR*, *mipA::kanR*, *pldA::kanR*, *yzcX::kanR*, *ydeT::kanR*, *blc::kanR*, *gspD::kanR*, *yjgL::kanR*

**S2 Table. List of all oligonucleotide primers used in this study.**

**S3 Table. List of all plasmids used in this study.**

The abbreviations in the selection column indicate the drug and its concentration that were used. Amp = ampicillin, Cam = chloramphenicol, Kan = kanamycin, Zeo = zeocin; 25 / 50 / 100 refer to 25 µg/ml, 50 µg/ml, and 100 = 100 µg/ml, respectively.

**S4 Table. Construction of all plasmids generated in this study.**

**S5 Table. List of all phages used in this study.**

The table lists all details regarding the taxonomic classification, isolation / source, host receptors, and genomic features of all phages used in this study. As *Caudovirales*, all these phages are classified as *Viruses* / *Duplodnaviria* / *Heunggongvirae* / *Uroviricota* / *Caudoviricetes* / *Caudovirales*, so only the classification on the level of family and below as defined by the ICTV [https://talk.ictvonline.org/taxonomy/; reference 142] is listed. For the reference phages, genome sizes and estimation of RM system recognition sites are based on the reference genomes in NCBI GenBank as indicated. Note that for phage T5 the reference genome includes the 10’219bp terminal repeat twice, unlike all other *Markadamsvirinae* genomes that we listed. The P2*vir* phage had been isolated as a spontaneous mutant of a P2 *cox3* lysogen (supposedly unable to excise [148]) and was analyzed by whole-genome sequencing (as described in *Materials and Methods*), revealing few single-nucleotide differences to the reference genome, the expected *cox3* mutation, and a one-basepair insertion in the lysogeny repressor gene *C* that results in a premature stop codon. Recognition sites of RM systems were identified using Geneious Prime 2021.0.1. The numbers include the two orientations of palindromic recognition sequences separately and do not account for the fact that the counts are not fully comparable, e.g., because type III RM systems require two unmodified recognition sites in head-to-head orientation for cleavage [49]. Given the high number of type III RM recognition sites in all genomes because of their short length (5 nt in case of both EcoCFT_II and EcoP1_I, see Fig 3B), this limitation is largely theoretical.

**S6 Table. List of all phage genomes used in the *in silico* analyses.**

**S1 Text. Construction of bacterial mutant strains.**

**S2 Text. Generation of the Maximum-Likelihood phylogenies shown in this article.**

**S3 Text. Technical considerations regarding the composition of the BASEL collection and the phenotyping of bacterial defense systems.**

**S1 Figure. Supplemental data for** Figs 4 and 5.

**(A)** Maximum-Likelihood phylogeny of *Drexlerviridae* and the *Dhillonvirus* genus of *Siphoviridae* based on several core genes with bootstrap support of branches shown if > 70/100. It is clearly apparent that *Drexlerviridae* are split into two major clades, one formed by *Braunvirinae* and *Rogunavirinae* and another one formed by *Tempevirinae*, *Tunavirinae*, plus a few other groups. Given that the phylogenies strongly agree on all major branches, the root of the *Drexlerviridae* phylogeny shown in Fig 4B was placed between these two major clades. **(B)** The locus encoding lateral tail fibers was analyzed in a sequence alignment of the thirteen *Drexlerviridae* phage genomes of the BASEL collection (see *Materials and Methods*). It is clearly visible that the upstream and downstream regions (encoding genes involved in recombination as well as primase / helicase proteins for genome replication) are highly conserved and fully syntenic, with exception of small insertions in a few sequences. Conversely, only the most 5’ end of the largest lateral tail fiber protein gene is very similar among all analyzed genomes (green circle), while the rest shows neither synteny nor clear homology across all genomes. **(C)** The *bona fide* RBP loci downstream of the *gpJ* homolog are shown for all small siphoviruses (*Drexlerviridae* and *Siphoviridae* of *Dhillonvirus*, *Nonagvirus*, and *Seuratvirus* genera) where we had experimentally determined the terminal receptor (together with selected representatives where previous work had determined the receptor specificity). **(D)** The *bona fide* RBP locus of *E. coli* phage RTP was aligned to the homologous locus of phage AugustePiccard (Bas01) as described in *Materials and Methods*. For the region comprising *rtp44* and *rtp45* of phage RTP, the pairwise identity of the two nucleotide sequences is ca. 93%.

**S2 Figure. Supplemental data for** Fig 6.

**(A)** The locus encoding lateral tail fibers was analyzed in sequence alignments of the five *Dhillonvirus* phage genomes of the BASEL collection (top) and the seven *Nonagvirus* + *Seuratvirus* phage genomes of the BASEL collection (bottom) as described in *Materials and Methods*. In both cases two clear dips in overall sequence similarity are obvious, once at the *bona fide* RBP locus and then at the lateral tail fiber locus downstream of the far 5’ end of its first gene. **(B)** Schematic comparison of representative *bona fide* RBP loci as shown in S1C Fig to the corresponding allele of *E. coli* phages JenK1, HdK1, and HdsG1 that does not match clearly match any of them.

**S3 Figure. Supplemental data for** Fig 7.

**(A)** The locus encoding lateral tail fibers was analyzed in a sequence alignment of the *Demerecviridae: Markadamsvirinae* phage genomes of the BASEL collection as described in *Materials and Methods*. Sequence identity is high upstream and downstream of the lateral tail fiber locus (with exception of presence / absence of a few putative homing endonucleases) but drops considerably at the lateral tail fiber genes. Note that, as described previously, the lateral tail fibers can either be composed of a single large polypeptide or by two (or more) separate proteins [51, 60]. The same diversity in architecture of lateral tail fibers can also be seen at the corresponding loci of small siphoviruses (S1B and S2A Figs). **(B)** The illustration shows a phylogeny of the RBPs of all *Markadamsvirinae* phages shown in Fig 7B. Briefly, the RBP genes of all genomes (invariably encoded directly upstream of the terminase genes) were translated, aligned, and then used to generate a phylogeny as described in *Materials and Methods*. Three clear clusters emerge, one including all phages known to bind BtuB (left), one including all phages known to bind FepA (top right), and one including all phages known to bind FhuA (bottom right). Based on similar analyses by others [62], we conclude that the position of RBPs in these clusters is predictive of terminal receptor specificity of the phages encoding them.

**S4 Figure. Supplemental data for** Fig 8.

**(A)** Top agar assays with different surface protein mutants of the KEIO collection in comparison to the ancestral *E. coli* K-12 BW25113 strain were performed with serial tenfold dilutions of all *Tevenvirinae* phages used in this study (undiluted high-titer stocks at the bottom and increasingly diluted samples towards the top). The phages show strongly or totally abolished growth on each one of the mutants which identifies the primary receptor of each phage (also indicated by the color code highlighted on the right). **(B)** Top agar assays of reference strain *E. coli* K-12 ΔRM carrying empty vector pBR322_ΔP*tet* or pAH213_rexAB were performed with serial tenfold dilutions of phage T4 wildtype, a T4 variant encoding an apparently hypomorphic *rIIAB* fusion (see (C)), and phage T5 (as control). The *rIIAB* mutant of phage T4 is unable to form plaques on the *rexAB*-expressing host and shows only “lysis from without” [98], while the T4 wildtype and phage T5 are not affected. **(D)** A T4 phage mutant was erroneously obtained from a culture collection instead of the wildtype and encoded a peculiar *rII* allele with fusion of the *rIIA* and *rIIB* open reading frames (shown as orange arrows; genome sequence determined by whole-genome sequencing). Since such a mutant seems unlikely to arise spontaneously during shipping, we find it likely that this phage strain is related to the *rIIAB* fusion mutants employed for discovery of the triplet nature of the genetic code that were once commonly used [concisely reviewed in reference 149, 150]. Notably, position and size of the deletion fusing *rIIA* and *rIIB* are indistinguishable from the sketch drawn by Benzer and Champe for the *rIIAB* fusion mutant *r1589* which was used in the aforementioned work [151]. Unlike T4 wildtype, the *rIIAB* fusion mutant is susceptible to *rexAB* when ectopically expressed from a plasmid vector (see (C)) and therefore validates functionality of the *rexAB* construct.

**S5 Figure. Supplemental data for** Fig 9.

The illustration shows a sequence alignment of the locus encoding lateral tail fiber genes in phage rV5 and new isolates DrSchubert, AlexBoehm, and JeffSchatz that broadly cover the phylogenetic range of this genus (Fig 9B). It extends from the tail tape measure protein of phage rV5 (*gp49*) to the last large lateral tail fiber gene (*gp27*) [70, 71]. Note that most genes are highly conserved including the lateral tail fiber component with sugar deacetylase domain (compare Fig 9D; around position 8’000 in this alignment) or the short tail fiber locus (compare Fig 9E). Only the three largest lateral tail fiber genes show considerable allelic variation that is very strong for two of them (positions 10’000-14’000 and 16’000-19’000) and moderate for another one (positions 29’000 to 34’000).

