## Supplementary material for "Systematic exploration of *Escherichia coli* phage-host interactions with the BASEL phage collection": Tables S1, S2, and S3

### S1 Table. List of all bacterial strains used in this study

| Strain | Genotype | relevant plasmid | Selection | Source/Description |
|---|---|---|---|---|
| AH-E02-148 | *Escherichia coli* K-12 MG1655: F⁻ λ⁻ *ilvG*⁻ *rfb-50 rph-1* | none | none | *E. coli* K-12 laboratory wildtype strain; obtained from the Coli Genetic Stock Center (CGSC #6300) via Prof. Urs Jenal |
| AH-E01-047 | *Escherichia coli* K-12 BW25113: F⁻ Δ(*araD-araB*)567, Δ*lacZ4787(::rrnB-3)*, λ⁻, *rph-1*, Δ(*rhaD-rhaB*)568, *hsdR514* | none | none | our laboratory collection [1] |
| AH-E02-160 | *E. coli* K-12 MG1655 Δ*mrr-hsdRMS-mcrBC* | pWRG99 | Amp100 | this study |
| AH-E03-200 | *E. coli* K-12 MG1655 Δ*mrr-hsdRMS-mcrBC* Δ*mcrA* = ΔRM | none | none | this study; strain lacking all known restriction systems of *E. coli* K-12 |
| AH-E03-217 | *E. coli* K-12 MG1655 ΔRM | pBR322_ΔP*tet* F(*pifA::zeoR*) | Amp50 Zeo50 | this study |
| AH-E06-427 | *E. coli* K-12 MG1655 ΔRM | pBR322_ΔP*tet* pAH200e | Amp50 Kan25 | this study; pAH200e (F-plasmid tagged with kanamycin resistance at *tn1000* obtained from Prof. Christoph Dehio) |
| AH-E07-554 | *E. coli* K-12 W1872 | F | none | *E. coli* K-12 strain carrying a wildtype F-plasmid |
| AH-E03-233 | *E. coli* K-12 MG1655 ΔRM *waaC::kanR* | none | Kan25 | this study |
| AH-E03-235 | *E. coli* K-12 MG1655 ΔRM *waaG::kanR* | none | Kan25 | this study |
| AH-E03-243 | *E. coli* K-12 MG1655 ΔRM *wbbL(+)* | none | none | this study |
| AH-E04-292 | *E. coli* K-12 BW25113 *wecB::FRT* | none | none | obtained from Prof. Urs Jenal |
| AH-E04-321 | *E. coli* K-12 BW25113 *btuB::kanR* | none | Kan25 | this study |
| MBu-E01-044 | *E. coli* K-12 BW25113 *tolC::kanR* | none | Kan25 | this study |
| MBu-E01-007 | *E. coli* K-12 BW25113 *fhuA::kanR* | none | Kan25 | obtained from Prof. Urs Jenal (KEIO collection [2]) |
| AH-E05-396 | *E. coli* K-12 BW25113 *yncD::kanR* | none | Kan25 | obtained from Prof. Urs Jenal (KEIO collection [2]) |
| MBu-E01-018 | *E. coli* K-12 BW25113 *lamB::kanR* | none | Kan25 | obtained from Prof. Urs Jenal (KEIO collection [2]) |
| MBu-E01-015 | *E. coli* K-12 BW25113 *tsx::kanR* | none | Kan25 | obtained from Prof. Urs Jenal (KEIO collection [2]) |
| MBu-E01-016 | *E. coli* K-12 BW25113 *fadL::kanR* | none | Kan25 | obtained from Prof. Urs Jenal (KEIO collection [2]) |
| MBu-E01-011 | *E. coli* K-12 BW25113 *ompA::kanR* | none | Kan25 | obtained from Prof. Urs Jenal (KEIO collection [2]) |
| MBu-E01-012 | *E. coli* K-12 BW25113 *ompC::kanR* | none | Kan25 | obtained from Prof. Urs Jenal (KEIO collection [2]) |
| MBu-E01-013 | *E. coli* K-12 BW25113 *ompF::kanR* | none | Kan25 | obtained from Prof. Urs Jenal (KEIO collection [2]) |
| AH-E07-546 | *E. coli* K-12 BW25113 *lptD*_Δ(L394-V396)::Y | none | none | this study; spontaneous mutant resistant to LptD-targeting siphoviruses |
| AH-E07-545 | *E. coli* K-12 BW25113 *lptD*_Δ(Y658-Y678)::H | none | none | this study; spontaneous mutant resistant to LptD-targeting siphoviruses |
| AH-E01-044 | *E. coli* B REL606 | none | none | obtained from Dr. Jenna Gallie |
| AH-E03-168 | *E. coli* UTI89 | none | none | obtained from Prof. Urs Jenal |
| AH-E04-284 | *E. coli* CFT073 *rpoS(+)* | none | none | our laboratory collection [3] |
| AH-E06-481 | *E. coli* 55989 | none | none | our laboratory collection [3] |
| AH-E04-297 | *Salmonella enterica* subsp. *enterica* serovar Typhimurium 12023s (also known as ATCC 14028) | none | none | obtained from Prof. Dirk Bumann |
| AH-E06-438 | *S.* Typhimurium SL1344 | none | none | obtained from Prof. Mederic Diard |
| AH-E03-169 | *E. coli* K-12 EMG2 | none | none | most ancestral available *E. coli* K-12 strain; obtained from the Coli Genetic Stock Center (CGSC #4401) |
| AH-E01-053 | *E. coli* EB1484 (lysogen of phage P1 *clr100Km*) | P1 prophage | none | lysogen of a temperature-inducible P1 prophage tagged with kanamycin resistance; obtained from Prof. Kenneth Kreuzer |

The abbreviations in the selection column indicate the drug and its concentration that were used. Amp = ampicillin, Cam = chloramphenicol, Kan = kanamycin, Zeo = zeocin; 25 / 50 / 100 refer to 25 µg/ml, 50 µg/ml, and 100 = 100 µg/ml, respectively. The following mutants of the KEIO collection were used for qualitative top agar assays but are not included in the strain list because no phage showed any growth phenotype on them: *ompW::kanR*, *phoE::kanR*, *flgG::kanR*, *fepA::kanR*, *hofQ::kanR*, *cirA::kanR*, *fhuE::kanR*, *fiu::kanR*, *ompN::kanR*, *pgaA::kanR*, *chiP::kanR*, *ompL::kanR*, *yddB::kanR*, *fecA::kanR*, *uidC::kanR*, *nanC::kanR*, *yfaZ::kanR*, *bglH::kanR*, *bcsC::kanR*, *cusC::kanR*, *gfcE::kanR*, *mdtP::kanR*, *ompG::kanR*, *ompX::kanR*, *yfeN::kanR*, *csgF::kanR*, *wza::kanR*, *flu::kanR*, *nmpC::kanR*, *eaeH::kanR*, *ydiY::kanR*, *yiaT::kanR*, *yaiO::kanR*, *mdtQ::kanR*, *pgaB::kanR*, *mipA::kanR*, *pldA::kanR*, *yzcX::kanR*, *ydeT::kanR*, *blc::kanR*, *gspD::kanR*, *yjgL::kanR*

#### References (S1 Table)


1. Datsenko KA, Wanner BL. One-step inactivation of chromosomal genes in *Escherichia coli* K-12 using PCR products. Proc Natl Acad Sci USA. 2000;97(12):6640-5. doi: 10.1073/pnas.120163297. PubMed PMID: 10829079; PubMed Central PMCID: PMC18686.
2. Baba T, Ara T, Hasegawa M, Takai Y, Okumura Y, Baba M, et al. Construction of *Escherichia coli* K-12 in-frame, single-gene knockout mutants: the Keio collection. Mol Syst Biol. 2006;2:2006 0008. doi: 10.1038/msb4100050. PubMed PMID: 16738554; PubMed Central PMCID: PMC1681482.
3. Fino C, Vestergaard M, Ingmer H, Pierrel F, Gerdes K, Harms A. PasT of *Escherichia coli* sustains antibiotic tolerance and aerobic respiration as a bacterial homolog of mitochondrial Coq10. Microbiologyopen. 2020;9(8):e1064. Epub 2020/06/20. doi: 10.1002/mbo3.1064. PubMed PMID: 32558363; PubMed Central PMCID: PMCPMC7424257.

#### S2 Table. List of all oligonucleotide primers used in this study

| *Primer name* | *Sequence (5'-3')* |
|---|---|
| prAH1815 | GCCGCTCCCGATGTGGTGTCGGGAGCGGTATTTTCTATAAAACTTACCGCGGAACTTCATTTAAATGGCG |
| prAH1816 | AGTAAGGGGTTATGGGCCGGATAAGGCGCAGCCGCATCCGGCCTGATATTGGTCCATATGAATATCCTCCTTAG |
| prAH1817 | ATGTGGTGTCGGGAGCGGTATTTTCTATAAAACTTACCGCAATATCAGGCCGGATGCGGCTGCGCCTTATCCGGCCCATA |
| prAH1818 | TATGGGCCGGATAAGGCGCAGCCGCATCCGGCCTGATATTGCGGTAAGTTTTATAGAAAATACCGCTCCCGACACCACAT |
| prAH1823 | TCAATAAAAGTAGTATTGTCGTGAAAAATTGATTAAAGATTAATATTATGGGTCCATATGAATATCCTCCTTAG |
| prAH1825 | ACGCCCGTTTCAATATTTAACACATGGAGAGATTACATGTTTTCGATGATGGAACTTCATTTAAATGGCG |
| prAH1826 | TAGTATTGTCGTGAAAAATTGATTAAAGATTAATATTATGATCATCGAAAACATGTAATCTCTCCATGTGTTAAATATTG |
| prAH1827 | CAATATTTAACACATGGAGAGATTACATGTTTTCGATGATCATAATATTAATCTTTAATCAATTTTTCACGACAATACTA |
| prAH1944 | GCTGGCGGAGGCCCTGACCGGGCTGCGCTGCCTGTACGGAGATCTGTAATTACAACTTTTTTTACTTCTTGTTCATTAG |
| prAH1945 | ATCTCCTTCATTTAGTAAGAAAAAGGCCGCTAAGCGGCCTTAATTTTTGGCTTTTGCTCACATGTTGGTC |
| prAH1948 | GGGAACGCGGCCGCACCTACATCTGTATTAACGAAGCG |
| prAH1949 | CTGCTTCTCGAGACACGGTGCCTGACTGC |
| prAH1950 | GGGTGACTCGAGATGAAGAATGGTTTTTATGCG |
| prAH1951 | GCATTTGCGGCCGCTTATTCTTGTTCTCTGGTCAAATTATAT |
| prAH1956 | GGGAGAATCGATAGAATCAGGTAGATGTTTTTCGG |
| prAH1957 | GCATTTGCGGCCGCTCAGGATTTTTTACGTGAGGC |
| prAH1981 | CACCTACATCTGTATTAACGAAGC |
| prAH1982 | GATAAGCTGTCAAACATGAGAATTC |
| prAH2009 | AATCAACAACCGTATCAGAATAGATACTTTCTTTAGGAATTTTTGTTTTACGTAATTTTTTTAAGGCAGTTATTG |
| prAH2010 | AAATTTCCTGTGCTTTCTGATTTTATTGTGTCATTTATGTTAGGGATTAACTTACTGTCCCTAGTGCTTGG |
| prAH2013 | GTGAAAAACTGATGAAATTCGAT |
| prAH2014 | GACATGAAGACTACATCAAAAAATTACT |
| prAH2015 | GTTATCACCAGAGCTTAATCGAC |
| prAH2016 | TATCTCTAAAATCATTGATGATTTCAG |
| prAH2019 | GTTTTTTGACCTCTGCAAAAG |
| prAH2020 | TGACCGCAACAAAAAAATATC |
| prAH2184 | GAATTCTCATGTTTGACAGCTTATCACAGCTTAAAAACGAACTTGAAG |
| prAH2185 | CGCTTCGTTAATACAGATGTAGGTGCTTACTTCACCACTTCCATCAG |
| prAH2224 | CCCTTTGATATGTAACGGTGAAC |
| prAH2225 | GTTAATGTCATGATAATAATGGTTTCTTAGAC |
| prAH2266 | GGTATTTTCTCCTTACGCATCTG |
| prAH2343 | GGTTTCTTAGACGTCAGGTGG |
| prAH2344 | AGTGCCACCTGACGTCTAAGAAACCCATTACAAGAGTTTGCTGACAAG |
| prAH2345 | CACAGATGCGTAAGGAGAAAATACCCTATTTTTAACGACCTGAGCG |
| prAH2346 | AGTGCCACCTGACGTCTAAGAAACCTGTTAGAGTTGATACGGTTCCTG |
| prAH2347 | CACAGATGCGTAAGGAGAAAATACCTGGTGATGTGAATAAAGCGG |
| prAS046 | GAATTCTCATGTTTGACAGCTTATCTTAAATCCATTTTATGAAATCTTCC |
| prAS047 | CGCTTCGTTAATACAGATGTAGGTGACTAATGAGCCATCAGTATTTCC |
| prAS048 | GAATTCTCATGTTTGACAGCTTATCTCAATTGAGTATCGATTTTCGT |
| prAS049 | CGCTTCGTTAATACAGATGTAGGTGACAGCACAGTACTAAACCAATAGTG |
| prAS050 | GAATTCTCATGTTTGACAGCTTATCATTACTATGAGGTGAATGGCAAG |
| prAS051 | CGCTTCGTTAATACAGATGTAGGTGTCTGACAGTTTCCTTTGAGC |
| prMBu0025 | TAATATTGATGAAACCTGCGGCATCCTTCTTCTATTGTGGATGCTTTACAACCTGCAGTTCGAAGTTCC |
| prMBu0026 | CGTGTCGGTAATCGCATTGCGCGCATCGACATAATCATAACTCACAGTATGAGCTGCTTCGAAGTTCCTA |

| | |
|---|---|
| prMBu0075 | TACAGTTTGATCGCGCTAAATACTGCTTCACCACAAGGAATGCAAACCTGCAGTTCGAAGTTCC |
| prMBu0076 | TACGTTGCCTTACGTTCAGACGGGGCCGAAGCCCCGTCGTCGTCATGTAGGCTGGAGCTGCTTC |
| prMBu0078 | AAACCATTATTATCATGACATTAACTTCAAGAATACGGCTGGTC |
| prMBu0079 | CTGTTCACCGTTACATATCAAAGGGTCAATCATCTTATCGACTACCTTG |

### S3 Table. List of all plasmids used in this study

| name | Selection | Description | Source |
|---|---|---|---|
| pWRG99 | Amp100 | lambda red recombineering plasmid with inducible I-SceI | our laboratory collection [1] |
| pWRG100 | Cam25 | template plasmid for the recombineering double-selectable cassette encoding chloramphenicol resistance and an I-SceI recognition site | our laboratory collection [1] |
| pJM05 | Kan25 | template plasmid for the recombineering double-selectable cassette encoding kanamycin resistance and *sacB* | our laboratory collection [2] |
| pUA139 | Kan25 | SC101 origin of replication, kanamycin resistance, and *gfpmut2* (originally to clone promoter-GFP fusions) | our laboratory collection [3] |
| pUA139_T7(*gp17*) | Kan25 | encoding part of the *gp17* tail fiber gene of phage T7 with flanking sequence | this study |
| pUA139_cat-sacB_v3 | Cam25, Kan25 | template plasmid for recombineering double-selectable cassettes encoding *sacB* and either chloramphenicol or kanamycin resistance | our laboratory collection |
| pPICZa | Zeo50 | *Pichia pastoris* expression vector with a zeocin resistance cassette | obtained from Prof. Urs Jenal |
| pAR280 | Amp50 | mini-R1 plasmid encoding the intact *E. coli* K-12 *wbbL* open reading frame | obtained from Prof. Urs Jenal |
| pBR322_ΔP*tet* | Amp50 | variant of pBR322 in which the tetracycline resistance cassette and its promoter have been deleted; empty-vector control for immunity experiments | obtained from Prof. Călin Guet via Dr. David Thaler [4] |
| pAH186_SC101e | Amp50 | plasmid encoding ampicillin resistance and an SC101 low-copy origin of replication | our laboratory collection [5] |
| pAH213_EcoKI | Amp50 | pBR322 derivative expressing type I RM system EcoKI of *E. coli* K-12 | this study |
| pAH213_EcoCFT_I | Amp50 | pAH186_SC101e derivative expressing type I RM system EcoCFT_I of *E. coli* CFT073 | this study |
| pEcoRI | Amp50 | pBR322 derivative expressing type II RM system EcoRI | obtained from Prof. Călin Guet via Dr. David Thaler [4] |
| pEcoRV | Amp50 | pBR322 derivative expressing type II RM system EcoRII | obtained from Prof. Călin Guet via Dr. David Thaler [4] |
| pAH213_EcoCFT_II | Amp50 | pBR322 derivative expressing type III RM system EcoCFT_II of *E. coli* CFT073 | this study |
| pAH213_EcoP1_I | Amp50 | pAH186_SC101e derivative expressing type I RM system EcoP1_I of *E. coli* phage P1 | this study |
| pAH213_RexAB | Amp50 | pBR322 derivative expressing the RexAB Abi system of *E. coli* phage lambda | this study |
| pAH200e | Kan25 | F-plasmid in which the *tn1000* locus was replaced with a kanamycin resistance cassette by recombineering | obtained from Prof. Christoph Dehio |
| pAH213_Fun/Z | Amp50 | pBR322 derivative expressing the Fun/Z Abi system of *E. coli* phage P2 | this study |
| pAH213_Old | Amp50 | pBR322 derivative expressing the Old Abi system of *E. coli* phage P2 | this study |
| pAH213_Tin | Amp50 | pBR322 derivative expressing the Tin Abi system of *E. coli* phage P2 | this study |

The abbreviations in the selection column indicate the drug and its concentration that were used. Amp = ampicillin, Cam = chloramphenicol, Kan = kanamycin, Zeo = zeocin; 25 / 50 / 100 refer to 25 µg/ml, 50 µg/ml, and 100 = 100 µg/ml, respectively.

### References (S3 Table)
