## Supplementary material for "Systematic exploration of *Escherichia coli* phage-host interactions with the BASEL phage collection": S1 Text

### S1 Text. Construction of bacterial mutant strains

Mutants strains of *Escherichia coli* were generally constructed using recombineering [1]. For this purpose, strain *E. coli* K-12 MG1655 (laboratory wildtype; CGSC #6300; see S1 Table) and derivatives or variants were transformed with one of several plasmids carrying the lambda red recombineering functions. In some cases, we used plasmid pWRG99 that encodes the lambda red recombineering functions as well as I-SceI for negative selection of the double-selectable chloramphenicol resistance cassette (of template plasmid pWRG100) carrying an I-SceI site (called *camR-I-SceI* in the following) [2]. Alternatively, plasmid pKM208 was used [3]. A clean deletion or allelic exchange was achieved using a two-step procedure in which first a target locus was replaced by recombineering with a double-selectable cassette carrying an antibiotic resistance gene for positive selection (chloramphenicol or kanamycin resistance denoted as *camR* or *kanR*). Subsequently, the double-selectable cassette was replaced with the desired alternative allele or deletion allele using recombineering and negative selection marker against an I-SceI site or the *sacB* gene (conferring sucrose sensitivity) on the double-selectable cassette. As an alternative to pWRG100 encoding a *camR-I-SceI* double-selectable cassette, we sometimes used another double-selectable cassette (based on plasmid pJM05) encoding a kanamycin resistance cassette and *sacB* for negative selection (in the following called *kanR-sacB*) that we had already used earlier [4]. Alternatively, a variant of plasmid pUA139 [5] carrying the *camR-sacB* double-selectable cassette of pKO4 [6] – known as pUA139_cat-sacB_v3 – was used. This plasmid is a convenient PCR template for *sacB* either in combination with a kanamycin or chloramphenicol marker. For increased efficiency of recombineering in *E. coli* K-12 strains expressing functional EcoKI, we had initially edited out the EcoKI site in the chloramphenicol resistance cassette of pKO4.

#### 1) Construction of *Escherichia coli* K-12 MG1655 ΔRM

All known strains of *E. coli* K-12 have long lost costly production of O-antigen glycan chains, the outermost part of the lipopolysaccharide (LPS) on the cell surface [7], which in other *E. coli* would effectively shield the receptors of many phages on the bacterial cell surface [8-10]. However, K-12 strains encode the well-known EcoKI type I RM system and three type IV restriction systems as well as the RexAB and PifA abortive infection systems (carried by the lambda prophage and the F-plasmid, respectively) [11, 12]. In order to avoid an interference of these systems with phage isolation, we used the common MG1655 laboratory strain of *E. coli* K-12 (which had been cured of lambda prophage as well as F-plasmid long ago [13]), and additionally deleted the four restriction systems, creating strain *E. coli* K-12 MG1655 ΔRM. Subsequently, we re-introduced the F-plasmid because its sex pilus can be used as a phage receptor, but used a variant on which we had deleted *pifA* (see *Materials and Methods* and below).

As a first step of in the construction of *E. coli* K-12 MG1655 ΔRM the parental strain *E. coli* K-12 MG1655 was transformed with plasmid pWRG99. In order to delete type I restriction-modification (RM) system EcoKI and type IV restriction systems Mrr and mcrBC that are encoded close to each other, we amplified a *camR-I-SceI* with suitable 50 bp homologies using oligonucleotide primers prAH1815 / prAH1816 from template plasmid pWRG100 and recombineered this cassette into the chromosome of *E. coli* K-12 MG1655 to generate strain *E. coli* K-12 MG1655 *mrr-hsdRMS-mcrBC::DScas(I-SceI)*. For a clean deletion, we annealed complementary 80 nt oligonucleotides spanning the desired deletion site with 40 nt on each side (prAH1817 / prAH1818) and subsequently recombineered the resulting double-stranded DNA molecule into *E. coli* K-12 MG1655 *mrr-hsdRMS-mcrBC::DScas(I-SceI)*. *E. coli* K-12 MG1655 Δ*mrr-hsdRMS-mcrBC* carrying pWRG99 was stocked as AH-E02-160. Note that the deletion also removes the *symER* type I toxin-antitoxin module that has no known biological function [14].

In order to delete the *mcrA* type IV restriction system in *E. coli* K-12 MG1655 Δ*mrr-hsdRMS-mcrBC* (AH-E02-160), we amplified a *kanR-sacB* doubleselectable cassette with suitable 50 bp homologies using oligonucleotide primers prAH1823 / prAH1825 from template plasmid pJM05 and recombineered this cassette into the chromosome of this strain to generate *E. coli* K-12 M1655 Δ*mrr-hsdRMS-mcrBC mcrA::DScas*. A clean deletion of *mcrA* was achieved as described above using recombineering with annealed 80 nt oligonucleotides prAH1826 / prAH1827 that span the desired deletion site, followed by the elimination of temperature-sensitive plasmid pWRG99 by growth at 43°C. The resulting strain *E. coli* K-12 M1655 Δ*mrr-hsdRMS-mcrBC* Δ*mcrA* (called ΔRM throughout our work) was stocked as AH-E03-200.

#### 2) Construction of plasmid F(*pifA::zeoR*)

In order to knock out *pifA* on the F-plasmid of *E. coli* K-12 and at the same time introduce a selection marker onto this plasmid, the *pifA* gene was replaced with a zeocin resistance cassette by recombineering. For this purpose, *E. coli* K-12 W1872 (CGSC #6538) that carries a wildtype F-plasmid was transformed with recombineering plasmid pKM208 [3] and the zeocin resistance cassette (*zeoR* in short) was amplified from template plasmid pPICZa with suitable 50 bp homologies using oligonucleotide primers prAH1944 / prAH1945. The resulting strain *E. coli* K-12 W1872 F(*pifA::zeoR*) was mated with *E. coli* K-12 MG1655 ΔRM carrying pBR322_ ΔP*tet* by simply mixing small amounts of overnight cultures of both strains (washed once in LB medium to remove antibiotic supplements) in an Eppendorf tube and incubating the mixture at 37°C for three hours. Plating on LB agar plates containing 100 μg/ml ampicillin and 50 μg/ml zeocin enabled selection for *E. coli* K-12 MG1655 ΔRM carrying pBR322_ ΔP*tet* and F(*pifA::zeoR*) which was stocked as AH-E03-217. Successful transfer of F(*pifA::zeoR*) was verified by confirming sensitivity of AH-E03-217 to phages targeting the F sex pilus while retaining sensitivity to phage T7 (like *E. coli* K-12 W1872 F(*pifA::zeoR*), wile the ancestral *E. coli* W1872 was resistant to phage T7).

##### 3) Construction of *Escherichia coli* K-12 MG1655 ΔRM mutants with altered surface glycans

*waaC* and *waaG* were knocked out in the *E. coli* K-12 MG1655 ΔRM strain background using the pKD13-derived kanamycin resistance cassette (*kanR*) constructs of the KEIO collection [15]. The *waaC::kanR* and *waaG::kanR* constructs were amplified from the respective KEIO collection mutant strains using primer pairs prAH2015 / prAH2016 and prAH2019 / prAH2020, respectively. Subsequently, the cassettes were recombineered into *E. coli* K-12 MG1655 ΔRM carrying pWRG99. After eliminating temperature-sensitive plasmid pWRG99 by growth at 43°C, the resulting strains *E. coli* K-12 MG1655 ΔRM *waaC::kanR* and *E. coli* ΔRM *waaG::kanR* were stocked as AH-E03-233 and AH-E03-235, respectively.

In order to remove the IS5 element disrupting the *wbbL* gene in *E. coli* K-12 MG1655 ΔRM carrying pWRG99, we first used recombineering to precisely replace it with the *kanR-sacB* cassette of pUA139_cat-sacB_v3 that had been amplified with prAH2009 / prAH2010. Subsequently, we amplified part of the intact open reading frame of *wbbL* from plasmid pAR280 using prAH2013 / prAH2014 and used the PCR product to cut out the *kanR-sacB* cassette previously inserted at *wbbL*. After eliminating temperature-sensitive plasmid pWRG99 by growth at 43°C, the resulting strain *E. coli* K-12 MG1655 ΔRM *wbbL(+)* was stocked as AH-E04-243.

##### 4) Construction of *Escherichia coli* K-12 BW25113 *btuB::kanR* and *tolC::kanR* mutants

The KEIO collection lacks a true *btuB* mutant strain [16], probably because the attempted deletion of *btuB* by recombineering apparently used homologies that affected functional expression of the essential *murI* gene [17, 18] which is encoded downstream of *btuB* with an overlap of almost 60 bp. We therefore selected alternative 50 bp homologies that would 1) remove the complete 5' end of *btuB* including the start codon and 2) leave the last ca. 300 bp at the 3' end of *btuB* intact in order to not abolish *murI* expression. The *kanR* cassette of classical recombineering template plasmid pKD13 was amplified with these homologies using oligonucleotide primers prMBu0025 / prMBu0026 [19]. Subsequently, the PCR product was recombineered into *E. coli* K-12 BW25113 carrying pKM208. After curing temperature-sensitive plasmid pKM208 by growth at 43°C, the resulting strain *E. coli* K-12 BW25113 *btuB::kanR* was stocked as AH-E04-321.

The *E. coli* K-12 BW25113 *tolC::kanR* mutant included in our copy of the KEIO collection was unavailable due to apparent contamination of the frozen stock. We therefore amplified the generated a new *tolC::kanR* deletion mutant in which the full *tolC* open reading frame was replaced by the *kanR* cassette of pKD13. For this purpose, we amplified *kanR* with suitable 50 bp homologies using prMBu0075 and prMBu0076. Subsequently, the PCR product was recombineered into *E. coli* K-12 BW25113 carrying pKM208. After curing temperature-sensitive plasmid pKM208 by growth at 43°C, the resulting strain *E. coli* K-12 BW25113 *tolC::kanR* was stocked as MBu-E01-044.

### References (S1 Text)
