## Supplementary material for "Systematic exploration of *Escherichia coli* phage-host interactions with the BASEL phage collection": S2 Text

### S2 Text. Generation of the Maximum-Likelihood phylogenies shown in this article.

For the *Drexlerviridae* phylogeny shown in Fig 4B, the orthologs of a DNA helicase gene (*T1p47* in T1), the major capsid protein gene (*T1p47* in T1), the tail tape measure protein gene (*T1p38* in T1), the large terminase subunit gene (*T1p53* in T1), and another DNA helicase (*T1p09* in T1) were used. We aligned each set of orthologs, curated the alignments, concatenated them, and calculated a Maximum-Likelihood phylogeny as described above.

The phylogeny of *Drexlerviridae* and *Dhillonvirus* phages shown in S1A Fig is based on a concatenated alignment of amino acid sequences of major capsid protein (T1p47 in T1) and large terminase subunit (T1p53 in T1) of all phages that were included.

For the phylogenies of the three *Siphoviridae* genera shown in Fig 6B and Fig. 6D, each one whole-genome alignment was manually curated and then used to calculate a Maximum-Likelihood phylogeny. The lengths of these alignments were 21.8 kb (*Dhillonvirus*), 30.7 kb (*Nonagvirus*), and 33 kb (*Seuratvirus*). We present the phylogenies of *Nonagvirus* and *Seuratvirus* in Fig 6D on opposite sides of a single phylogeny because the two genera are well-known to be sister clades [1].

The phylogeny of *Markadamsvirinae* shown in Fig 7B was calculated based on a concatenation of curated nucleotide sequence alignments of the DNA polymerase gene (*T5.122* in T5), a DNA helicase gene (*T5.124* in T5), the major capsid protein gene (*T5.149* in T5), the DNA primase gene (*T5.108* in T5), another DNA helicase gene (*T5.119* in T5), the tail tape measure protein gene (*T5.140* in T5), and the large terminase subunit gene (*T5.155* in T5).

For the phylogeny of *Tevenvirinae* genera *Tequatrovirus* and *Mosigvirus* in Fig 8B, we used each one whole-genome alignment that was manually curated and then used to calculate Maximum-Likelihood phylogeny. The lengths of these alignments were 29.1 kb (*Tequatrovirus*) and 57.9 kb (*Mosigvirus*). The phylogeny of short tail fibers in Fig 8D was generated based on an amino acid sequence alignment of the orthologs in all included *Tevenvirinae* genomes (T4p157/Gp12 in T4).

The phylogeny of *Vequintavirinae* and relatives in Fig 9B was assembled from a phylogeny of *Vequintavirinae sensu stricto* (top) the related clusters of phages including phAPEC8 and phi92 (bottom). We present these two phylogenies with a common root since these groups of phages are known to be closely related [1-3]. For the *Vequintavirinae sensu stricto*, the phylogeny was calculated based on a concatenation of curated nucleotide sequence alignments of three conserved loci of the phage core, the DNA packaging region (from the i-spanin gene (*rv5_gp068* in rV5) until the first tail genes (*rv5_gp054* in rV5), a locus comprising *rIIAB* and DNA replication functions (from *rIIA* (*rv5_001* in rV5) to the DNA polymerase (*rv5_gp223* in rV5), and a locus around the NTP reductase genes (from a *phoH*-like gene (*rv5_gp115* in rV5) to the thymidylate synthase (*rv5_gp102* in rV5). We curated the alignments, concatenated them, and calculated a Maximum-Likelihood phylogeny as described above. The phylogeny of phAPEC8-like and phi92-like phages was calculated based on a whole-genome alignment that was manually curated, resulting in a final length of 22.2 kb.

For the phylogeny of *Autographiviridae* phages in Fig 10B, we extracted the orthologs of the DNA polymerase gene (*T7p29* in T7), the DNA primase / helicase gene (*T7p22* in T7), the T3/T7 family RNA polymerase gene (*T7p07* in T7), and the large terminase subunit gene (*T7p57* in T7). We aligned each set of orthologs, curated the alignments, concatenated them, and calculated a Maximum-Likelihood phylogeny.

The phylogeny of *Enquatrovirus* phages and related *Podoviridae* in Fig 10E was calculated based on a concatenation of curated nucleotide sequence alignments of the DNA primase gene (*EPNV4_gp43* in N4), the terminase large subunit gene (*EPNV4_gp68* in N4), and the virion RNA polymerase gene (*EPNV4_gp50* in N4).

For the phylogeny of *Felixounavirus* phages and related *Ounavirinae* in Fig 11B, we extracted the genes coding for a DNA ligase (*Felix01p163* in Felix O1), a DNA primase / helicase (*Felix01p188* in Felix O1), the major capsid protein (*Felix01p112* in Felix O1), and the tail tape measure protein (*Felix01p122* in Felix O1). Each set of orthologs was aligned and the alignments curated, concatenated, and used to calculate a Maximum-Likelihood phylogeny.

### References (S2 Text)


1. Korf IHE, Meier-Kolthoff JP, Adriaenssens EM, Kropinski AM, Nimtz M, Rohde M, et al. Still Something to Discover: Novel Insights into *Escherichia coli* Phage Diversity and Taxonomy. Viruses. 2019;11(5). doi: 10.3390/v11050454. PubMed PMID: 31109012; PubMed Central PMCID: PMCPMC6563267.
2. Kropinski AM, Waddell T, Meng J, Franklin K, Ackermann HW, Ahmed R, et al. The host-range, genomics and proteomics of *Escherichia coli* O157:H7 bacteriophage rV5. Virol J. 2013;10:76. Epub 2013/03/19. doi: 10.1186/1743-422X-10-76. PubMed PMID: 23497209; PubMed Central PMCID: PMCPMC3606486.
3. Schwarzer D, Buettner FF, Browning C, Nazarov S, Rabsch W, Bethe A, et al. A multivalent adsorption apparatus explains the broad host range of phage phi92: a comprehensive genomic and structural analysis. Journal of virology. 2012;86(19):10384-98. Epub 2012/07/13. doi: 10.1128/JVI.00801-12. PubMed PMID: 22787233; PubMed Central PMCID: PMCPMC3457257.
