## Supplementary material for "Systematic exploration of *Escherichia coli* phage-host interactions with the BASEL phage collection": S3 Text

### S3 Text. Technical considerations regarding the composition of the BASEL collection and the phenotyping of bacterial defense systems

#### Considerations regarding the composition of the BASEL collection

The choice of isolation host is known to be "perhaps the most critical part of the isolation process" because it pre-determines the properties and range of phages that can be sampled [1]. We therefore specifically designed a highly phage-sensitive host strain, *Escherichia coli* K-12 ΔRM, in order to eliminate as many biases in the isolation process as possible (see *Materials and Methods* as well as *Composition of the BASEL collection* in the main text).

However, one important feature of *E. coli* K-12 ΔRM is that it displays the rough LPS of all K12-lineage laboratory strains and lacks the O-antigen chains that are otherwise fully covering the cell surface of natural enterobacterial isolates (Figs 3A and 12A) [2]. On one hand, the absence of this formidable barrier enabled the isolation of a huge diversity of phages from all families that are, with very few exceptions, largely unable to infect *E. coli* K-12 with restored O16-type O-antigen expression (Figs 3A, 4, and 6-11). However, using a strain with rough LPS as isolation host intrinsically excludes any phages for which the O-antigen is not only an optional primary receptor but an essential part of the host recognition. Two common examples for such phages are iconic *Salmonella* phage P22 and *Gamaleyavirus* G7C, a relative of N4 (see Fig 10E), that both bind to very specific types of O-antigen and then target the glycan chain by enzymatic activities of their tailspikes to generate directional movement towards the cell surface [3, 4]. It is clear that this kind of phages are not highly abundant among *E. coli* phages because, e.g., a study that had isolated fifty phages primarily using strains with smooth LPS found 1) mostly the same taxonomic groups as those in the BASEL collection and reported that 2) many phages isolated on smooth hosts can also infect *E. coli* K-12 [5]. However, the targeting of O-antigen chains as an essential primary receptor is not uncommon among podoviruses (and some myoviruses) [6, 7], and the isolation of *E. coli* phages with hosts expressing full smooth LPS indeed revealed a bigger diversity of (narrow-host-range) podoviruses than was reported in studies solely relying on *E. coli* K-12 [5]. Given seemingly low abundance of O-antigen specialists and that most realistic applications of the BASEL collection will be based on infecting *E. coli* K-12 strains with rough LPS, we do not feel that the absence of these O-antigen-dependent phages compromises the usefulness of our work. However, we imagine that a future study could generate a dedicated, optional expansion of the BASEL collection containing a diverse and representative selection of these phages that are specialized in the O16-type O-antigen of *E. coli* K-12. Similarly, we excluded all tailless phages like *Microviridae* and *Inoviridae* from the BASEL collection because their biology, evolution, and host interactions are so different from the lytic *Caudovirales* that we feel their investigation is beyond the scope of our current work [8, 9].

Any bacteriophage collection of finite size is inherently unable to include all rare phage groups and cannot comprehensively cover all genera of highly diverse and abundant families such as the *Drexlerviridae* (Fig 4B). Despite this shortcoming, we feel that the BASEL collection provides a reasonably complete snapshot of *E. coli* phage diversity because, e.g., all common and almost all previously described protein receptors of all included phage groups are covered (Figs 4-8). Increasing the number of phage isolates in the BASEL collection might therefore not greatly increase its biological diversity but could jeopardize its usefulness by making the handling more complicated. It also does not appear that sampling most phages from sewage plant inflow is a major limitation, because we did not observe any difference in the phages sampled from sewage and, e.g., river water (S5 Table). Previous work had sampled overall similar sets of phage groups on matter if the phages came from sewage or from infant guts [5, 10-15], possibly because many *E. coli* phages in the environment might derive directly or indirectly from fecal contaminations.

Another limitation of our isolation host strain *E. coli* K-12 ΔRM is that it might introduce additional biases beyond its rough LPS phenotype. As an example, laboratory adaptation might have inactivated also other possible phage receptors on the cell surface. A relevant example for the possible consequences of such a bias is the observation that phage T5 of the T phages is a highly unusual member of the *Markadamsvirinae* subfamily of *Demerecviridae* because it uses the FhuA protein as its terminal receptor (Fig 7) [16]. This peculiar feature of the T phages is easily explained with the *btuB* loss-of-function mutation of many early *E. coli* B strains that were used to sample the T phages but later reverted back to functional BtuB expression, e.g., in the lineage leading to *E. coli* B REL606 that is sensitive to all tested *Markadamsvirinae* (Fig 12B) [16, 17]. Similarly, the T phages do not contain any *Vequintavirinae* phage despite the abundance of this group, and we indeed find that none of our twelve *Vequintavirinae sensu stricto* can lyse *E. coli* B REL606 (Fig 12B). Since this defect is probably due to impaired adsorption, we speculate that the truncated *E. coli* B LPS core and / or changes of the ECA or any other primary receptor might be responsible for this surprising phenotype.

Another possible factor biasing the diversity of isolated bacteriophages could be remaining immunity systems in the sampling strain *E. coli* K-12 ΔRM. While we strain lacks all known restriction systems of *E. coli* K-12 as well as the RexAB and PifA Abi systems (S1 Text), several systems remain that either have been poorly studied or are supposed to have only a narrow target range [18]. As an example, the cryptic prophage *e14* of *E. coli* K-12 encodes the Lit Abi system that cleaves the elongation factor Tu (EF-Tu) when sensing the major capsid protein of *Tevenvirinae* [18, 19]. It is intuitive that the presence of *lit* in *E. coli* K-12 ΔRM might somehow bias the range of *Tevenvirinae* phages that we are sampling. However, although we confirmed by whole-genome sequencing that both the host as well as the T4 phage should carry alleles of *lit* and the major capsid protein gene triggering abortive infection [19], we observe robust plaque formation of T4 on our *E. coli* K-12 strains (see, e.g., in S4B Fig). Similarly, recent work growing phage T4 on *E. coli* K-12 hosts never reported any problems with abortive infection [5, 20, 21]. We are therefore skeptical if abortive infection by *lit* can significantly affect the isolation of bacteriophages with *E. coli* K-12 ΔRM. Besides Lit, the RnlAB type II toxin-antitoxin system aborts the growth of T4 by activation of the RnlA RNase toxin when the phage lacks the *dmd* antitoxin gene [18, 22]. However, given the broad conservation of *dmd* among *Tevenvirinae* and the absence of any reports that RnlAB could target different phages, we do not think that this system has a significant impact on our bacteriophage isolation experiments. Like for Lit and RnlAB, we see no evidence that the other proposed immunity systems of *E. coli* K-12 will have relevant impact on the diversity of sampled phages. These are mostly ill-characterized or, in case of DicB, were only shown to work when ectopically expressed [23].

#### Considerations regarding the phenotyping of bacterial immunity systems

Our phage phenotyping experiments with diverse immunity systems were performed as quantitative top agar assays to determine the efficiency of plating (EOP) of each phage on a host carrying a given immunity system to generate robust, quantitative data (see *Materials and Methods*). However, there are a few technical caveats associated with our approach that need to be considered for a comprehensive interpretation of our results. Most importantly, for the phenotyping in the *E. coli* K-12 ΔRM host we cloned the different immunity systems onto plasmids which, though most were cloned with their native promoter, might affect their functionality by change of copy number (see *Materials and Methods* as well as S3 and S4 Tables). As an example, it was shown that phage T4 wildtype is resistant to RexAB but becomes sensitive when this immunity system is overexpressed from a multicopy plasmid [24]. However, the ColE1 and SC101 origins of replications used in this study have rather low copy numbers (of around 40 and 3-4, respectively [25]) and, e.g., our *rexAB* construct with ColE1 origin of replication has no detectable effect on the growth of phage T4 or any relative (Fig 8C), while an *rIIAB* mutant of this phage was sensitive as described previously (S4B and S4C Figs) [18]. The use of two different origins of replication is a consequence of our initial choice to clone immunity systems onto plasmids isogenic to the EcoRI and EcoRV plasmids of Pleška, Qian et al. [26] which later resulted in problems with toxicity for some constructs. These were then cloned into the lower-copy plasmid with SC101 origin of replication instead (S4 Table). Though it is clear that expression levels could be different between these backbones as already evidenced form the difference in toxicity, we do no feel that this compromises our approach considerably. As an example, the two type III RM systems EcoP1_I and EcoCFT_II have a similar recognition sequence (Fig 3B) that is similarly abundant in the diverse phage genomes (S5 Table), but EcoP_1 was cloned with an SC101 origin of replication while EcoCFT_II was cloned with a ColE1 origin of replication. However, the phenotyping data do not show EcoCFT_II would be more potent compared to EcoP1_I (Figs 4 and 6-11) – sometimes one or the other has the stronger effect on EOP, sometimes they are similar.

Another more transparent caveat is that the graphs displaying the EOP of different phages on hosts with restriction-modification (RM) systems are not directly comparable quantitatively. The reason is that, due to differences in genome size, GC content, and evolution towards restriction site avoidance, the different genomes have vastly different numbers of recognition sites for a given RM system [27] (S5 Table). In extreme cases such as for type II RM systems and many tested podoviruses, recognition sites can be completely absent (S5 Table). One might argue that displaying an EOP (of ca. 1) for phage / RM interactions in the absence of recognition sites is not useful because – just biochemically – a cleavage of the phage chromosome was barely possible. However, given that the number of these sites is strongly under selection and part of the phages' strategy to counter their host's defenses, we find it more appropriate to show all data and highlight for the reader when no recognition sites were present.

#### References (S3 Text)

204
