## Supplemental Figures for "Systematic exploration of *Escherichia coli* phage-host interactions with the BASEL phage collection"

Figure S1

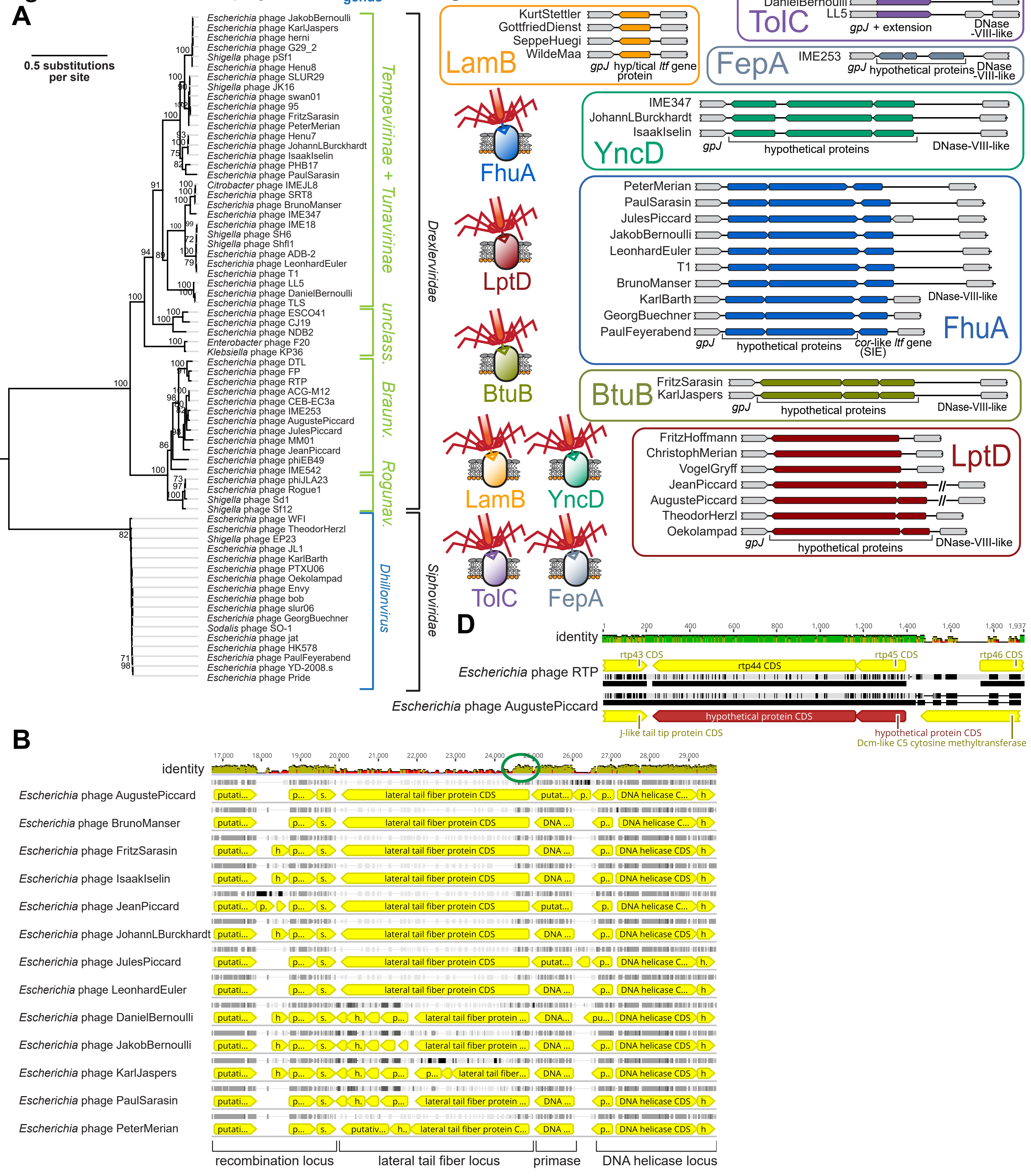
A
B
C
D
phage name
subfamily
genus
family
0.5 substitutions per site
Escherichia phage JakobBernoulli
Escherichia phage KarlJaspers
Escherichia phage herni
Escherichia phage G29_2
Shigella phage pSf1
Escherichia phage Henu8
Escherichia phage SLUR29
Shigella phage JK16
Escherichia phage swan01
Escherichia phage 95
Escherichia phage FritzSarasin
Escherichia phage PeterMerian
Escherichia phage Henu7
Escherichia phage JohannLBurckhardt
Escherichia phage IsaakIselin
Escherichia phage PHB17
Escherichia phage PaulSarasin
Citrobacter phage IMEJL8
Escherichia phage SRT8
Escherichia phage BrunoManser
Escherichia phage IME347
Escherichia phage IME18
Shigella phage SH6
Shigella phage Shfl1
Escherichia phage ADB-2
Escherichia phage LeonhardEuler
Escherichia phage T1
Escherichia phage LL5
Escherichia phage DanielBernoulli
Escherichia phage TLS
Escherichia phage ESCO41
Escherichia phage CJ19
Escherichia phage NDB2
Enterobacter phage F20
Klebsiella phage KP36
Escherichia phage DTL
Escherichia phage FP
Escherichia phage RTP
Escherichia phage ACG-M12
Escherichia phage CEB-EC3a
Escherichia phage IME253
Escherichia phage AugustePiccard
Escherichia phage JulesPiccard
Escherichia phage MM01
Escherichia phage JeanPiccard
Escherichia phage phiEB49
Escherichia phage IME542
Escherichia phage phiJLA23
Escherichia phage Rogue1
Shigella phage Sd1
Shigella phage Sf12
Escherichia phage WFI
Escherichia phage TheodorHerzl
Shigella phage EP23
Escherichia phage JL1
Escherichia phage KarlBarth
Escherichia phage PTXU06
Escherichia phage Oekolampad
Escherichia phage Envy
Escherichia phage bob
Escherichia phage slur06
Escherichia phage GeorgBuechner
Sodalis phage SO-1
Escherichia phage jat
Escherichia phage HK578
Escherichia phage PaulFeyerabend
Escherichia phage YD-2008.s
Escherichia phage Pride
Tempevirinae + Tunavirinae
unclass.
Braunv.
Rogunav.
Dhillonvirus
Drexlerviridae
Siphoviridae
1 200 400 600 800 1000 nucleotides
LamB
KurtStettler
GottfriedDienst
SeppeHuegi
WildeMaa
gpJ hyp/tical ltf gene protein
TolC
TLS
DanielBernoulli
LL5
gpJ + extension
DNase -VIII-like
FepA
IME253
gpJ hypothetical proteins DNase -VIII-like
YncD
IME347
JohannLBurckhardt
IsaakIselin
gpJ hypothetical proteins DNase-VIII-like
FhuA
PeterMerian
PaulSarasin
JulesPiccard
JakobBernoulli
LeonhardEuler
T1
BrunoManser
KarlBarth
GeorgBuechner
PaulFeyerabend
DNase-VIII-like
gpJ hypothetical proteins cor-like ltf gene (SIE)
LptD
BtuB
FritzSarasin
KarlJaspers
gpJ hypothetical proteins DNase-VIII-like
FritzHoffmann
ChristophMerian
VogelGryff
JeanPiccard
AugustePiccard
TheodorHerzl
Oekolampad
gpJ hypothetical proteins DNase-VIII-like
identity
Escherichia phage RTP
Escherichia phage AugustePiccard
rtp43 CDS
rtp44 CDS
rtp45 CDS
rtp46 CDS
hypothetical protein CDS
J-like tail tip protein CDS
hypothetical protein CDS
Dcm-like C5 cytosine methyltransferase
Escherichia phage AugustePiccard
Escherichia phage BrunoManser
Escherichia phage FritzSarasin
Escherichia phage IsaakIselin
Escherichia phage JeanPiccard
Escherichia phage JohannLBurckhardt
Escherichia phage JulesPiccard
Escherichia phage LeonhardEuler
Escherichia phage DanielBernoulli
Escherichia phage JakobBernoulli
Escherichia phage KarlJaspers
Escherichia phage PaulSarasin
Escherichia phage PeterMerian
lateral tail fiber protein CDS
DNA helicase CDS
recombination locus
lateral tail fiber locus
primase
DNA helicase locus

#### S1 Figure. Supplemental data for Figs 4 and 5.

**(A)** Maximum-Likelihood phylogeny of *Drexlerviridae* and the *Dhillonvirus* genus of *Siphoviridae* based on several core genes with bootstrap support of branches shown if > 70/100. It is clearly apparent that *Drexlerviridae* are split into two major clades, one formed by *Braunvirinae* and *Rogunavirinae* and another one formed by *Tempevirinae*, *Tunavirinae*, plus a few other groups. Given that the phylogenies strongly agree on all major branches, the root of the *Drexlerviridae* phylogeny shown in Fig 4B was placed between these two major clades. **(B)** The locus encoding lateral tail fibers was analyzed in a sequence alignment of the thirteen *Drexlerviridae* phage genomes of the BASEL collection (see *Materials and Methods*). It is clearly visible that the upstream and downstream regions (encoding genes involved in recombination as well as primase / helicase proteins for genome replication) are highly conserved and fully syntenic, with exception of small insertions in a few sequences. Conversely, only the most 5' end of the largest lateral tail fiber protein gene is very similar among all analyzed genomes (green circle), while the rest shows neither synteny nor clear homology across all genomes. **(C)** The *bona fide* RBP loci downstream of the *gpJ* homolog are shown for all small siphoviruses (*Drexlerviridae* and *Siphoviridae* of *Dhillonvirus*, *Nonagvirus*, and *Seuratvirus* genera) where we had experimentally determined the terminal receptor (together with selected representatives where previous work had determined the receptor specificity). **(D)** The *bona fide* RBP locus of *E. coli* phage RTP was aligned to the homologous locus of phage AugustePiccard (Bas01) as described in *Materials and Methods*. For the region comprising *rtp44* and *rtp45* of phage RTP, the pairwise identity of the two nucleotide sequences is ca. 93%.

### Figure S2

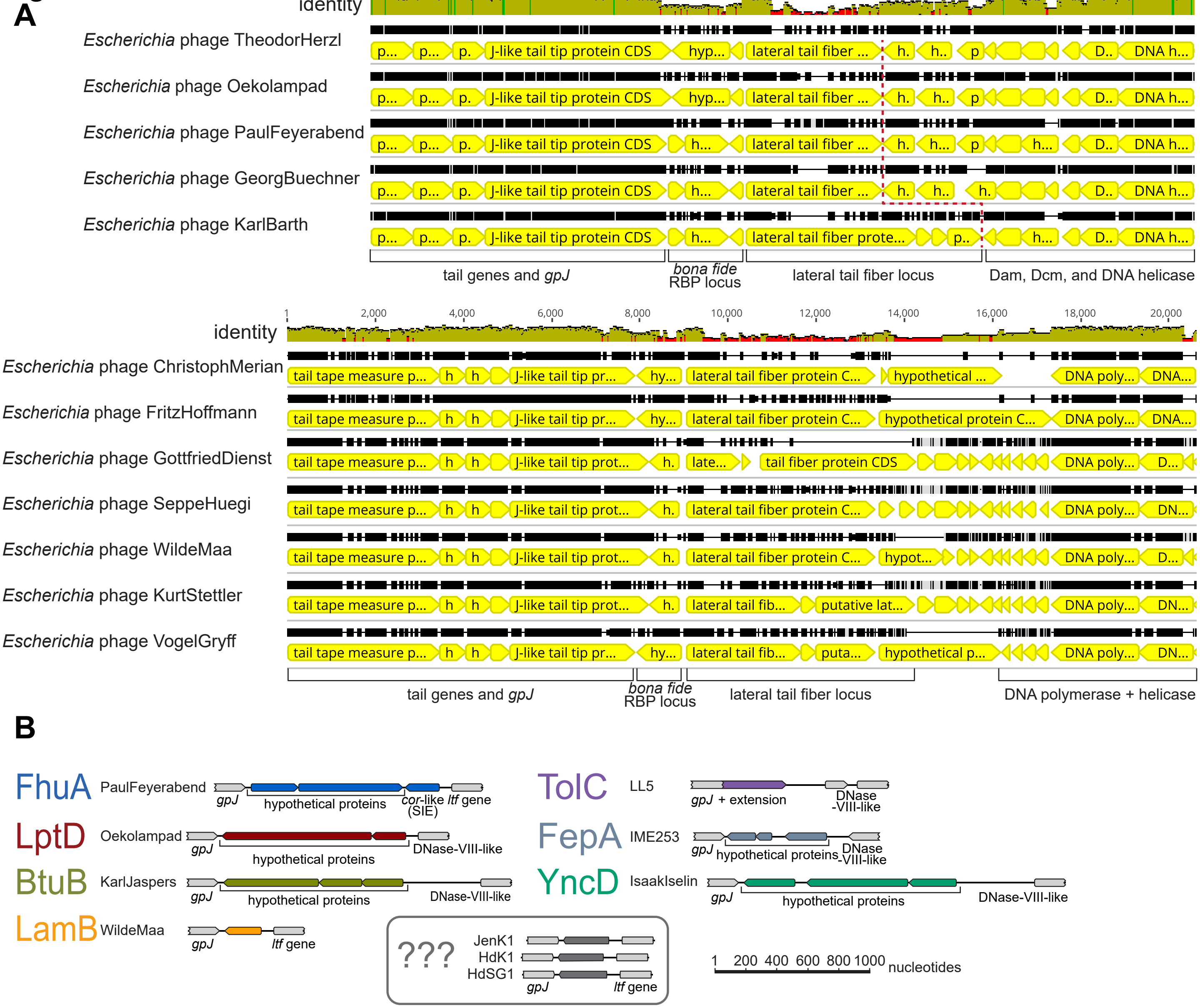
A
identity
Escherichia phage TheodorHerzl
Escherichia phage Oekolampad
Escherichia phage PaulFeyerabend
Escherichia phage GeorgBuechner
Escherichia phage KarlBarth
J-like tail tip protein CDS
lateral tail fiber ...
tail genes and gpJ
bona fide RBP locus
lateral tail fiber locus
Dam, Dcm, and DNA helicase
Escherichia phage ChristophMerian
Escherichia phage FritzHoffmann
Escherichia phage GottfriedDienst
Escherichia phage SeppeHuegi
Escherichia phage WildeMaa
Escherichia phage KurtStettler
Escherichia phage VogelGryff
tail tape measure p...
lateral tail fiber protein C...
tail fiber protein CDS
DNA polymerase + helicase
B
FhuA PaulFeyerabend
gpJ
hypothetical proteins
cor-like ltf gene (SIE)
LptD Oekolampad
DNase-VIII-like
BtuB KarlJaspers
LamB WildeMaa
ltf gene
TolC LL5
gpJ + extension
FepA IME253
YncD IsaakIselin
???
JenK1
HdK1
HdSG1
1 200 400 600 800 1000 nucleotides

#### S2 Figure. Supplemental data for Fig 6.

**(A)** The locus encoding lateral tail fibers was analyzed in sequence alignments of the five *Dhillonvirus* phage genomes of the BASEL collection (top) and the seven *Nonagvirus* + *Seuratvirus* phage genomes of the BASEL collection (bottom) as described in *Materials and Methods*. In both cases two clear dips in overall sequence similarity are obvious, once at the *bona fide* RBP locus and then at the lateral tail fiber locus downstream of the far 5' end of its first gene. **(B)** Schematic comparison of representative *bona fide* RBP loci as shown in S1C Fig to the corresponding allele of *E. coli* phages JenK1, HdK1, and HdsG1 that does not match clearly match any of them.

### Figure S3

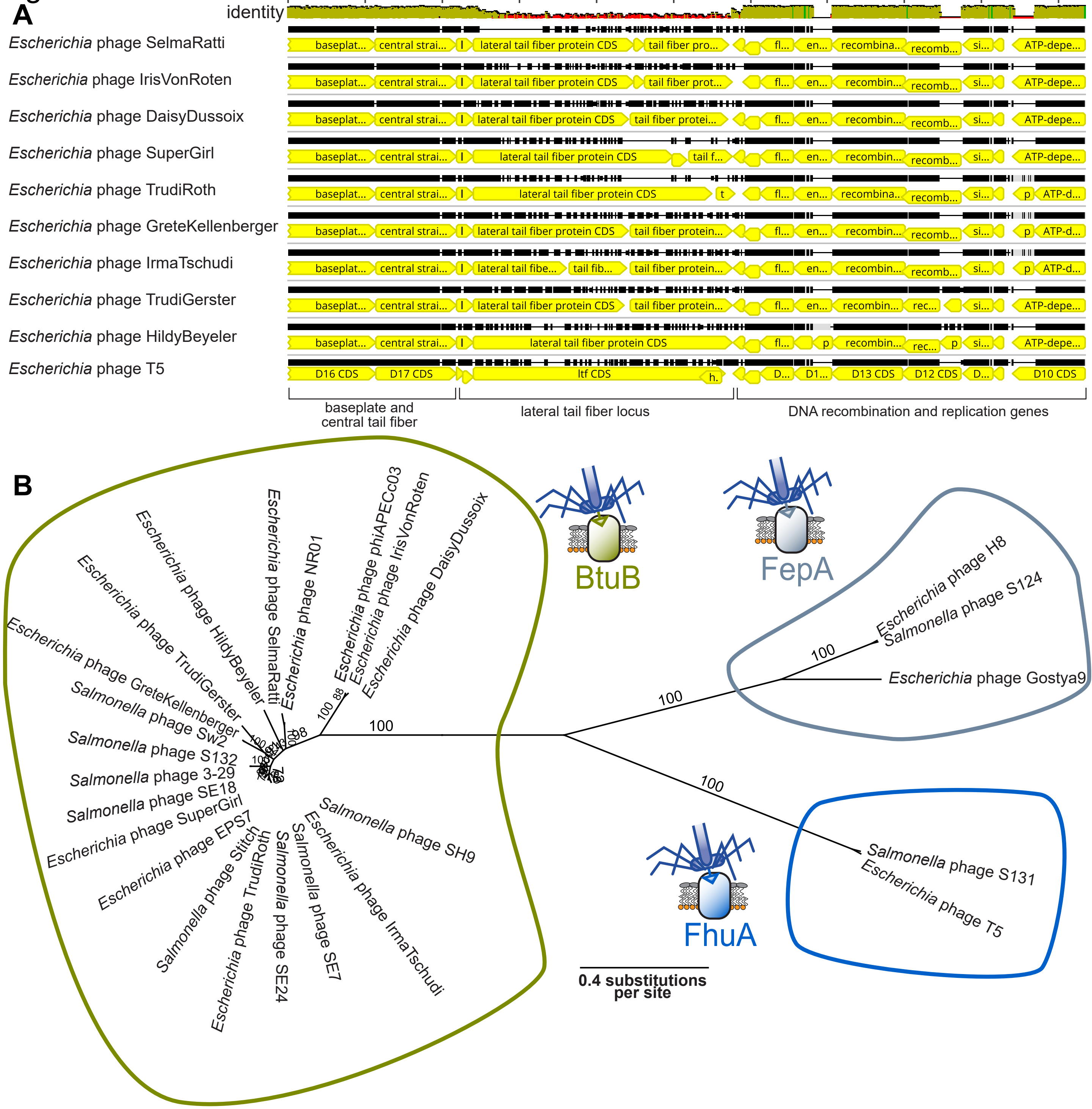

A
identity
Escherichia phage SelmaRatti
Escherichia phage IrisVonRoten
Escherichia phage DaisyDussoix
Escherichia phage SuperGirl
Escherichia phage TrudiRoth
Escherichia phage GreteKellenberger
Escherichia phage IrmaTschudi
Escherichia phage TrudiGerster
Escherichia phage HildyBeyeler
Escherichia phage T5
lateral tail fiber protein CDS
D16 CDS
D17 CDS
ltf CDS
D13 CDS
D12 CDS
D10 CDS
baseplate and central tail fiber
lateral tail fiber locus
DNA recombination and replication genes
B
Escherichia phage phiAPECc03
Escherichia phage NR01
Escherichia phage SelmaRatti
Escherichia phage HildyBeyeler
Escherichia phage TrudiGerster
Escherichia phage GreteKellenberger
Salmonella phage Sw2
Salmonella phage S132
Salmonella phage 3-29
Salmonella phage SE18
Escherichia phage SuperGirl
Escherichia phage EPS7
Salmonella phage Stitch
Escherichia phage TrudiRoth
Salmonella phage SE24
Salmonella phage SE7
Escherichia phage IrmaTschudi
Salmonella phage SH9
Escherichia phage IrisVonRoten
Escherichia phage DaisyDussoix
BtuB
FepA
FhuA
Escherichia phage H8
Salmonella phage S124
Escherichia phage Gostya9
Salmonella phage S131
Escherichia phage T5
100
0.4 substitutions per site

#### S3 Figure. Supplemental data for Fig 7.

**(A)** The locus encoding lateral tail fibers was analyzed in a sequence alignment of the *Demerecviridae: Markadamsvirinae* phage genomes of the BASEL collection as described in *Materials and Methods*. Sequence identity is high upstream and downstream of the lateral tail fiber locus (with exception of presence / absence of a few putative homing endonucleases) but drops considerably at the lateral tail fiber genes. Note that, as described previously, the lateral tail fibers can either be composed of a single large polypeptide or by two (or more) separate proteins [51, 60]. The same diversity in architecture of lateral tail fibers can also be seen at the corresponding loci of small siphoviruses (S1B and S2A Figs). **(B)** The illustration shows a phylogeny of the RBPs of all *Markadamsvirinae* phages shown in Fig 7B. Briefly, the RBP genes of all genomes (invariably encoded directly upstream of the terminase genes) were translated, aligned, and then used to generate a phylogeny as described in *Materials and Methods*. Three clear clusters emerge, one including all phages known to bind BtuB (left), one including all phages known to bind FepA (top right), and one including all phages known to bind FhuA (bottom right). Based on similar analyses by others [62], we conclude that the position of RBPs in these clusters is predictive of terminal receptor specificity of the phages encoding them.

### Figure S4

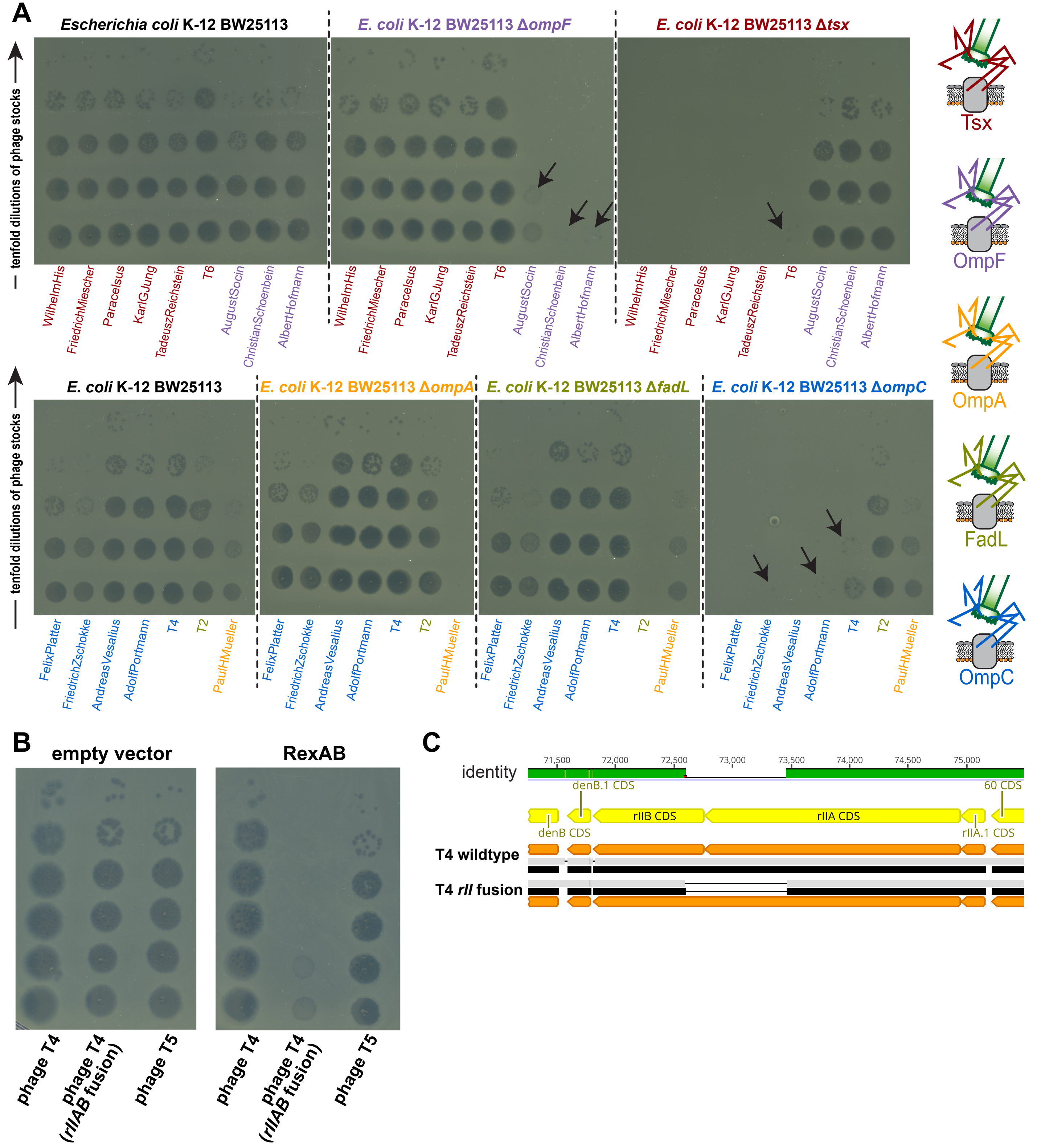
A
Escherichia coli K-12 BW25113
E. coli K-12 BW25113 ΔompF
E. coli K-12 BW25113 Δtsx
tenfold dilutions of phage stocks
WilhelmHis
FriedrichMiescher
Paracelsus
KarlGJung
TadeuszReichstein
T6
AugustSocin
ChristianSchoenbein
AlbertHofmann
Tsx
OmpF
OmpA
FadL
OmpC
E. coli K-12 BW25113
E. coli K-12 BW25113 ΔompA
E. coli K-12 BW25113 ΔfadL
E. coli K-12 BW25113 ΔompC
FelixPlatter
FriedrichZschokke
AndreasVesalius
AdolfPortmann
T4
T2
PaulHMueller
B
empty vector
RexAB
phage T4
phage T4 (rIIAB fusion)
phage T5
C
identity
71,500
72,000
72,500
73,000
73,500
74,000
74,500
75,000
denB.1 CDS
60 CDS
rIIB CDS
rIIA CDS
denB CDS
rIIA.1 CDS
T4 wildtype
T4 rII fusion

#### S4 Figure. Supplemental data for Fig 8.

**(A)** Top agar assays with different surface protein mutants of the KEIO collection in comparison to the ancestral *E. coli* K-12 BW25113 strain were performed with serial tenfold dilutions of all *Tevenvirinae* phages used in this study (undiluted high-titer stocks at the bottom and increasingly diluted samples towards the top). The phages show strongly or totally abolished growth on each one of the mutants which identifies the primary receptor of each phage (also indicated by the color code highlighted on the right). **(B)** Top agar assays of reference strain *E. coli* K-12 ΔRM carrying empty vector pBR322_ΔP*tet* or pAH213_rexAB were performed with serial tenfold dilutions of phage T4 wildtype, a T4 variant encoding an apparently hypomorphic *rIIAB* fusion (see (C)), and phage T5 (as control). The *rIIAB* mutant of phage T4 is unable to form plaques on the *rexAB*-expressing host and shows only "lysis from without" [98], while the T4 wildtype and phage T5 are not affected. **(D)** A T4 phage mutant was erroneously obtained from a culture collection instead of the wildtype and encoded a peculiar *rII* allele with fusion of the *rIIA* and *rIIB* open reading frames (as revealed by whole-genome sequencing). Since such a mutant seems unlikely to arise spontaneously during shipping, we find it likely that this phage strain is related to the *rIIAB* fusion mutants employed for discovery of the triplet nature of the genetic code that were once commonly used [concisely reviewed in reference 149, 150]. Notably, position and size of the deletion fusing *rIIA* and *rIIB* are indistinguishable from the sketch drawn by Benzer and Champe for the *rIIAB* fusion mutant *r1589* which was used in the aforementioned work [151]. Unlike T4 wildtype, the *rIIAB* fusion mutant is susceptible to *rexAB* when ectopically expressed from a plasmid vector (see (C)) and therefore validates functionality of the *rexAB* construct.

### Figure S5

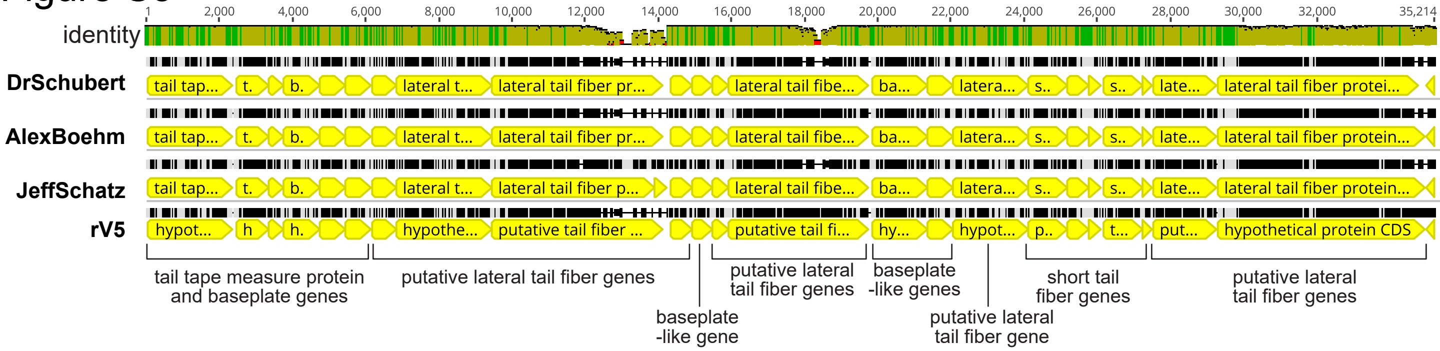
identity
1
2,000
4,000
6,000
8,000
10,000
12,000
14,000
16,000
18,000
20,000
22,000
24,000
26,000
28,000
30,000
32,000
35,214
DrSchubert
AlexBoehm
JeffSchatz
rV5
tail tape measure protein and baseplate genes
putative lateral tail fiber genes
baseplate -like gene
putative lateral tail fiber genes
baseplate -like genes
putative lateral tail fiber gene
short tail fiber genes
putative lateral tail fiber genes

#### S5 Figure. Supplemental data for Fig 9.

The illustration shows a sequence alignment of the locus encoding lateral tail fiber genes in phage rV5 and new isolates DrSchubert, AlexBoehm, and JeffSchatz that broadly cover the phylogenetic range of this genus (Fig 9B). It extends from the tail tape measure protein of phage rV5 (*gp49*) to the last large lateral tail fiber gene (*gp27*) [70, 71]. Note that most genes are highly conserved including the lateral tail fiber component with sugar deacetylase domain (compare Fig 9D; around position 8'000 in this alignment) or the short tail fiber locus (compare Fig 9E). Only the three largest lateral tail fiber genes show considerable allelic variation that is very strong for two of them (positions 10'000-14'000 and 16'000-19'000) and moderate for another one (positions 29'000 to 34'000).
